# A Small-molecule Antagonist Radiotracer for Positron Emission Tomography Imaging of the Mu Opioid Receptor

**DOI:** 10.1101/2024.10.12.618019

**Authors:** Konstantinos Plakas, Chia-Ju Hsieh, Dinahlee Saturnino Guarino, Catherine Hou, Wai-Kit Chia, Anthony Young, Alexander Schmitz, Yi-Pei Ho, Chi-Chang Weng, Hsiaoju Lee, Shihong Li, Thomas J.A. Graham, Robert H. Mach

## Abstract

The opioid crisis is a catastrophic health emergency catalyzed by the misuse of opioids that target and activate the mu opioid receptor. Traditional radioligands used to study the mu opioid receptor are often tightly regulated owing to their abuse and respiratory depression potential. In the present study, we sought to design and characterize a library of 24 non-agonist ligands for the mu opioid receptor. Ligands were evaluated for the binding affinity, intrinsic activity, and predicted blood-brain barrier permeability. Several ligands demonstrated single-digit nM binding affinity for the mu opioid receptor while also demonstrating selectivity over the delta and kappa opioid receptors. The antagonist behavior of **1A** and **3A** at the mu opioid receptor indicate that these ligands would likely not induce opioid-dependent respiratory depression. Therefore, these ligands can enable a safer means to interrogate the endogenous opioid system. Based on binding affinity, selectivity, and potential off-target binding, [^11^C]**1A** was prepared via metallophotoredox of the aryl-bromide functional group to [^11^C]methyl iodide. The nascent radiotracer demonstrated brain uptake in a rhesus macaque model and accumulation in the caudate and putamen. Naloxone was able to reduce [^11^C]**1A** binding, though the interactions were not as pronounced as naloxone’s ability to displace [^11^C]carfentanil. These results suggest that GSK1521498 and related congeners are amenable to radioligand design and can offer a safer way to query opioid neurobiology.

## Introduction

The mu opioid receptor (MOR) is one of the oldest drug targets known to humankind. Drugs targeting the MOR have been known since Homer’s Odyssey.^1^ Agonists for the MOR have been demonstrated to treat pain,^2^ cough suppression,^3^ and diarrhea.^4^ Antagonists for the MOR are equally useful, and have found use in the treatment of opioid overdose,^5^ alcohol-use disorder,^6^ and constipation.^7^ There is continued ardent interest in the development of next-generation MOR agonist analgesics to provide analgesia without risk of side-effects.^8,9^ Measuring the *in vivo* biodistribution of these drugs is paramount to predicting their safety and efficacy. One such technique capable of making real-time *in vivo* measurements about drug biodistribution and target engagement is positron emission tomography.

Positron Emission Tomography (PET) is an *in vivo* diagnostic imaging technique that utilizes a radiotracer to evaluate biological processes.^10^ PET imaging has been used to study several neurobiological processes, including neurodegenerative diseases,^11–13^ neuroinflammation,^14,15^ and addiction.^16,17^ PET is also used to determine target engagement between putative drugs and drug targets in vivo.^18,19^ Indeed, the interactions between the MOR and MOR radiotracer agonist [^11^C] carfentanil (CFN) have been well-characterized.^20,21^ [^11^C]CFN has been used to characterize the receptor engagement between the MOR and naloxone,^18^ and GSK1521498,^22^ both MOR antagonists.

An inherent hindrance to the deployment of [^11^C]CFN is its potent agonist nature that can lead to life-threatening respiratory depression, even at tracer doses.^23–26^ To circumvent this, there have been several reports detailing the radiosynthesis and evaluation of novel MOR PET radiotracers. However, these radiotracers are either agonists,^27^ or are non-selective (**Figure 1A**).^28–31^ Selective agonist radiotracers can still expose the patient to significant risk while the non-selective radiotracers can make interpreting of MOR function difficult to isolate. Therefore, there is an unmet need for the development of non-agonist (antagonist or inverse agonist), selective MOR PET tracers.

**Figure 1.**
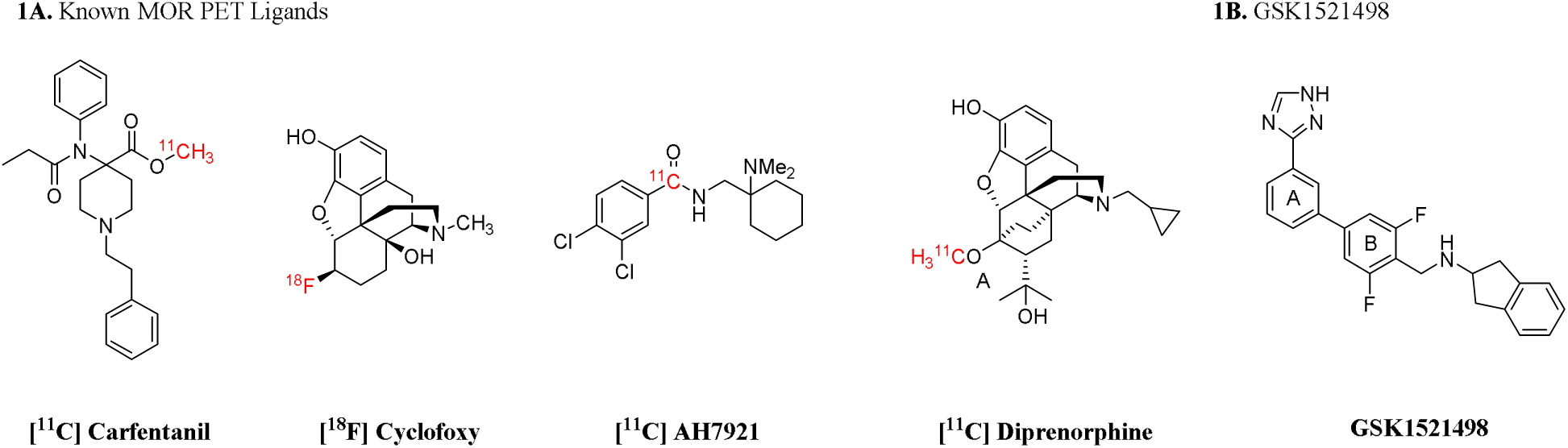
Several MOR PET tracers have been reported in the literature, including (A) potent agonist [^11^C] CFN, non-selective agonist [^18^F]cyclofoxy, agonist [^11^C]AH7921, and agonist [^11^C]diprenorphine. (B) GSK1521498 offers a new scaffold upon which to synthesize selective MOR antagonist radiotracers.

To ameliorate the lack of selective MOR non-agonists amenable to radiotracer design, we were inspired by the development of GSK1521498 (**Figure 1B**).^22,32–34^ GSK1521498 demonstrates a favorable safety profile and has been validated to be non-activating at the MOR.^34^ In addition, GSK1521498 has been shown to displace [^11^C]CFN from the brain in a human model, suggesting it can readily cross the blood-brain barrier (BBB), a key requirement for MOR PET tracer design.^22^

We synthesized a novel library of analogues structurally related to lead compound GSK1521498. The distinguishing feature of this library is installation of a methyl group to confer compatibility with metallophotoredox chemistry to enable radiomethyl installation.^35^ The nascent library was screened to determine *in vitro* binding affinity for the family of opioid receptors and potential off-target liabilities. The most potent and selective hits identified from radioligand competition bindings assays were further characterized using the PRESTO-Tango assay to determine their intrinsic activity. Selective non-agonist hits were radiolabeled with carbon-11 using late-stage radiomethylation.^35^ Subsequently, these radiotracers were evaluated for in a non-human primate (NHP) model for their ability to perform as non-agonist radiotracers for the MOR.

## Results

**Scheme 1:**
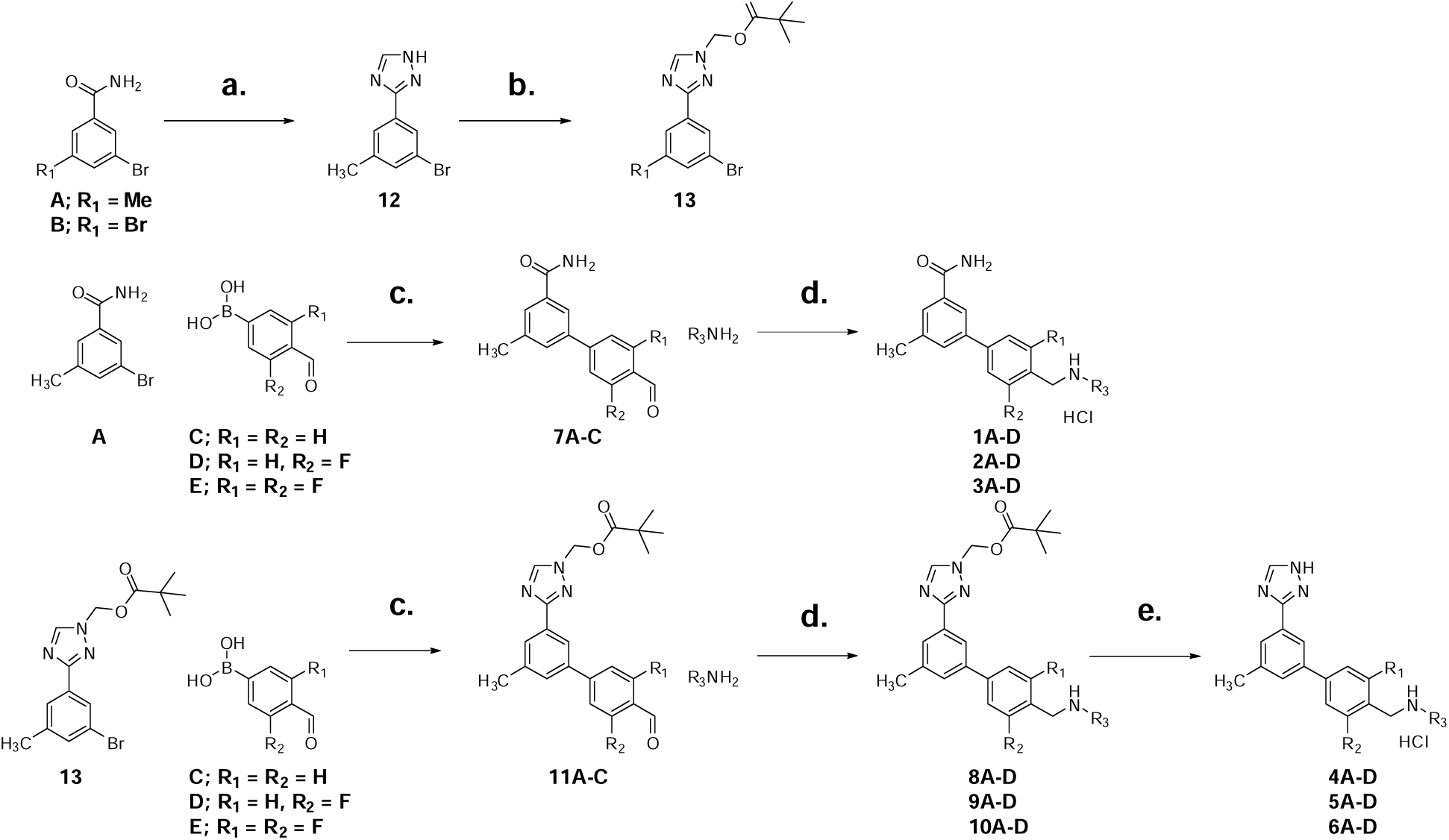
Synthesis of GSK1521498 analogue library evaluated in this study. **Conditions:** (a) 1. DMFDMA, 2. AcOH, hydrazine hydrate; (b) chloromethylpivalate, K_2_CO_3_, acetonitrile; (c) Pd(dppf)Cl_2_, NaHCO_3_, toluene, water, ethanol; (d) sodium cyanoborohydride, AcOH, 1:1 methanol:methylene chloride; (e) 1. sodium methoxide/methanol, ethanol, 2. 1.2 N HCl/ethanol.

### Chemistry

The MOR antagonist GSK152148 possesses favorable ligand properties; it is a potent and selective MOR non-agonist with a favorable pharmacodynamic profile as demonstrated by the drug’s advancement to phase II clinical trials. We therefore reasoned that structurally related starting material benzamide **A** would furnish a library of congeners that would possess similar physicochemical properties while being amenable to PET tracer design. The methyl at the 5-position of **A** promotes compatibility with the chosen radiolabeling methodology. The 5-position is also preferentially located in a position that should preclude inference from rotation of conjoined substituted phenyl rings, enabling a minimized energy conformation to be realized (**figure 2**).

**Figure 2:**
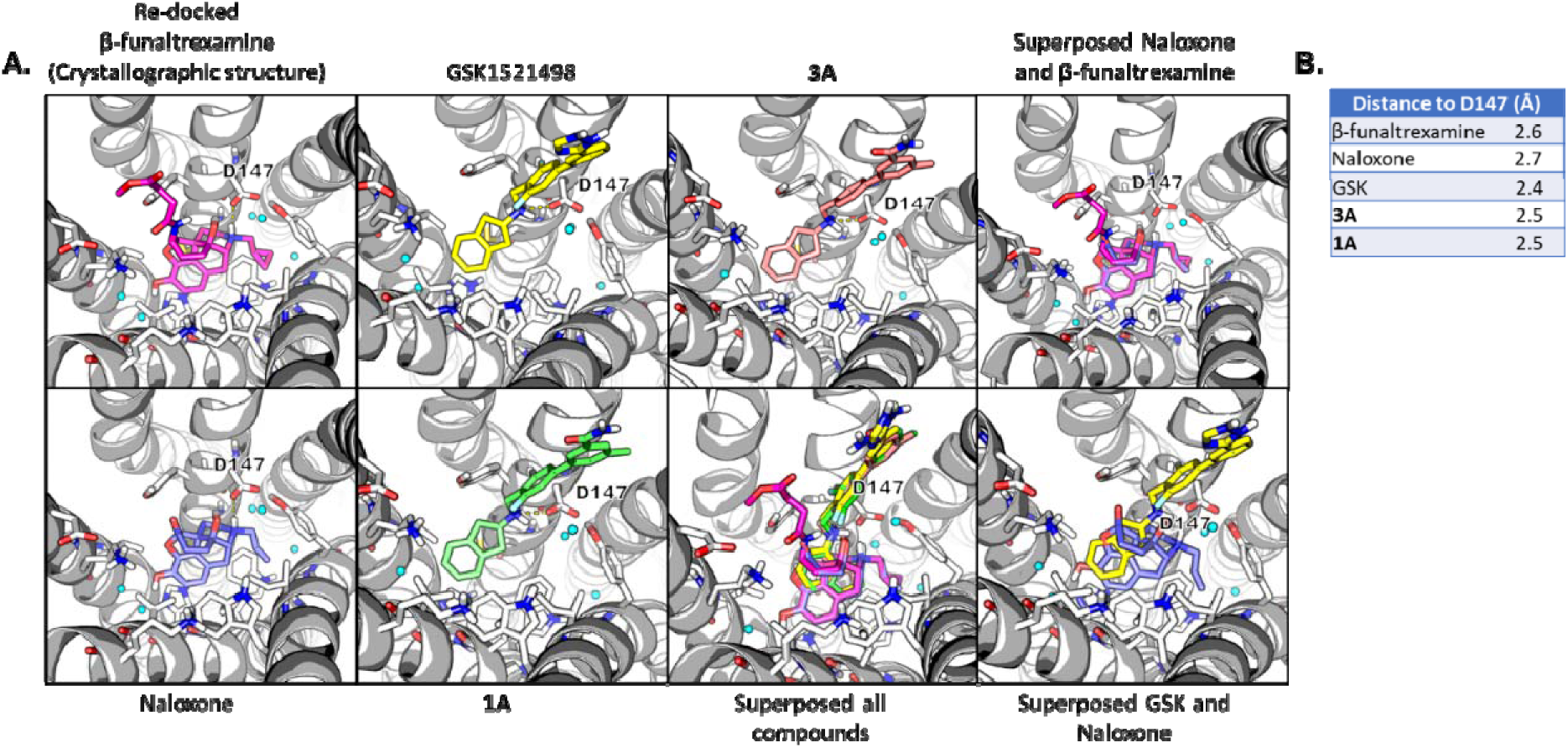
Docking was used to guide radioligand design. A) Lead ligand GSK1521498, putative radiotracers **1A** and **3A** and naloxone were iteratively docked to the mu opioid receptor (PDB: 4DKL). The 5’-position of ring A was chosen was a radiolabeling site with [^11^C]CH_3_I as the protein can accommodate the substitution. B) Opioids are alkaloids the contain a basic nitrogen. Key to opioid binding is the interaction of the basic nitrogen atom with aspartate residue D147.

Benzamide **A** was converted to the corresponding 1,2,4-triazole using previously reported methodology. The reaction proceeds through the formation of an intermediate N,N-acylformamidine before subsequent ring closure, yielding **12**.^36^ Triazole **12** was then reacted with pivaloyl chloride to form pivaloyl ester-protected triazole **13**. Intermediate **13** was reacted with commercially available formylboronic acid derivatives **C**, **D**, or **E** using Suzuki-Miyaura conditions to yield intermediates **11A**, **11B**, and **11C**. Carbon-nitrogen bond formation was carried out using a reductive amination between aldehyde containing compounds **11A**, **11B**, or **11C** and commercially available amines to furnish advanced intermediates **8A-D**, **9A-D**, and **10A-D**. At this point, the pivaloyl ester protecting groups were removed using a mixture of sodium methoxide and ethanol. Final compounds **4A-4D**, **5A-5D**, and **6A-6D** were formulated as HCl salts.

Benzamide **A** can also be directly coupled to commercially available formylboronic acid derivatives **C**, **D**, or **E** using Suzuki-Miyaura conditions to yield intermediates **7A-7C**. Carbon-nitrogen bond formation was carried out using a reductive amination between aldehyde containing compounds **7A**, **7B**, or **7C** and commercially available amines to yield final compounds **1A-1D**, **2A-2D**, and **3A-3D**, which were formulated as HCl salts. The synthesis of the triazole and benzamide containing compounds is presented in **scheme 1**.

### *In vitro* Radioligand Competition Binding Assays

A total of 24 compounds, **1A-D** ➔ **6A-6D** were evaluated for their binding affinity to the mu (MOR), delta (DOR), and kappa (KOR) opioid receptors (**table 1**) with select ligands also evaluated for potential off-target liabilities (**table S1**) at the Psychoactive Drug Screening Program (PDSP).^37^ Submitted ligands were initially screened at a single concentration of 10 μM and evaluated for their ability to displace a well-characterized tritiated ligand from a receptor of interest. Ligands showing a minimum of 50% inhibition at this concentration were further characterized in a 10-point radioligand competition binding assay. The 10-point radioligand competition binding assay evaluated compounds at 0.1, 0.3, 1, 3, 10, 30,100, 300 nM, 1, 3, 10 µM and was performed in triplicate. Nonlinear regression analysis yielded K_i_ values indicative ligand-receptor binding affinity. Binding affinity to the MOR, DOR, and KOR were used to evaluate ligands for their potential to function as MOR PET radiotracers, with ligands demonstrating potent binding to the MOR and selectivity over for DOR and KOR deemed as desirable. The results are summarized in **table 1**. In each series of compounds, the 2-aminoindane containing derivatives demonstrated the most potent binding affinity. These ligands were prioritized for off-target binding evaluation (**table S1)** intrinsic activity evaluation (**figure S2**).

**Table 1:** Binding Affinity and Selectivity of the Compounds Evaluated in This Study at the MOR, DOR, and KOR.

| Entry | MOR | DOR | KOR | Selectivity | Entry | MOR | DOR | KOR | Selectivity |
| --- | --- | --- | --- | --- | --- | --- | --- | --- | --- |
| **1A** | 8.3 | 410.3 | 308.3 | 1: 49.4: 37.1 | **4A** | 17.1 | 228.7 | 246.2 | 1: 13.3: 14.4 |
| **1B** | 339.2 | >10,000 | 86.1 | 1: NA: 0.2 | **4B** | 55.8 | 1006.2 | 26.9 | 1: 18:0: 0.4 |
| **1C** | 96.0 | 1420.6 | 123.2 | 1: 14.7: 1.2 | **4C** | 60.2 | 760.8 | 388.4 | 1: 12.6: 6.4 |
| **1D** | 181.3 | >10,000 | 876.0 | 1: NA 0.2 | **4D** | 11.3 | 145.7 | 73.2 | 1: 12.9: 2.8 |
| **2A** | 9.19 | 349.9 | 104.8 | 1: 38.0: 11.4 | **5A** | 13.7 | 171.4 | 67.3 | 1: 12.5: 4.9 |
| **2B** | 93.9 | >10,000 | 54.2 | 1: NA: 0.5 | **5B** | 124.4 | 885.5 | 94.4 | 1: 7.1: 0.7 |
| **2C** | 104.9 | >10,000 | 192.6 | 1: NA: 1.8 | **5C** | 44.5 | 314.9 | 100.3 | 1: 7.1: 2.3 |
| **2D** | 22.3 | 819.4 | 99.3 | 1: 36.6: 4.4 | **5D** | 12.1 | 108.6 | 33.0 | 1: 9.0: 2.7 |
| **3A** | 7.7 | 440.6 | 67.6 | 1: 56.7: 8.7 | **6A** | 4.20 | 98.0 | 19.4 | 1: 23.3: 4.6 |
| **3B** | 62.50 | >10,000 | 88.2 | 1: NA: 1.4 | **6B** | 267.8 | >10,000 | 137.7 | 1: NA: 0.5 |
| **3C** | 90.7 | 1884.0 | 89.9 | 1: 20.7: 0.9 | **6C** | 18.4 | 473.8 | 67.3 | 1: 25.7: 3.6 |
| **3D** | 15.2 | 660.8 | 54.6 | 1: 43.3: 3.5 | **6D** | 11.5 | 181.4 | 29.6 | 1: 15.8: 2.6 |

$K_i$ values of analogues **1A-1D** → **6A-6D** determined through radioligand competition binding experiments. Each value is the average of three experiments. Ligands were screened at the Psychoactive Drug Screening Program (PDSP). From the initial SAR, the -aminoindane containing analogues generally possessed the most potent binding at the MOR while demonstrating selectivity over the DOR and KOR. These ligands were prioritized for further study.

**Table 2:**
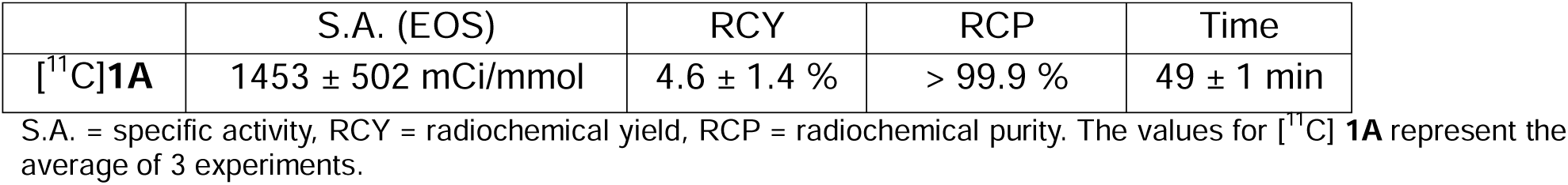
Radiochemical and Quality Control Parameters for [^11^C] 1A and [^11^C] 3A.

### Intrinsic Activity Evaluation through the PRESTO Tango Assay

As aforementioned, the archetypical clinical PET radiotracer for MOR imaging is [^11^C]CFN, which is a potent agonist. We sought to screen our library for intrinsic activity to limit our evaluation of PET tracers to non-agonists. The library was screened for intrinsic activity at the MOR, DOR, and KOR utilizing the PRESTO Tango assay developed at the PDSP.^38^ In agonist mode, ligands were evaluated for their ability to induce β-arrestin signaling as measured through an increase in luciferase-induced fluorescence. Candidate ligands were compared to known opioid receptor agonists. While known agonists demonstrated a marked increase in fluorescence indicative of ligand-induced signaling, ligands **1A** and **3A** did not exhibit an increase in fluorescence. Thus, the ligands developed here likely do not induce opioid signaling at the MOR, DOR, or KOR. Ligands were subsequently evaluated in antagonist mode. Here, ligands were preincubated with 1 μM of a known agonist. For example, MOR-expressing HEK cells were preincubated with 1 μM of DAMGO. Addition of a **1A** induced a decrease in luciferase-induced fluorescence, suggesting that **1A** competes for the same binding site as DAMGO. This suggests that **1A** exhibits non-agonist behavior. The same behavior was observed for **3A**. The results are summarized in **figure S2**.

### Predicted Blood-Brain Barrier (BBB) Permeability

The ligands containing the 2-aminoindane moiety (**1A** ➔ **6A**) were the most potent and selective MOR-binding ligands evaluated in this study. Ligands **1A** and **3A** were also devoid of MOR signaling, suggesting that they would be less likely to induce respiratory depression through MOR activation. To prioritize ligands for radiolabeling, -aminoindane containing derivatives were subjected to the BOILED-Egg plot^39^ analysis to predict BBB permeability. Methylated GSK1521498 congener **4A** exhibited an increase in lipophilicity such that it was not expected to be a viable candidate for BBB penetration. Isosteric replacement of the triazole moiety with a benzamide moiety reduced the lipophilicity in the order of **4A** ➔ **1A, 5A** ➔ **2A,** and **6A** ➔ **3A**. Furthermore, iterative fluorine atom replacement with hydrogen decreased lipophilicity in the order of **4A** ➔ **5A** ➔ **6A** and **1A*****2A** ➔ **3A**, as summarized in **figure 3**. Based on binding affinity, selectivity, intrinsic activity, and predicted BBB, permeability, **1A** was prioritized for radiolabeling.

### Radiochemistry

To enable the radiosynthesis of [^11^C]**1A** precursor **16** was prepared (**scheme 2**). Commercially available 3,5-dibromobenzamide **B** was coupled to commercially available formylphenylboronic acid **E** to yield intermediate **14.** A reductive amination was utilized to connect 2-aminoindane to **14** furnishing advanced intermediate **15**. Here, the secondary amine moiety weas protected using the fluorenylmethoxyloxycarbonyl (Fmoc) protecting group which can be easily removed with mild basic conditions, such as piperidine.

Radiotracer [^11^C]**1A** was prepared from precursors **16** as shown in **scheme 2**. In brief, [^11^C]CH_3_I was prepared using a Synthra MeI module. The nascent [^11^C]CH_3_I was coupled to the appropriate precursor using previously reported methods.^35^ This step took place in an integrated custom-made photoreactor, prepared by Synthra. The protecting group was removed using piperidine. The crude reaction mixtures were purified on semi-preparative HPLC and formulated in saline for injection. The radiochemical purity of [^11^C]**1A** was > 99% as confirmed by analytical HPLC (**figure S4, figure S5 and figure S6).** Relevant radiochemical characterization is shown in **table 3**.

**Scheme 2:**
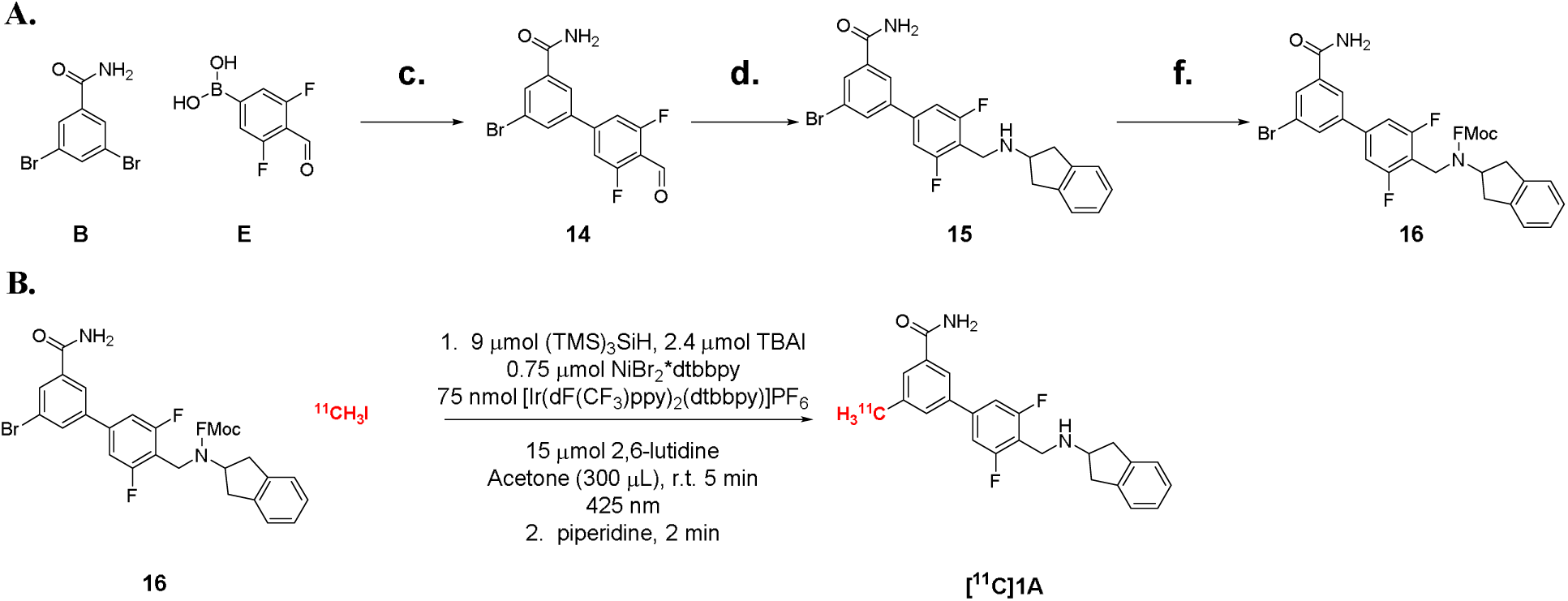
A) Synthesis of radiosynthetic precursors **16A** and **16B**. B) Radiosynthesis of [^11^C] **1A** and [^11^C] **3A** was enabled through metallophotoredox chemistry. **Conditions: (A)** (c) Pd(dppf)Cl2, NaHCO_3_, toluene, water, ethanol; (d) sodium cyanoborohydride, AcOH, 1:1 methanol:methylene chloride; (f) Fmoc-Chloride, sodium carbonate, 1:1 acetonitrile/water. **(B)** Radiolabeling of **16** was accomplished via organometallophotoredox chemistry.

### *In Vivo* Brain PET Analysis of [^11^C]1A in a Non-Human Primate (NHP) Model

Following the radiosynthesis of [^11^C]**1A**, the radiotracer was evaluated in a rhesus macaque model. The radiotracer demonstrated brain uptake and accumulation in the thalamus and putamen, areas of the brain known to express the mu opioid receptor.

Following baseline scan, a subsequent study was performed in the presence of naloxone, a known opioid antagonist that has been demonstrated to diminish [^11^C]CFN binding *in vivo* as shown in **figure 4**.

**Figure 4:**
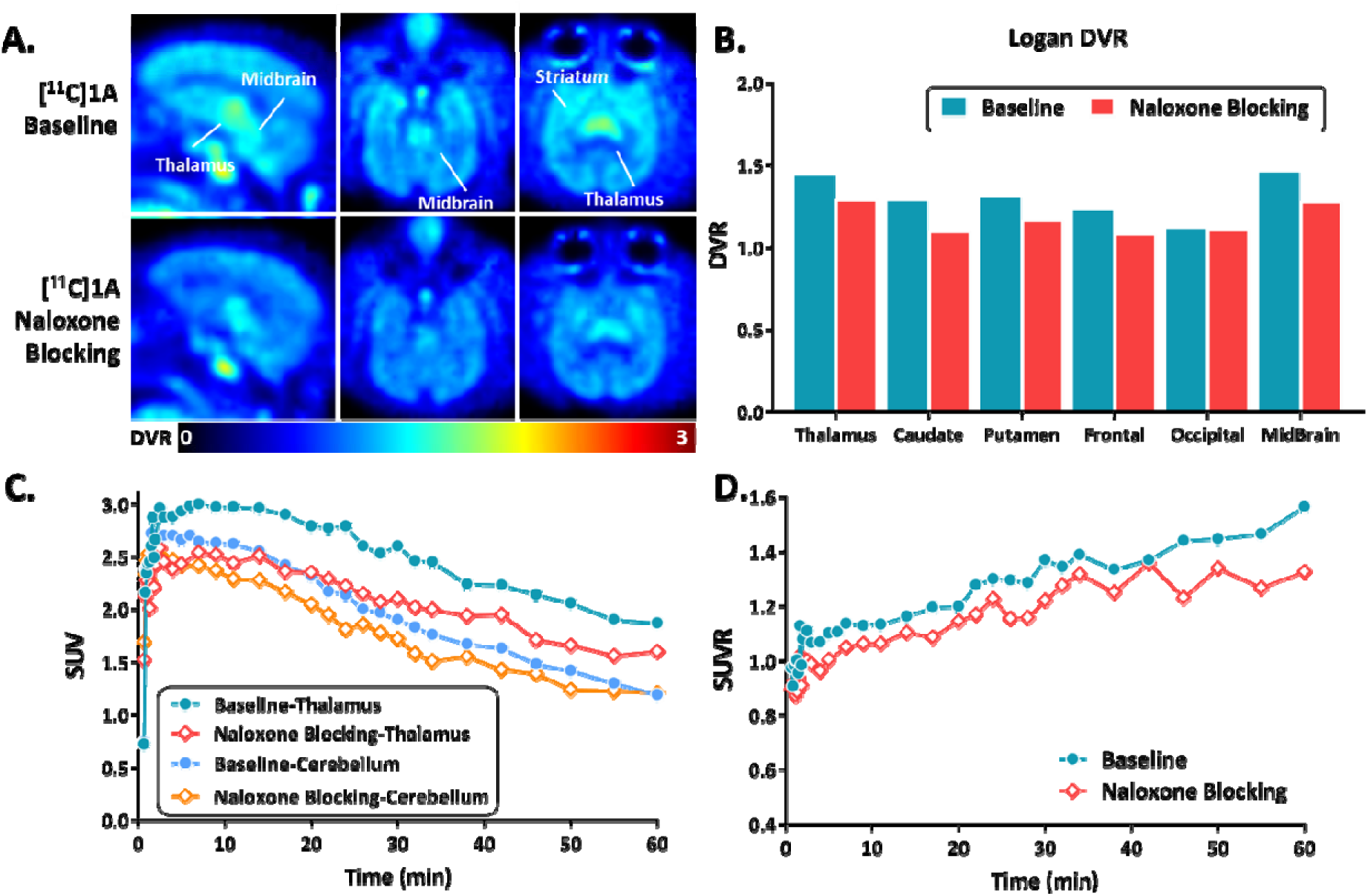
*In vivo* brain PET study of [^11^C]**1A** in a rhesus macaque model. (A) Brain uptake baseline images and blocking study performed with naloxone and [^11^C]**1A.** (B) Distribution volume ratio (DVR) of radiotracer uptake in th presence and asbsence of naloxone. Thalamus time-activity curves for radiotracer uptake in baseline and blocking study. (C) Time-activity curve of [^11^C]**1A** in the thalamus and cerebellum. (D) Distribution volume ratio (DVR) of radiotracer uptake for baseline and naloxone blocking studies.

## Discussion

GSK1521498 possesses a binding affinity towards the mu, delta, and kappa opioid receptors of 1.5 nM, 30.2 nM, and 20.4 nM respectively. We therefore reasoned that structurally similar chemical matter would exhibit favorable binding affinity to the mu opioid receptor while also displaying selectivity over the delta and kappa opioid receptors. The methyl group at the 5-position of aromatic ring A (**fig. 1**) seemed reasonable, as substitution would not induce any steric clashes (**fig. 2**). Various amines aromatic and aliphatic amines were selected from the patent literature and employed in synthesis to yield a diverse library of final compounds for screening. The compounds functionalized with the 2-aminoindane moiety possessed a more potent binding affinity than those containing the -4,4-geminaldimethyl, methylenecyclohexane, and -cyclohexyl derivatives, as shown in **table1**. Therefore the 2-aminoindane containing derivatives **1A** ➔ **6A** were prioritized for further study.

**4A** is the methylated analogue of GSK1521498. The binding affinity of **4A** towards the mu, delta and kappa opioid receptor was found to be 17.1 nM, 228.7 nM, and 246.1 nM respectively. The installation of the methyl group at the 5-position of ring A reduced the binding affinity of **4A** for the MOR, DOR, and KOR. Interestingly, the binding affinity of **4A** was improved by substituting the triazole with the isosteric benzamide function group, yielding **1A.** Triazoles and amides are known bioisoteres of one another.^40^ This trend was not uniform across the all -aminoindane containting derivatives tested, with **2A** demonstrating comparable MOR affinity as **5A** and **3A** having comparable MOR affinity as **6A**.

The use of fluorine to alter physiochemical properties of drug-like molecules has been well-documented.^41^ Here, fluorine atoms were iteratively removed from central aromatic ring, ring B. This led to a piecewise reduction in clogP as seen in **figure S3** in the order of **1A** ➔ **2A** ➔ **3A** and **4A** ➔ **5A** ➔ **6A.** Benzamide containing derivatives were less lipophilic than their triazole containing analogues in the order of **1A < 4A**, **2A < 5A**, and **3A < 6A**. Reduction in clogP is generally favorable many approved CNS drugs and imaging agents possess as clogP < 3.5.^42^ Based on the binding affinity and selectivity of the 24 ligands evaluated in this study, the 2-aminoindane containing derivatives were prioritized for off-target binding analysis as seen in **table S1.** Ligands **4A** and **5A** were excluded from further study as they were deemed not potent enough. Although ligand **6A** engaged more potently with the MOR, the binding selectivity over other opioid receptors was deemed to be problematic. Ligands **1A**, **2A**, and **3A** generally possessed favorable MOR binding, selectivity, and predicted BBB permeability. However, the off-target binding of **2A** with respect to the DAT and **3A** with respect to NET were concerns that precluded them from being radiolabeled. Therefore, ligand **1A** was advanced for intrinsic activity analysis.

The potent agonist activity of [^11^C]CFN requires stringent quality control analysis for (pre)clinical deployment. In this study, 2-aminoindane possessing ligand **1A** was evaluated for intrinsic activity at the PDSP using the PRESTO-Tango assay in cells expressing the mu opioid receptors. Agonist activation results in an increase in luminescence intensity resulting from β-arrestin-signaling-induced-luciferase transcription. This is best exemplified by known MOR agonist DAMGO as shown in **figure S2**. Ligands **1A** and **3A** do not exhibit a marked increase in luminescence. Therefore, it can be assumed that they will not behave as agonists. This is expected to limit the respiratory depression liability of these ligands. When the assay is run in antagonist mode, cells are preincubated with 1 μM of agonist compound. As non-agonist ligands such as naloxone (MOR non-agonist) are added in increasing quantities, they compete with the agonist for the binding site. This results in a decrease in luminescence. This behavior is exhibited by ligands **1A** and **3A** at the mu opioid receptors, suggesting that they behave as antagonists.

[^11^C]**1A** was radiosynthesized, validated, and evaluated for brain uptake in a rhesus macaque model. [^11^C]**1A** exhibited brain uptake with regional brain activity highest in the thalamus, caudate, and midbrain, areas of the rhesus macaque brain known to express the mu opioid receptor as shown in **figure 4**.

We next evaluated the ability of naloxone to block [^11^C]**1A** uptake. Naloxone is a known MOR antagonist and active pharmaceutical ingredient in Narcan, an opioid overdose rescue agent. Naloxone was pre-administered 10 minutes before radiotracer injection. Although naloxone demonstrated a significant reduction in [^11^C]CFN binding^43^, the ability of naloxone to affect [^11^C]**1A** binding was more subdued. Notably, GSK1521498 and related ligands (**1A** and **3A**) occupy different regions of the binding pocket than naloxone (**figure 2**). This may offer an explanation as to why [^11^C]**1A** is not displaced to the same extent as [^11^C]CFN binding by naloxone.^43^

## Conclusions

In summary, GSK1521498 and related congeners represent a rich area of chemical space for MOR PET radiotracer development. However, subsequent studies should prioritize the identification of more potent ligands for radiotracer development, either through structure-activity relationships or computer-assisted drug discovery. Key to the development of the ligands described above is their intrinsic activity; they are antagonists. This offers a safer way to study the neurobiology of opioid pharmacology. Ligand binding affinity were assessed using radioligand competition binding assays. Suitable candidates were selected based on adequate binding affinity, selectivity, and intrinsic activity. Radiolabeled ligand [^11^C]**1A** demonstrated uptake in the MOR rich areas of the brain including the caudate and putamen. Naloxone diminished binding, though not to the same extent as it diminishes [^11^C]carfentanil binding. Furthermore, these ligands may be of therapeutic value in the development of opioid overdose rescue agents.

## Experimental

### General

Commercial reagents and solvents were purchased from reputable sources, including Sigma-Aldrich (MO, U.S.A.), Combi Blocks (CA, U.S.A.), Enamine (Kyiv, UA), or AstaTech (PA, U.S.A.), and used as received. Thin layer chromatography (TLC) was performed using TLC silica gel 60W F254S plates for monitoring reactions, and the spots were visualized under UV light (254 nm). Flash chromatography purifications were conducted on a Biotage Isolera One system (Uppsala, Sweden). ^1^ H and ^13^C NMR spectra were recorded on a Bruker NEO-400 spectrometer (Bruker, MA, U.S.A.) with chemical shifts (δ) reported in parts per million (ppm) relative to the deuterated solvent as an internal reference. Mass spectra (m/z) were acquired on a 2695 Alliance LC-MS (Waters Corporation, MA, U.S.A.) with positive electrospray ionization (ESI+), and high-resolution mass spectra (HRMS, m/z) were obtained using a waters LCT premier mass spectrometer (Waters Corporation, MA, U.S.A.). The purity of investigated analogs was determined prior to the screening and confirmed to be >95% by UHPLC-MS.

## General Methods

### General Procedure A

Triazole formation for the synthesis of **12**. A 15-mL round bottom flask was charged with 3-bromo-5-methylbenzamide (4.67 mmol, 1.0 eq) and N,N-Dimethylformamide dimethyl acetal (2.50 mL). This mixture heated to 80 °C for 1 hr and was subsequently cooled to ambient temperature. The mixture was poured into crushed ice (30 mL) and stirred for 1 hr. The precipitated solid was collected by filtration and dried. The intermediate N-acyl formimadine was dissolved in acetic acid (6.15 mL) and hydrazine monohydrate (4.67 mmol, 1.0 eq) was added. This heated to 90 °C for 1 hr. The reaction was cooled, and dumped into crushed ice (30 mL) and stirred for 1 hr. This mixture was quenched with saturated sodium carbonate and the product was extracted with ethyl acetate (3 x 20 mL). The combined organic extracts were dried over sodium sulfate, filtered, and concentrated. The resulting white solid was used without further purification.

### General Procedure B

Pivalate-ester protection of triazole **12** yielding **13**. A 15-mL round bottom flask was charged with **12** (16.50 mmol, 1.0 eq), potassium carbonate (24.76 mmol, 1.50 eq) and anhydrous acetonitrile (20 mL). Subsequently, chloromethyl pivalate (24.76 mmol, 1.50 eq) was added. This mixture heated to 80 °C for 1.5 hr and was subsequently cooled to ambient temperature. The mixture was poured into crushed ice (30 mL) and stirred for 1 hr. The product was extracted with methylene chloride (3 x 15 mL). The combined organic extracts were dried over sodium sulfate, filtered, and concentrated. The resulting residue was purified on silica with a gradient 10% EtOAc/hexanes to 100% EtOAc to yield a white solid.

### General Procedure C

Suzuki-Miyaura Cross Coupling Yielding **7A-C** and **11A-C**. A 50-mL round-bottom flask was charged with 3-bromo-5-methylbenzamide (2.50 mmol, 1.0 eq), a commercially available formylphenyl boronic acid (2.75 mmol, 1.1 eq), and sodium bicarbonate (7.50 mmol, 3.0 eq). This was dissolved in a mixture of toluene, ethanol, and water (9.0 mL, 4.5 mL, and 5.0 mL). This mixture was evacuated and backfilled with nitrogen three times. [1,1′-Bis(diphenylphosphino)ferrocene]dichloropalladium(II) (36.6 mg, 2.0 mol %) was added and the resulting reaction mixture was heated to 80 °C for 18 hr. The reaction mixture was cooled to ambient temperature and quenched with water. The product was extracted with ethyl aceate (3 x 15 m) and the organic layer was dried over sodium sulfate, filtered and concentrated. The resulting solid was triturated with ether to yield a yellow solid.

### General Procedure D

Reductive Amination Yielding **1A-D**, **2A-D**, **3A-D**, **8A-D**, **9A-D**, and **10A-D**. A 20-mL vial was charged with 7**A-C** or **11A-11C** (0.800 mmol, 1.0 eq) and a commercially available amine (1.20 mmol, 1.5 eq). This was dissolved in 1:1 methylene chloride and methanol (9.60 mL). Acetic acid (480 μL) was added, and the resulting mixture was stirred at ambient for 1 hr. Subsequently, sodium cyanoborohydride (2.40 mmol, 3.0 eq) was added, and the mixture was stirred for 18 hr at ambient temperature. The mixture was concentrated, redissolved in methylene chloride, and washed with saturated brine. The organic layer was dried with sodium sulfate, filtered, and concentrated. The residue was purified on silica with a gradient CH_2_Cl_2_➔ 10% MeOH/CH_2_Cl_2_. Product containing fractions were combined and concentrated to yield a clear residue.

### General Procedure E

Pivalate Ester Deprotection. A 20-mL vial was charged with **8A-10D** (0.441 mmol, 1.0 eq). This was dissolved in ethanol (3.40 mL), and a mixture of sodium methoxide in methanol (25 wt% in methanol, 0.882 mmol, 2.0 eq) was added. This stirred at ambient for 1 hr. Subsequently, a 1.2 N solution of hydrochloric acid in ethanol (3.75 mmol, 8.5 eq) was added and the reaction heated to 80 °C for 1 hr. This was cooled to ambient temperature and quenched with saturated sodium bicarbonate (aq). The organic product was extracted with ethyl acetate (3 x 15 mL), dried over sodium sulfate, filtered, and concentrated. The residue was purified on silica with a gradient CH_2_Cl_2_➔ 10% MeOH/CH_2_Cl_2_. Product containing fractions were combined and concentrated to yield a clear residue. The residue was dissolved in 1:1 methanol and methylene chloride. To this solution was added HCl in ether (2N, 1.0 eq) and the mixture was left to stand. The crystallized material was collected to yield a white crystalline solid.

### General Procedure F

Fmoc Carbamate Protection. A 5-mL round bottom flask was charged with 15 (0.155 mmol, 1.0 eq), sodium carbonate (0.310 mmol, 2.0 eq) and a 1:1 mixture of acetonitrile:water (1.50 mL). Subsequently, Fmoc-Cl (0.186 mmol, 1.20 eq) was added and the reaction mixture stirred at ambient for 1.5 hr. The reaction mixture was diluted with water (5 mL) and the product was extracted with methylene chloride (3 x 3 mL). The organic layer was dried over sodium sulfate, filtered, and concentrated. The crude residue was purified on silica with a gradient CH_2_Cl_2_ ➔ 10% MeOH/CH_2_Cl_2_. Product containing fractions were combined and concentrated to yield a white crystalline solid.

**1A**: R_1_ = R_2_ = F, R_3_ = 2-aminoindane 35% white solid ^1^H-NMR (DMSO) 9.66 (br s, 2H), 8.14 (s, 1H), 8.08 (s, 1H), 7.79 (d, 2H, J = 9.7 Hz), 7.71 (d, 2H, J = 9.0 Hz), 7.45 (s, 1H), 7.30-7.28 (m, 2H), 7.23-7.20 (m, 2H), 4.27 (s, 2H), 4.11 (q, 1H, J = 7.4 Hz), 3.40-3.36 (m, 3H), 3.20 (dd, 2H, J = 16.1 Hz, 7.7 Hz, ), 2.44 (s, 3H). ^13^C-NMR (DMSO) 168.0, 163.3 (d, J = 8.3 Hz), 160.8 (d, J = 8.4 Hz), 140.1, 139.2, 137.3, 135.5, 130.6, 129.4, 127.4, 125.0, 123.4, 110.5, 110.2, 58.20, 37.1, 36.5, 21.4. HRMS (ESI) m/z [M + H] ^+^: 393.1773 (calculated), 393.1776 (found).

**1B**: R_1_ = R_2_ = F, R_3_ = cyclohexane 31% white solid:^1^H-NMR (DMSO) δ 8.07 (s, 1H), 8.01 (s, 1H), 7.74 (d, 2H, J = 5.3 Hz), 7.53 (d, 2H, J = 8.8 Hz), 7.42 (s, 1H), 3.85 (s, 2H), 2.42 (s, 3H), 1.87 (d, 2H, J = 11.2 Hz), 1.70-1.67 (m, 3H), 1.56-1.54 (m, 1H), 1.24-1.02 (m, 6H). ^13^C-NMR (DMSO) d 168.1, 163.2 (d, J = 9.7 Hz), 160.8 (d, J = 9.7 Hz), 139.0, 137.8, 135.4, 130.5, 128.9, 123.2, 110.2, 109.9, 55.8, 37.3, 32.8, 26.2, 24.7. 21.4. HRMS (ESI) m/z [M + H] ^+^: 359.1930 (calculated), 359.1937 (found).

**1C**: R_1_ = R_2_ = F, R_3_ = methylenecyclohexane 26% white solid: ^1^H-NMR (DMSO) δ 9.36 (s, 2H), 8.16 (s, 1H), 8.08 (s, 1H), 7.79 (d, 2H, J = 6.5 Hz), 7.69 (d, 2H, J = 9.0 Hz), 7.45 (s, 1H), 4.21 (s, 2H), 2.84 (d, 2H, J = 6.5 Hz), 2.43 (s, 3H), 1.83-1.76 (m, 3H), 1.70-1.68 (m, 2H), 1.63-1.61 (m, 1H), 1.25-1.09 (m, 4H), 0.99-0.91 (m, 2H). ^13^C-NMR (DMSO) δ 168.0, 163.3 (d, J = 8.1 Hz), 160.9 (d, J = 8.1 Hz), 144.3, 139.2, 137.3, 135.5, 130.6, 129.4, 123.4, 110.5, 110.2, 107.5, 53.2, 38.3, 34.5, 30.5, 26.0, 25.5, 21.41. HRMS (ESI) m/z [M + H] ^+^: 373.2086 (calculated), 373.2089 (found).

**1D**: R_1_ = R_2_ = F, R_3_ = 4,4-dimethylcyclohexane 33% white solid: ^1^H-NMR (DMSO) δ 9.48 (s, 2H), 8.17 (s, 1H), 8.09 (s, 1H), 7.79 (d, 2H, J = 8.4 Hz), 7.70 (d, 2H, J = 9.0 Hz), 7.44 (s, 1H), 4.22 (s, 1H), 3.05 (t, 1H, J = 10.9 Hz), 2.43 (s 3H), 1.98 (d, 2H, J = 9.8 Hz), 1.71 (q, 2H, J = 11.5 Hz), 1.45 (d, 2H, J = 13.1 Hz), 1.22 (q, 2H, J = 7.9 Hz), 0.93 (s, 3H), 0.91 (s, 3H). ^13^C-NMR (DMSO) δ 168.0, 163.3 (d, J = 8.3 Hz), 160.8 (d, J = 8.4 Hz), 144.2, 139.2, 137.3, 135.5, 130.6, 129.4, 123.4, 110.5, 57.4, 37.0, 35.2, 32.2, 29.8, 24.3, 21.41. HRMS (ESI) m/z [M + H] ^+^: 387.2243 (calculated), 387.2242 (found).

**2A**: R_1_ = H, R_2_ = F, R_3_ = 2-aminoindane 37% white solid: ^1^H-NMR (DMSO) δ 8.05 (s, 1H), 7.98 (s, 1H), 7.70 (d, 2H, J = 5.3 Hz), 7.57-7.54 (m, 3H), 7.38 (s, 1H), 7.20-7.17 (m, 2H), 7.11-7.09 (m, 2H), 3.84 (s, 2H), 3.55 (quin, 1H, J = 6.7 Hz), 3.18 (d, 1H, J = 4.9 Hz), 3.09 (dd, 2H, J = 7.0 Hz, J = 8.6 Hz), 2.74 (dd, 2H, J = 6.3 Hz, J = 9.3 Hz), 2.42 (s, 3H). ^13^C-NMR (DMSO) δ 168.1, 162.8, 160.3, 143.4, 140.0, 139.0, 138.4, 135.5, 133.5, 130.6, 128.7, 125.0, 123.4, 123.3, 118.9, 114.2, 114.0, 57.9, 42.4, 36.1, 21.5. HRMS (ESI) *m/z* [M + H] ^+^: 375.1873 (calculated), 375.1888 (found).

**2B**: (161B) R_1_ = H, R_2_ = H, R_3_ = cyclohexane 28% white solid: ^1^H-NMR (DMSO) δ 8.06 (s, 1H), 7.99 (s, 1H), 7.70 (d, 2H, J = 6.3 Hz), 7.61-7.55 (m, 3H), 7.39 (s, 1H), 3.86 (s, 2H), 2.42 (s, 3H), 1.90 (d, 2H, J = 12.0 Hz), 1.71-1.67 (m, 2H), 1.57-1.55 (1H), 1.25-1.06 (m, 6H). ^13^C-NMR (DMSO) δ 168.1, 162.8, 160.3, 143.3, 139.0, 138.4, 135.5, 133.5, 130.6, 128.7, 123.4, 123.2, 119.1, 119.0, 114.2, 113.9, 56.9, 28.9, 25.2, 24.5, 21.5. HRMS (ESI) *m/z* [M + H] ^+^: 341.2029 (calculated), 341.2029. (found).

**2C:** R_1_ = H, R_2_ = F, R_3_ = methylenecyclohexane 35% white solid: ^1^H-NMR (DMSO) δ 8.06 (s, 1H), 7.99 (s, 1H), 7.70 (d, 2H, J=5.4 Hz), 7.58-7.55 (m, 3H), 7.39 (s, 1H), 3.82 (s, 2H), 3.18 (s, 1H), 2.42 (s, 3H), 1.76 (d, 2H, J = 13.3 Hz), 1.69-1.65 (m, 3H), 1.50-1.41 (m, 1H), 1.25-1.10 (m, 3H), 0.93-0.83 (m, 2H). ^13^C-NMR (DMSO) δ 168.1, 162.8, 160.4, 143.3, 139.0, 138.4, 135.5, 130.6, 128.7, 123.4, 123.2, 118.6, 118.5, 114.1, 113.9, 53.0, 43.7, 34.7, 30.6, 26.0, 25.5, 21.5. HRMS (ESI) *m/z* [M + H] ^+^: 355.2186 (calculated), 355.2182 (found).

**2D:** R_1_ = H, R_2_ = F, R_3_ = 4,4-dimethylcyclohexane 24% white solid: ^1^H-NMR (DMSO) δ 8.06 (s, 1H), 7.98 (s, 1H), 7.70 (d, 2H, J = 5.9 Hz), 7.61-7.55 (m, 3H), 7.39 (s, 1H), 3.85 (s, 2H), 3.18 (s, 1H), 2.55 (s, 1H), 2.42 (s, 3H), 1.75-1.71 (m, 2H), 1.39-1.25 (m, 4H), 1.18-1.11 (m, 2H), 0.89 (s, 3H), 0.88 (s, 3H). ^13^C-NMR (DMSO) δ 168.1, 162.7, 160.4, 143.3, 139.0, 138.41, 135.5, 133.5, 130.6, 128.7, 123.4, 123.2, 114.2, 113.9, 57.0, 32.3, 29.7, 24.6, 24.2, 21.5. HRMS (ESI) *m/z* [M + H] ^+^: 369.2342 (calculated), 369.2337 (found).

**3A**: R_1_ = R_2_ = H, R_3_ = 2-aminoindane 22% white solid: ^1^H-NMR (DMSO) δ 8.04 (s, 1H), 7.95 (s, 1H), 7.70-7.76 (m, 3H), 7.64 (s, 1H), 7.48 (d, 2H, J = 8.2 Hz), 7.36 (s, 1H), 7.20-7.18 (m, 2H), 7.12-7.10 (m, 2H), 3.84 (s, 2H), 3.56 (q, 1H, J = 6.8 Hz), 3.09 (dd, 2H, J = 8.5 Hz, 7.1 Hz), 2.77 (dd, 2H, J = 9.1 Hz, 6.5 Hz), 2.42 (s, 3H). 13C-NMR (DMSO) d 168.4, 142.3, 140.4, 138.6, 138.5, 135.4, 130.3, 129.2, 127.5, 127.0, 126.6, 124.9, 123.3, 59.0, 51.4, 21.5. HRMS (ESI) m/z [M + H] ^+^: 357.1967 (calculated), 357.1978 (found).

**3B:** R_1_ = R_2_ = H, R_3_ = cyclohexane 30% white solid: ^1^H-NMR (DMSO) δ 8.04 (s, 1H), 7.95 (s, 1H), 7.68-7.77 (m, 3H), 7.64 (s, 1H), 7.46 (d, 2H, J = 8.2 Hz), 7.36 (s, 1H), 3.83 (s, 2H), 2.42 (s, 3H), 1.90 (d, 2H, J = 11.4 Hz), 1.71-1.67 (m, 2H), 1.56-1.55 (m, 1H), 1.24-1.11 (m, 6H). ^13^C-NMR (DMSO) δ 168.39, 140.39, 138.65, 135.37, 130.33, 129.18, 127.56, 127.01, 123.25, 55.83, 49.63, 32.76, 26.23, 24.85, 21.51. HRMS (ESI) m/z [M + H] ^+^: 323.2124 (calculated), 326.2123 (found).

**3C**: R_1_ = R_2_ = H, R_3_ = methylenecyclohexane 32%: ^1^H-NMR (DMSO) δ 8.04 (s, 1H), 7.96 (s, 1H), 7.68-7.67 (m, 3H), 7.64 (s, 1H), 7.46 (d, 2H, J = 8.18 Hz), 7.36 (s, 1H), 3.80 (s, 2H), 2.55 (s, 1H), 2.42 (s, 3H), 2.40 (s, 1H), 1.77 (d, 2H, J = 12.91 Hz), 1.68-1.61 (m, 23), 1.49-1.42 (m, 1H), 1.22-1.11 (m, 3H), 0.90-0.86 (m, 2H). ^13^C-NMR (DMSO) δ 168.39, 140.37, 138.64, 135.37, 130.33, 129.22, 127.58, 127.02, 123.27, 55.48, 52.86, 40.92, 37.55, 31.43, 26.68, 26.03, 21.53. HRMS (ESI) m/z [M + H] ^+^: 337.2275 (calculated), 337.2277 (found).

**3D**: R_1_ = R_2_ = H, R_3_ = 4,4-dimethylcyclohexane 35% white solid: ^1^H-NMR (DMSO) δ 8.04 (s, 1H), 7.95 (s, 1H), 7.68 (d,3H, J = 8.02 Hz), 7.64 (s, 1H), 7.47 (d, 2H, J = 8.13 Hz), 7.36 (s, 1H), 3.85 (s, 2H), 2.55 (s, 1H), 2.42 (s, 3H), 1.76-1.72 (m, 2H), 1.39-1.27 (m, 4H), 1.17-1.11 (m, 2H), 0.89 (s, 3H), 0.88 (s, 3H). ^13^C-NMR (DMSO) 168.38, 140.36, 138.65, 135.38, 130.33, 129.26, 127.57, 127.03, 123.26, 55.93, 49.65, 37.58, 32.11, 30.21, 28.26, 21.51. HRMS (ESI) m/z [M + H] ^+^: 351.2431 (calculated), 351.2424 (found).

**4A**: R_1_ = R_2_ = F, R_3_ = 2-aminoindane 66% white solid: ^1^H-NMR (DMSO) δ 8.45 (br s, 1H), 8.12 (s, 1H), 7.89 (s, 1H), 7.66 (s, 1H), 7.49 (d, 2H, J = 8.23 Hz), 7.19-7.16 (m, 2H), 7.10-7.09 (m. 2H), 3.85 (s, 2H), 3.53 (t, 1H, J = 6.5 Hz), 3.09 (dd, 2H, J = 8.7, 7.0 Hz), 2.74 (dd, 2H, J = 9.6, 6.1 Hz), 2.46 (s, 3H). ^13^C-NMR (DMSO) 163.3 (d, J = 8.3 Hz), 160.8 (d, J = 8.3 Hz), 140.0, 139.7, 138.0, 128.9, 127.5, 125.0, 122.0, 110.5, 110.2, 65.4, 58.2, 55.4, 49.1, 37.1, 36.4, 21.48, 15.6. HRMS (ESI) m/z [M + H] ^+^: 417.1886 (calculated), 417.1889 (found).

**4B**: R_1_ = R_2_ = F, R_3_ = cyclohexane 68% white solid: ^1^H-NMR (DMSO) 14.35 (s, 1H), 9.41 (s, 1H), 8.65 (s, 1H), 8.32-8.14 (m, 1H), 7.94 (s, 1H), 7.66 (s, 3H), 4.22 (s, 2H), 3.11 (t, 3H, J = 11.2 Hz), 2.46 (s, 3H), 2.17 (d, 2H, J = 10.6 Hz), 1.80 (d, 2H, J = 13.0 Hz), 1.62 (d, 1H, J = 12.5 Hz), 1.50-1.40 (m, 2H), 1.27 (q, 2H, J = 12.8 Hz), 1.17-1.06 (m, 2H). ^13^C-NMR (DMSO) 163.3 (d, J = 8.5 Hz), 160.8 (d, J = 8.5 Hz), 144.7, 139.7, 138.0, 128.7, 127.6, 122.0, 110.4, 110.2, 108.0, 65.4, 57.3, 35.0, 28.8, 25.2, 24.5, 21.5, 15.6. HRMS (ESI) m/z [M + H] ^+^: 383.2042 (calculated), 383.2040 (found).

**4C**: R_1_ = R_2_ = F, R_3_ = methylenecyclohexane 63% white solid: ^1^H-NMR (DMSO) δ 14.44 (1H, s), 9.10 (s, 2H), 8.21 (s, 1H), 7.94 (1H, s),7.69-7.64 (m, 3H), 4.20 (s, 2H), 3.17 (s, 1H), 2.83 (d, 2H, J = 6.6 Hz), 2.46 (s, 3H), 1.82-1.61 (m, 7H), 1.23-1.09 (m, 4H), 1.00-0.94 (m, 2H). ^13^C-NMR (DMSO) 163.3 (d, J = 8.3 Hz), 160.9 (d, J = 8.3 Hz), 139.7, 138.0, 128.9, 127.6, 122.0, 110.5, 110.2, 53.4, 49.1, 38.5, 34.7, 30.5, 26.1, 25.5, 21.5. HRMS (ESI) m/z [M + H] ^+^: 397.2199 (calculated), 397.2203 (found).

**4D**: R_1_ = R_2_ = F, R_3_ = 4,4-dimethylcyclohexane 54% white solid: ^1^H-NMR (DMSO) 14.41 (br s, 1H), 9.32 (s, 1H), 8.21 (s, 1H), 7.94 (s, 1H), 7.70-7.64 (m, 3H), 4.23 (s, 2H), 3.06 (t, 1H, J = 11.3 Hz), 2.47 (s, 3H), 1.98 (d, 2H, J = 10.1 Hz), 1.68 (q, 2H, J = 9.3 Hz), 1.45 (d, 2H, J = 13.1 Hz), 1.26-1.20 (m, 2H), 1.09 (t, 1H, J = 7.0 Hz), 0.93 (s, 3H), 0.91 (s, 3H). ^13^C-NMR (DMSO) 163.23 (d, J = 8.5 Hz), 160.8 (d, J = 8.5 Hz), 139.7, 138.0, 128.9, 127.6, 122.0, 110.5, 110.2, 65.4, 57.4, 37.0, 35.3, 32.2, 29.8, 24.6, 24.3, 21.5, 15.6. HRMS (ESI) m/z [M + H] ^+^: 411.2355 (calculated), 411.2359 (found).

**5A**: R_1_ = H, R_2_ = F, R_3_ = 2-aminoindane 80% white solid: ^1^H NMR δ (400 MHz, DMSO) δ 14.26 (s, 1H), 8.42 (br s, 1H), 8.11 (s, 1H), 7.86 (s, 1H), 7.63-7.59 (m, 2H), 7.56-7.51 (m, 2H), 7.21-7.18 (m, 2H), 7.12-7.10 (m, 2H), 3.86 (s, 2H), 3.10 (t, J = 7.6 Hz, 1H), 3.01 (dd, J = 7.0 Hz, J = 5.0 Hz, 2H), 2.76 (dd, J = 6.3 Hz, J = 9.3 Hz, 2H), 2.45 (s, 3H). ^13^C-NMR (DMSO) δ 162.7, 160.3, 143.1, 140.4, 139.5, 139.2, 133.2, 128.9, 127.3, 126.9, 125.0, 123.2, 121.9, 114.1, 113.8, 58.1, 55.4, 42.9, 36.8, 21.5. HRMS (ESI) *m/z* [M + H] ^+^ :399.1985 (calculated), 399.1999 (found).

**5B**: R_1_ = H, R_2_ = F, R_3_ = cyclohexane 77% white solid ^1^H-NMR (DMSO) δ 9.48 (s, 2H), 8.82-8.76 (m, 1H), 8.28 (s, 1H), 7.96 (s, 1H), 7.90 (s, 1H), 7.73 (s, 3H), 4.22 (s, 2H), 3.06 (s, 1H), 2.47 (s, 3H), 2.16 (d, 2H, J = 10.5 Hz), 1.80 (d, 2H, J = 12.3 Hz), 1.62 (d, 1H, J = 11.7 Hz), 1.46 (q, 2H, J = 11.0 Hz), 1.26 (q, 2H, J = 12.5 Hz), 1.14 (t, 1H, J = 12.5 Hz). ^13^C-NMR (DMSO) δ162.8, 160.3, 139.8, 139.2, 133.6, 127.2, 123.2, 122.5, 114.2, 113.9, 56.9, 28.8, 25.2, 24.5, 21.5. HRMS (ESI) *m/z* [M + H] ^+^: 365.2142 (calculated), 365.2158 (found).

**5C**: R_1_ = H, R_2_ = F, R_3_ = methylenecyclohexane 31% white solid: ^1^H-NMR (DMSO) δ 9.36 (s, 2H), 8.70 (s, 1H), 8.26 (s, 1H), 7.95 (s, 1H), 7.87 (t, 1H, J = 7.7 Hz), 7.73-7.71 (m, 3H), 4.22 (s, 2H), 2.82 (s 2H), 2.47 (s, 3H), 1.81 (d, 3H, J = 11.7 Hz), 1.70-1.61 (m, 4H), 1.23-1.15 (m, 4H), 0.96-0.94 (m, 2H). ^13^C-NMR (DMSO) δ 162.8, 160.4, 139.7, 139.2, 133.6, 127.2, 123.21, 122.24, 114.1, 113.9, 55.4, 53.0, 43.8, 34.7, 30.6, 26.0, 25.5, 21.51. HRMS (ESI) *m/z* [M + H] ^+^: 379.2298 (calculated), 379.2291 (found).

**5D**: R_1_ = H, R_2_ = F, R_3_ = 4,4-dimethylcyclohexane 2 ^1^H-NMR (DMSO) δ 9.57 (s, 2H), 8.93 (s, 1H), 8.34 (s, 1H), 7.98 (s 1H), 7.92 (t, 1H, J = 8.0 Hz), 7.73 (t, 2H, J = 7.2 Hz), 4.22 (s, 2H), 3.01 (s, 1H), 2.47 (s, 3H), 1.97 (d, 2H, J = 10.7 Hz), 1.74 (q, 2H, J = 11.6 Hz), 1.44 (d, 2H, J = 13.0 Hz), 1.20 (q, 2H, J = 8.0 Hz), 0.93 (s, 3H), 0.91 (s, 3H). ^13^C-NMR (DMSO) δ 162.7, 160.3, 156.2, 146.2, 143.0, 142.9, 139.9, 139.2, 133.6, 133.5, 129.9, 128.4, 127.3, 123.2, 122.6, 119.4, 119.2, 114.2, 113.9, 56.9, 55.4, 37.0, 32.2, 29.8, 24.5, 24.3. HRMS (ESI) *m/z* [M + H] ^+^: 393.2455 (calculated), 393.2475 (found).

**6A**: R_1_ = R_2_ = H, R_3_ = 2-aminoindane. ^1^H-NMR (DMSO) δ 9.91 (s, 1H), 8.64 (s, 1H), 8.16 (s, 1H), 7.89 (s, 1H), 7.80-7.76 (m, 3H), 7.60 (s, 1H), 7.29-7.27 (m, 2H), 7.21-7.19 (m, 2H), 4.28 (s, 2H), 4.05 (t, 1H, J = 7.83 Hz), 3.39-3.25 (m, 5H), 2.46 (s, 3H). ^13^C-NMR (DMSO) δ 140.4, 139.9, 132.0, 131.3, 127.5, 127.4, 126.4, 125.5, 121.9, 57.7, 49.1, 36.1, 21.6. HRMS (ESI) *m/z* [M + H] ^+^: 381.2079 (calculated), 381.2077 (found).

**6B**: R_1_ = R_2_ = H, R_3_ = cyclohexane. ^1^H-NMR (DMSO) δ 9.31 (s, 1H), 8.56 (s, 1H), 8.18 (s, 1H), 7.90 (S, 1H), 7.81 (d, 2H, J = 8.30 Hz), 7.73 (d, 2H, J = 8.28 Hz), 7.63 (s 1H), 4.20 (s, 2H), 2.99 (t, 1H, J = 3.83 Hz), 2.46 (s, 3H), ), 2.16 (d, 2H, J = 10.35 Hz), 1.79 (d, 2H, J = 13.17 Hz), 1.62 (d, 1H, J = 12.21 Hz), 1.49-1.39 (m, 2H), 1.25-1.13 (m, 3H). ^13^C-NMR (DMSO) δ 140.5, 140.4, 139.5, 132.2, 131.2, 129.1, 127.3, 126.4, 122.1, 65.4, 56.5, 47.0, 28.9, 25.3, 24.5, 21.2, 15.6. HRMS (ESI) *m/z* [M + H] ^+^: 347.2236 (calculated), 347.2249 (found).

**6C**: R_1_ = R_2_ = H, R_3_ = methylenecyclohexane. ^1^H-NMR (DMSO) δ 9.30 (s, 1H), 8.70 (s, 1H), 8.22 (s, 1H), 7.92 (s, 1H), 7.82 (d, 2H, = 8 .29 Hz), 7.72 (d, 2H, J = 8.28 Hz), 7.65 (s 1H), 4.18 (t, 2H, J = 5.25 Hz), 2.78-2.74 (m, 2H), 2.47 (s, 3H), 1.80-1.74 (m, 3H), 1.69-1.60 (m, 4H), 1.22-1.14 (m, 3H), 0.97-0.89 (m, 2H). ^13^C-NMR (DMSO) δ 140.5, 140.4, 139.6, 131.9, 131.4, 129.4, 127.3, 126.6, 122.2, 52.6, 50.5, 34.7, 30.6, 26.01, 25.5, 21.6. HRMS (ESI) *m/z* [M + H] ^+^: 361.2392 (calculated), 361.2400 (found).

**6D**: (157D) R_1_ = R_2_ = H, R_3_ = 4,4-dimethylcyclohexane. ^1^H-NMR (DMSO) δ 9.32 (s, 1H), 8.50 (S, 1H), 8.17 (s, 1H), 7.89 (s, 1H), 7.81 (d, 2H, J = 8.24 Hz), 7.72 (d, 2H, J = 8.15 Hz), 7.61 (s, 1H), 4.20 (d, 2H, J = 5.39 Hz), 3.17 (s, 1H), 2.96-2.92 (m, 1H), 2.46 (s, 3H), 1.96 (d, 2H, J = 9.86 Hz), 1.70 (q, 2H, J = 11.61 Hz), 1.44 (d, 2H, J = 13.00 Hz), 1.22-1.16 (m, 2H), 0.93 (s, 3H), 0.91 (s, 3H). ^13^C-NMR (DMSO) δ 140.6, 140.4, 139.4, 132.2, 131.2, 128.9, 127.3, 126.4, 122.0, 65.4, 56.5, 47.2, 37.0, 32.2, 29.8, 24.6, 24.3, 21.6, 15.6.

Radiochemistry. Radiosynthesis of [^11^C]**1A** was performed in a Synthra MeI Plus module with [^11^C] CH_3_I that was prepared via the following gas phase method. [^11^C]CO_2_ was produced by ^14^N(p,α)^11^C nuclear reaction. This was reduced to [^11^C]CH_4_ using Ni catalyst and then, converted to [^11^C]CH_3_I through an iodine column. The nascent [^11^C]CH_3_I was bubbled directly into the reaction vial containing the precursor. The mixture was irradiated at 450 nm for 5 min in a custom made integrated photoreactor designed by Synthra. Subsequently, the intermediate was hydrolyzed with piperidine for 2 min at ambient temperature. The reaction was quenched at RT with 700 μL of HPLC mobile phase. The crude product was purified by semi-preparative HPLC (column: Phenomenex Gemini 5 μm C18 100 Å, 10 × 250 mm; eluent: CH_3_CN/pH 10 sodium carbonate/sodium bicarbonate buffer (55:45); wavelength: 254 nm; flow rate: 4.0 mL/min; R_t_ [^11^C] **1A** = 15 min.

Radiochemical purity and the molar activity were determined by analytical UPLC prior to the PET study (column: Waters Acquity UHPLC-BEH C18 3.5 μm C18 130 Å, 4.6 × 50 mm; eluent: CH_3_CN/ 0.1% TFA in water (28:72); wavelength: 254 nm; flow rate: 0.5 mL/ min; **1A** R_t_ = 1.26 min.

PET Studies of [^11^C]**1A** in Nonhuman Primates. Dynamic total-body **[^11^C]1A** PET scan wAS performed on 2 adult male rhesus macaques using the PennPET Explorer scanner A baseline of **[^11^C]1A** and a naloxone pretreatment (0.14 mg/kg, intramuscular administration 10 min prior to **[^11^C]1A** injection) studies were taken on 19 years old, 10 kg male rhesus macaque in two different days (injection dose of **[^11^C]1A**: 82.9 and 61.8 MBq respectively). The monkey, fasted for 12 h prior to the PET study, was initially anesthetized by intramuscular injection with ketamine (4 mg/kg) and dexmedetomidine (0.05 mg/kg). The monkey was intubated, and anesthesia was maintained with 0.75-2% isoflurane/oxygen. A percutaneous catheter was placed for the tracer injection. A low dose CT scan was performed to confirm positioning and attenuation correction followed by 90 min dynamic PET image acquired for 90 min (12×10 sec, 2×30 sec, 4×60 sec, 7×120 sec, 4×240 sec and 8×300 sec) in list mode after venous injection of **[^11^C]1A.** The PET image was reconstructed using time-of-flight list-mode ordered subsets expectation maximization (OSEM, 25 subsets) reconstruction algorithm.^44^ The PET/CT imaging data was analyzed by using Pmod software (version 3.7, PMOD Technologies Ltd., Zurich, Switzerland). Six volumes of interest (VOIs) for brain including caudate, putamen, thalamus, frontal cortex, occipital cortex, and cerebellar cortex were manually delineated on a T1-weighted MR image. Time activity curves were extracted from all the VOIs and expressed as standardized uptake value (SUV).

Molecular Docking. Compound **1A**, **3A**, naloxone, and GSK1521498 were performed docking studies on mu-opioid receptor. Open Babel v3.1.0^45^ were used for predicting protonated status at physiological pH of each compound. Then, the protonated 3-dimensional structures were imported to UCSF Chimera version 1.17.3 and minimized using AMBER ff99SB force field calculations for preparation for molecular docking studies.^46^ Molecular docking studies were performed via the AutoDock 4.2^47^ plugin on PyMOL (pymol.org). The X-ray crystal structure of mu-opioid receptor (PDB ID 4DKL, resolution 2.80 Å) was obtained from the RCSB Protein Data Bank (https://www.rcsb.org). Water molecular outside the 5 Å radius of the crystallographic ligand (morphinan antagonist), and heteroatoms including cholesterol, ions, and other small molecules were removed from the structure. Then, the protein was imported to H++ web server (http://biophysics.cs.vt.edu/H++)^48–50^ to protonate the mu-opioid receptor structure at pH 7.0. A grid box with a dimension of 24 x 24 x 24 Å^3^ was applied to the mu-opioid receptor structure covering binding pocket. The Lamarckian Genetic Algorithm with a maximum of 2,500,000 energy evaluations was used to calculate 100 mu-opioid receptor-ligand binding poses for each compound. The mu-opioid receptor-ligand complex that reproduced the crystallographic ligand binding pose and with good docking score was reported for each compound.

## Acknowledgements

We greatly appreciate the efforts of Synthra, who designed the integrated photoreactor used in the radiolabeling experiments described in this manuscript. We are also grateful to the PDSP at the University of North Carolina at Chapel Hill under the direction of Dr. Bryan L. Roth. We acknowledge the work of cyclotron engineers Larry Toto and David Schuab.

## References

(1) Brownstein, M. J. A Brief History of Opiates, Opioid Peptides, and Opioid Receptors. Proc. Natl. Acad. Sci. 1993, 90 (12), 5391–5393. 10.1073/pnas.90.12.5391.

(2) Ogura, T.; Egan, T. Chapter 15. Opioid Agonists and Antagonists. Pharmacol. Physiol. Anesth. Found. Clin. Appl. 2013, 253–271. 10.1016/B978-1-4377-1679-5.00015-6.

(3) Matthys, H.; Bleicher, B.; Bleicher, U. Dextromethorphan and Codeine: Objective Assessment of Antitussive Activity in Patients with Chronic Cough. J. Int. Med. Res. 1983, 11 (2), 92–100. 10.1177/030006058301100206.

(4) Hanauer, S. The Role of Loperamide in Gastrointestinal Disorders. Rev. Gastroenterol. Disord. 2008, 8, 15–20.

(5) Ryan, S.; Dunne, R. Pharmacokinetic Properties of Intranasal and Injectable Formulations of Naloxone for Community Use: A Systematic Review. Pain Manag. 2018, 8. 10.2217/pmt-2017-0060.

(6) Aboujaoude, E.; Salame, W. Naltrexone: A Pan-Addiction Treatment? CNS Drugs 2016, 30. 10.1007/s40263-016-0373-0.

(7) Thomas, J.; Karver, S.; Cooney, G. A.; Chamberlain, B. H.; Watt, C. K.; Slatkin, N. E.; Stambler, N.; Kremer, A. B.; Israel, R. J. Methylnaltrexone for Opioid-Induced Constipation in Advanced Illness. N. Engl. J. Med. 2008, 358 (22), 2332–2343. 10.1056/NEJMoa0707377.

(8) Bohn, L. M.; Lefkowitz, R. J.; Gainetdinov, R. R.; Peppel, K.; Caron, M. G.; Lin, F.-T. Enhanced Morphine Analgesia in Mice Lacking β-Arrestin 2. Science 1999, 286 (5449), 2495– 2498. 10.1126/science.286.5449.2495.

(9) Manglik, A.; Lin, H.; Aryal, D. K.; McCorvy, J. D.; Dengler, D.; Corder, G.; Levit, A.; Kling, R. C.; Bernat, V.; Hübner, H.; Huang, X.-P.; Sassano, M. F.; Giguère, P. M.; Löber, S.; Da Duan; Scherrer, G.; Kobilka, B. K.; Gmeiner, P.; Roth, B. L.; Shoichet, B. K. Structure-Based Discovery of Opioid Analgesics with Reduced Side Effects. Nature 2016, 537 (7619), 185–190. 10.1038/nature19112.

(10) Jae Min Jeong. Radiopharmaceutical Chemistry. J. Nucl. Med. 2019, 60 (12), 1833. 10.2967/jnumed.119.237479.

(11) Drzezga, A.; Bischof, G. N.; Giehl, K.; van Eimeren, T. Chapter 67 - PET and SPECT Imaging of Neurodegenerative Diseases. In Molecular Imaging (Second Edition); Ross, B. D., Gambhir, S. S., Eds.; Academic Press, 2021; pp 1309–1334. 10.1016/B978-0-12-816386-3.00085-5.

(12) Zhu, L.; Ploessl, K.; Kung, H. F. PET/SPECT Imaging Agents for Neurodegenerative Diseases. Chem. Soc. Rev. 2014, 43 (19), 6683–6691. 10.1039/C3CS60430F.

(13) Klunk, W. E.; Engler, H.; Nordberg, A.; Wang, Y.; Blomqvist, G.; Holt, D. P.; Bergström, M.; Savitcheva, I.; Huang, G.-F.; Estrada, S.; Ausén, B.; Debnath, M. L.; Barletta, J.; Price, J. C.; Sandell, J.; Lopresti, B. J.; Wall, A.; Koivisto, P.; Antoni, G.; Mathis, C. A.; Långström, B. Imaging Brain Amyloid in Alzheimer’s Disease with Pittsburgh Compound-B. Ann. Neurol. 2004, 55 (3), 306–319. 10.1002/ana.20009.

(14) Werry, E. L.; Bright, F. M.; Piguet, O.; Ittner, L. M.; Halliday, G. M.; Hodges, J. R.; Kiernan, M. C.; Loy, C. T.; Kril, J. J.; Kassiou, M. Recent Developments in TSPO PET Imaging as A Biomarker of Neuroinflammation in Neurodegenerative Disorders. Int. J. Mol. Sci. 2019, 20 (13). 10.3390/ijms20133161.

(15) Hou, C.; Hsieh, C.-J.; Li, S.; Lee, H.; Graham, T. J.; Xu, K.; Weng, C.-C.; Doot, R. K.; Chu, W.; Chakraborty, S. K.; Dugan, L. L.; Mintun, M. A.; Mach, R. H. Development of a Positron Emission Tomography Radiotracer for Imaging Elevated Levels of Superoxide in Neuroinflammation. ACS Chem. Neurosci. 2018, 9 (3), 578–586. 10.1021/acschemneuro.7b00385.

(16) Fowler, J. S.; Volkow, N. D. PET Imaging Studies in Drug Abuse. J. Toxicol. Clin. Toxicol. 1998, 36 (3), 163–174. 10.3109/15563659809028936.

17. Wiers, C. E.; Cabrera, E.; Skarda, E.; Volkow, N. D.; Wang, G.-J. Chapter 9 - PET Imaging for Addiction Medicine: From Neural Mechanisms to Clinical Considerations. In Progress in Brain Research; Ekhtiari, H., Paulus, M. P., Eds.; Elsevier, 2016; Vol. 224, pp 175–201. 10.1016/bs.pbr.2015.07.016.

(18) Kang, Y.; O’Conor, K. A.; Kelleher, A. C.; Ramsey, J.; Bakhoda, A.; Eisenberg, S. M.; Zhao, W.; Stodden, T.; Pearson, T. D.; Guo, M.; Brown, N.; Liow, J.-S.; Fowler, J. S.; Kim, S. W.; Volkow, N. D. Naloxone’s Dose-Dependent Displacement of [11C]Carfentanil and Duration of Receptor Occupancy in the Rat Brain. Sci. Rep. 2022, 12 (1), 6429. 10.1038/s41598-022-09601-2.

(19) Matthews, P. M.; Rabiner, E. A.; Passchier, J.; Gunn, R. N. Positron Emission Tomography Molecular Imaging for Drug Development. Br. J. Clin. Pharmacol. 2012, 73 (2), 175–186. 10.1111/j.1365-2125.2011.04085.x.

(20) Hirvonen, J.; Aalto, S.; Hagelberg, N.; Maksimow, A.; Ingman, K.; Oikonen, V.; Virkkala, J.; Någren, K.; Scheinin, H. Measurement of Central μ-Opioid Receptor Binding in Vivo with PET and [11C]Carfentanil: A Test-Retest Study in Healthy Subjects. Eur. J. Nucl. Med. Mol. Imaging 2008, 36, 275–286. 10.1007/s00259-008-0935-6.

(21) Frost, J. J.; Douglass, K. H.; Mayberg, H. S.; Dannals, R. F.; Links, J. M.; Wilson, A. A.; Ravert, H. T.; Crozier, W. C.; Wagner, H. N. Multicompartmental Analysis of [11C]-Carfentanil Binding to Opiate Receptors in Humans Measured by Positron Emission Tomography. J. Cereb. Blood Flow Metab. 1989, 9 (3), 398–409. 10.1038/jcbfm.1989.59.

(22) Rabiner, E. A.; Beaver, J.; Makwana, A.; Searle, G.; Long, C.; Nathan, P. J.; Newbould, R. D.; Howard, J.; Miller, S. R.; Bush, M. A.; Hill, S.; Reiley, R.; Passchier, J.; Gunn, R. N.; Matthews, P. M.; Bullmore, E. T. Pharmacological Differentiation of Opioid Receptor Antagonists by Molecular and Functional Imaging of Target Occupancy and Food Reward-Related Brain Activation in Humans. Mol. Psychiatry 2011, 16 (8), 826–835. 10.1038/mp.2011.29.

(23) Newberg, A.; Ray, R.; Scheuermann, J.; Wintering, N.; Reddin, J.; Schmitz, A.; Freifelder, R.; Karp, J.; Lerman, C.; Divgi, C. Dosimetry of11C-Carfentanil, a μ-Opioid Receptor Imaging Agent. Nucl. Med. Commun. 2009, 30, 314–318. 10.1097/MNM.0b013e328329a0ec.

(24) Liu, S.; Kim, D.-I.; Oh, T. G.; Pao, G.; Kim, J.-H.; Palmiter, R.; Banghart, M.; Lee, K.-F.; Evans, R.; Han, S. Neural Basis of Opioid-Induced Respiratory Depression and Its Rescue. Proc. Natl. Acad. Sci. 2021, 118, e2022134118. 10.1073/pnas.2022134118.

(25) Algera, M. H.; Kamp, J.; van der Schrier, R.; van Velzen, M.; Niesters, M.; Aarts, L.; Dahan, A.; Olofsen, E. Opioid-Induced Respiratory Depression in Humans: A Review of Pharmacokinetic–Pharmacodynamic Modelling of Reversal. Br. J. Anaesth. 2019, 122 (6), e168–e179. 10.1016/j.bja.2018.12.023.

(26) Renata C.N. Marchette; Erika R. Carlson; Emma V. Frye; Lyndsay E. Hastings; Janaina C.M. Vendruscolo; Gustavo Mejias-Torres; Stephen J. Lewis; Aidan Hampson; Nora D. Volkow; Leandro F. Vendruscolo; George F. Koob. Heroin- and Fentanyl-Induced Respiratory Depression in a Rat Plethysmography Model: Potency, Tolerance, and Sex Differences. J. Pharmacol. Exp. Ther. 2023, 385 (2), 117. 10.1124/jpet.122.001476.

(27) Rafique, W.; Khanapur, S.; Spilhaug, M. M.; Riss, P. J. Reaching out for Sensitive Evaluation of the Mu Opioid Receptor in Vivo: Positron Emission Tomography Imaging of the Agonist [11C]AH7921. ACS Chem. Neurosci. 2017, 8 (9), 1847–1852. 10.1021/acschemneuro.7b00075.

(28) Rothman, R. B.; McLean, S. An Examination of the Opiate Receptor Subtypes Labeled by [3H]Cyclofoxy: An Opiate Antagonist Suitable for Positron Emission Tomography. Biol. Psychiatry 1988, 23 (5), 435–458. 10.1016/0006-3223(88)90016-9.

(29) Rothman, R. B.; Bykov, V.; Reid, A.; De Costa, B. R.; Newman, A.-H.; Jacobson, A. E.; Rice, K. C. A Brief Study of the Selectivity of Norbinaltorphimine, (−)-Cyclofoxy, and (+)-Cyclofoxy among Opioid Receptor Subtypes in Vitro. Neuropeptides 1988, 12 (3), 181–187. 10.1016/0143-4179(88)90052-2.

(30) Carson, R. E.; Channing, M. A.; Blasberg, R. G.; Dunn, B. B.; Cohen, R. M.; Rice, K. C.; Herscovitch, P. Comparison of Bolus and Infusion Methods for Receptor Quantitation: Application to [18F]Cyclofoxy and Positron Emission Tomography. J. Cereb. Blood Flow Metab. 1993, 13 (1), 24–42. 10.1038/jcbfm.1993.6.

(31) Quelch, D. R.; Katsouri, L.; Nutt, D. J.; Parker, C. A.; Tyacke, R. J. Imaging Endogenous Opioid Peptide Release with [11C]Carfentanil and [3H]Diprenorphine: Influence of Agonist-Induced Internalization. J. Cereb. Blood Flow Metab. 2014, 34 (10), 1604–1612. 10.1038/jcbfm.2014.117.

(32) Giuliano, C.; Goodlett, C. R.; Economidou, D.; García-Pardo, M. P.; Belin, D.; Robbins, T. W.; Bullmore, E. T.; Everitt, B. J. The Novel μ-Opioid Receptor Antagonist GSK1521498 Decreases Both Alcohol Seeking and Drinking: Evidence from a New Preclinical Model of Alcohol Seeking. Neuropsychopharmacology 2015, 40 (13), 2981–2992. 10.1038/npp.2015.152.

(33) Cambridge, V. C.; Ziauddeen, H.; Nathan, P. J.; Subramaniam, N.; Dodds, C.; Chamberlain, S. R.; Koch, A.; Maltby, K.; Skeggs, A. L.; Napolitano, A.; Farooqi, I. S.; Bullmore, E. T.; Fletcher, P. C. Neural and Behavioral Effects of a Novel Mu Opioid Receptor Antagonist in Binge-Eating Obese People. Food Addict. 2013, 73 (9), 887–894. 10.1016/j.biopsych.2012.10.022.

(34) Diane M. Ignar; Aaron S. Goetz; Kimberly Nichols Noble; Luz Helena Carballo; Andrea E. Stroup; Julie C. Fisher; Joyce A. Boucheron; Tracy A. Brainard; Andrew L. Larkin; Andrea H. Epperly; Todd W. Shearer; Scott D. Sorensen; Jason D. Speake; Jonathan D. Hommel. Regulation of Ingestive Behaviors in the Rat by GSK1521498, a Novel μ-Opioid Receptor-Selective Inverse Agonist. J. Pharmacol. Exp. Ther. 2011, 339 (1), 24. 10.1124/jpet.111.180943.

(35) Pipal, R. W.; Stout, K. T.; Musacchio, P. Z.; Ren, S.; Graham, T. J. A.; Verhoog, S.; Gantert, L.; Lohith, T. G.; Schmitz, A.; Lee, H. S.; Hesk, D.; Hostetler, E. D.; Davies, I. W.; MacMillan, D. W. C. Metallaphotoredox Aryl and Alkyl Radiomethylation for PET Ligand Discovery. Nature 2021, 589 (7843), 542–547. 10.1038/s41586-020-3015-0.

(36) Kidjemet, D. N,N-Dimethylformamide Dimethyl Acetal. Synlett 2002, 2002 (10), 1741–1742. 10.1055/s-2002-34251.

(37) Besnard, J.; Ruda, G. F.; Setola, V.; Abecassis, K.; Rodriguiz, R. M.; Huang, X.-P.; Norval, S.; Sassano, M. F.; Shin, A. I.; Webster, L. A.; Simeons, F. R. C.; Stojanovski, L.; Prat, A.; Seidah, N. G.; Constam, D. B.; Bickerton, G. R.; Read, K. D.; Wetsel, W. C.; Gilbert, I. H.; Roth, B. L.; Hopkins, A. L. Automated Design of Ligands to Polypharmacological Profiles. Nature 2012, 492 (7428), 215–220. 10.1038/nature11691.

(38) Kroeze, W. K.; Sassano, M. F.; Huang, X.-P.; Lansu, K.; McCorvy, J. D.; Giguère, P. M.; Sciaky, N.; Roth, B. L. PRESTO-Tango as an Open-Source Resource for Interrogation of the Druggable Human GPCRome. Nat. Struct. Mol. Biol. 2015, 22 (5), 362–369. 10.1038/nsmb.3014.

(39) Daina, A.; Zoete, V. A BOILED-Egg To Predict Gastrointestinal Absorption and Brain Penetration of Small Molecules. ChemMedChem 2016, 11 (11), 1117–1121. 10.1002/cmdc.201600182.

(40) Sun, S.; Jia, Q.; Zhang, Z. Applications of Amide Isosteres in Medicinal Chemistry. Bioorg. Med. Chem. Lett. 2019, 29 (18), 2535–2550. 10.1016/j.bmcl.2019.07.033.

(41) Gillis, E. P.; Eastman, K. J.; Hill, M. D.; Donnelly, D. J.; Meanwell, N. A. Applications of Fluorine in Medicinal Chemistry. J. Med. Chem. 2015, 58 (21), 8315–8359. 10.1021/acs.jmedchem.5b00258.

(42) Pike, V. W. PET Radiotracers: Crossing the Blood–Brain Barrier and Surviving Metabolism. Trends Pharmacol. Sci. 2009, 30 (8), 431–440. 10.1016/j.tips.2009.05.005.

(43) Hsieh, C.-J.; Hou, C.; Lee, H. S.; Tomita, C.; Schmitz, A.; Plakas, K.; Dubroff, J. G.; Mach, R. H. Total-Body Imaging of Mu-Opioid Receptors with [11C]Carfentanil in Non-Human Primates. Eur. J. Nucl. Med. Mol. Imaging 2024.

(44) Pantel, A. R.; Viswanath, V.; Daube-Witherspoon, M. E.; Dubroff, J. G.; Muehllehner, G.; Parma, M. J.; Pryma, D. A.; Schubert, E. K.; Mankoff, D. A.; Karp, J. S. PennPET Explorer: Human Imaging on a Whole-Body Imager. J. Nucl. Med. 2020, 61 (1), 144. 10.2967/jnumed.119.231845.

(45) O’Boyle, N. M.; Banck, M.; James, C. A.; Morley, C.; Vandermeersch, T.; Hutchison, G. R. Open Babel: An Open Chemical Toolbox. J. Cheminformatics 2011, 3 (1), 33. 10.1186/1758-2946-3-33.

(46) Pettersen, E. F.; Goddard, T. D.; Huang, C. C.; Couch, G. S.; Greenblatt, D. M.; Meng, E. C.; Ferrin, T. E. UCSF Chimera—A Visualization System for Exploratory Research and Analysis. J. Comput. Chem. 2004, 25 (13), 1605–1612. 10.1002/jcc.20084.

(47) Morris, G. M.; Huey, R.; Lindstrom, W.; Sanner, M. F.; Belew, R. K.; Goodsell, D. S.; Olson, A. J. AutoDock4 and AutoDockTools4: Automated Docking with Selective Receptor Flexibility. J. Comput. Chem. 2009, 30 (16), 2785–2791. 10.1002/jcc.21256.

(48) Anandakrishnan, R.; Aguilar, B.; Onufriev, A. V. H++ 3.0: Automating pK Prediction and the Preparation of Biomolecular Structures for Atomistic Molecular Modeling and Simulations. Nucleic Acids Res. 2012, 40 (W1), W537–W541. 10.1093/nar/gks375.

(49) Gordon, J. C.; Myers, J. B.; Folta, T.; Shoja, V.; Heath, L. S.; Onufriev, A. H++: A Server for Estimating p Ka s and Adding Missing Hydrogens to Macromolecules. Nucleic Acids Res. 2005, 33 (suppl_2), W368–W371. 10.1093/nar/gki464.

(50) Myers, J.; Grothaus, G.; Narayanan, S.; Onufriev, A. A Simple Clustering Algorithm Can Be Accurate Enough for Use in Calculations of pKs in Macromolecules. Proteins Struct. Funct. Bioinforma. 2006, 63 (4), 928–938. 10.1002/prot.20922.

