## Supplemental Information for "A Small-molecule Antagonist Radiotracer for Positron Emission Tomography Imaging of the Mu Opioid Receptor"

Table S1. Binding Profile of Final Compounds

| **GPCR** | | **1A** | **2A** | **3A** | **4A** | **5A** | **6A** |
| --- | --- | --- | --- | --- | --- | --- | --- |
| Dopamine | D1 | 997.9 | 1273.5 | 1318 | 1866.3 | 1268.5 | 716.5 |
|  | D2 | 6531.3 | > 10,000 | > 10,000 | > 10,000 | > 10,000 | > 10,000 |
|  | D3 | 465.6 | 747.8 | > 10,000 | > 10,000 | 181 | > 10,000 |
|  | D4 | 594.2 | 748.3 | 27.2 | 1767.3 | 5501.7 | 1771.1 |
|  | D5 | > 10,000 | > 10,000 | > 10,000 | > 10,000 | > 10,000 | > 10,000 |
|  | DAT | 532.7 | 14.3 | 849.8 | 497.2 | 302.5 | 286.6 |
| Serotonin | 5HT1A | > 10,000 | > 10,000 | > 10,000 | > 10,000 | > 10,000 | > 10,000 |
|  | 5HT1B | N.A. | N.A. | N.A. | N.A. | 3475.6 | N.A. |
|  | 5HT1D | N.A. | 1655.4 | > 10,000 | N.A. | 1422.3 | 1323.4 |
|  | 5HT1E | > 10,000 | > 10,000 | N.A. | N.A. | > 10,000 | > 10,000 |
|  | 5HT2A | > 10,000 | > 10,000 | > 10,000 | > 10,000 | 1024.9 | 1000 |
|  | 5HT2B | > 10,000 | > 10,000 | > 10,000 | 483.6 | 948.2 | 825.7 |
|  | 5HT2C | > 10,000 | > 10,000 | 2268.8 | > 10,000 | > 10,000 | 470.3 |
|  | 5HT3 | > 10,000 | > 10,000 | > 10,000 | > 10,000 | > 10,000 | > 10,000 |
|  | 5HT5A | > 10,000 | > 10,000 | > 10,000 | > 10,000 | 8044.5 | 3942.8 |
|  | 5HT6 | > 10,000 | > 10,000 | > 10,000 | > 10,000 | > 10,000 | > 10,000 |
|  | 5HT7A | > 10,000 | 890.8 | 976.8 | > 10,000 | 707.1 | 746.6 |
|  | SERT | 523.4 | 173.7 | 310.3 | 334.2 | 115.5 | 129.6 |
| Histamine | H1 | > 10,000 | > 10,000 | > 10,000 | > 10,000 | 6181.6 | 1454.5 |
|  | H2 | N.A. | N.A. | N.A. | N.A. | N.A. | N.A. |
|  | H3 | > 10,000 | > 10,000 | > 10,000 | 1253.1 | 1885.4 | 474.2 |
|  | H4 | > 10,000 | > 10,000 | > 10,000 | > 10,000 | > 10,000 | > 10,000 |
| Opioid | MOR | 8.3 | 9.2 | 7.8 | 17.1 | 13.7 | 4.2 |
|  | DOR | 410.3 | 350 | 440.7 | 228.7 | 171.4 | 98 |
|  | KOR | 308.3 | 104.8 | 67.6 | 246.2 | 67.4 | 19.4 |
|  | NOP | > 10,000 | > 10,000 | > 10,000 | > 10,000 | 559.6 | 167 |

Table S1. Continued.

| **GPCR** | | **1A** | **2A** | **3A** | **4A** | **5A** | **6A** |
| --- | --- | --- | --- | --- | --- | --- | --- |
| Muscarinic | M1 | > 10,000 | 558.3 | 1441.5 | > 10,000 | > 10,000 | 3376.8 |
|  | M2 | 396.8 | 153.3 | 31.6 | 628.2 | 154.1 | 12.5 |
|  | M3 | 1619.2 | 509.6 | 954.6 | 1262.7 | 1299.6 | 234.9 |
|  | M4 | 454.5 | 182.2 | 263.9 | 603.8 | 311.4 | 326.5 |
|  | M5 | > 10,000 | > 10,000 | 708.1 | > 10,000 | 1284.7 | 472.9 |
| Adrenergic | alpha1A | 1215.1 | 462.2 | 631.4 | 1768.9 | 497.9 | 1713.2 |
|  | alpha1B | > 10,000 | > 10,000 | > 10,000 | 4026.2 | 3474.6 | 8400.4 |
|  | alpha1D | > 10,000 | > 10,000 | 657.6 | 855.9 | 1051 | 991.5 |
|  | alpha2A | 2160.7 | 1365.8 | 132.5 | > 10,000 | 601.3 | 616 |
|  | beta1 | > 10,000 | > 10,000 | N.A. | N.A. | > 10,000 | > 10,000 |
|  | beta2 | > 10,000 | > 10,000 | N.A. | N.A. | > 10,000 | > 10,000 |
|  | beta3 | N.A. | N.A. | N.A. | N.A. | N.A. | N.A. |
| Sigma | sigma1 | 405.5 | 188.2 | 429 | 649.1 | 173.3 | 191.5 |
|  | sigma2 | 849.4 | 578.8 | 421.1 | 426.4 | 557.3 | 113.5 |
| NET | | 281 | 153.9 | 46.4 | 446.2 | 73.7 | 14.4 |

Comprehensive binding profiles for aminoindane functionalized final compounds across a variety of GPCRs. Each number represents the average of 3 experiments. N.A. = Not Active.

Figure S1. Full radioligand competition binding curves for -aminoindane functionalized final compounds across the opioid family of receptors. Each data point represents the average of three individual experiments.

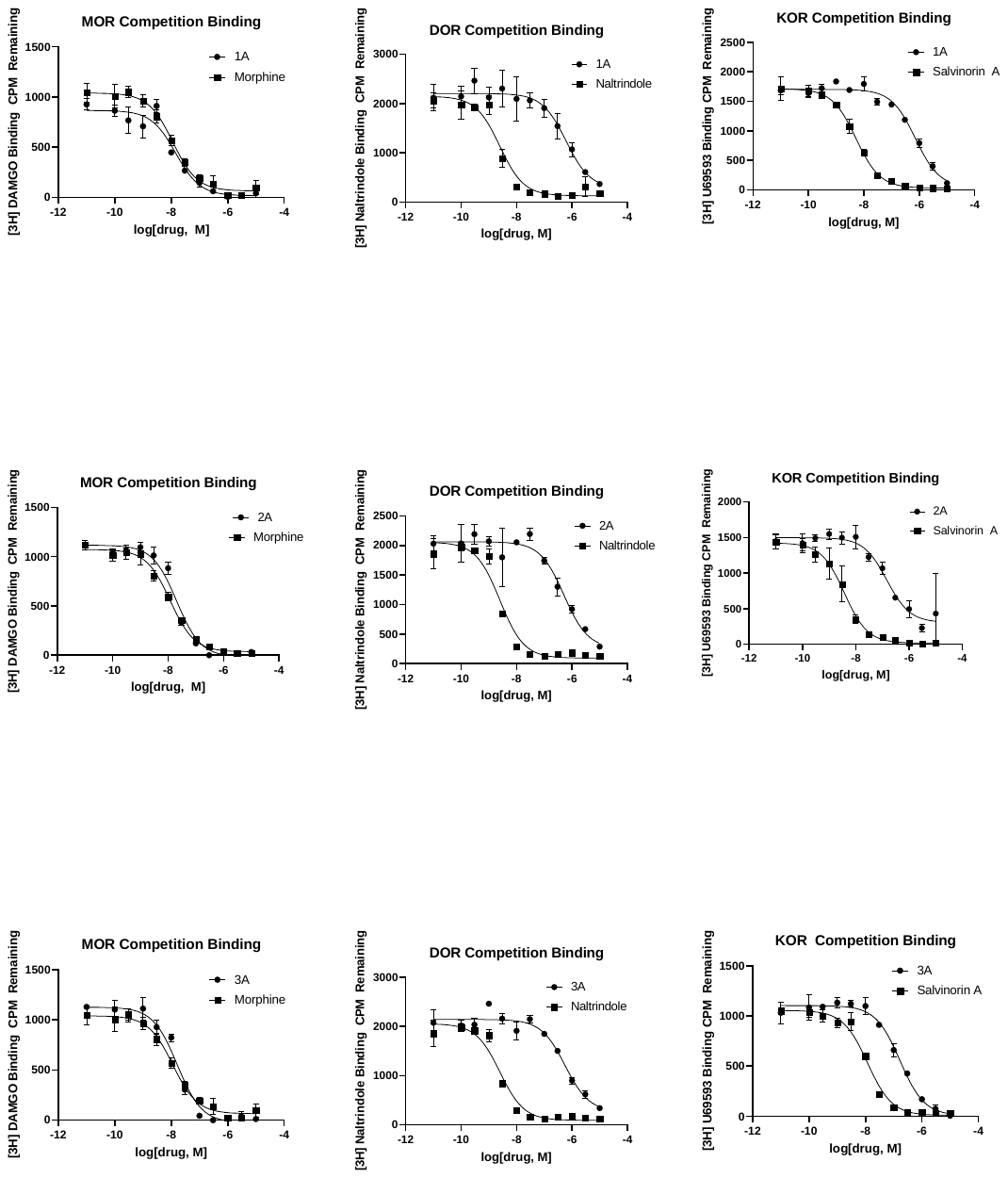

Figure S1 Continued.

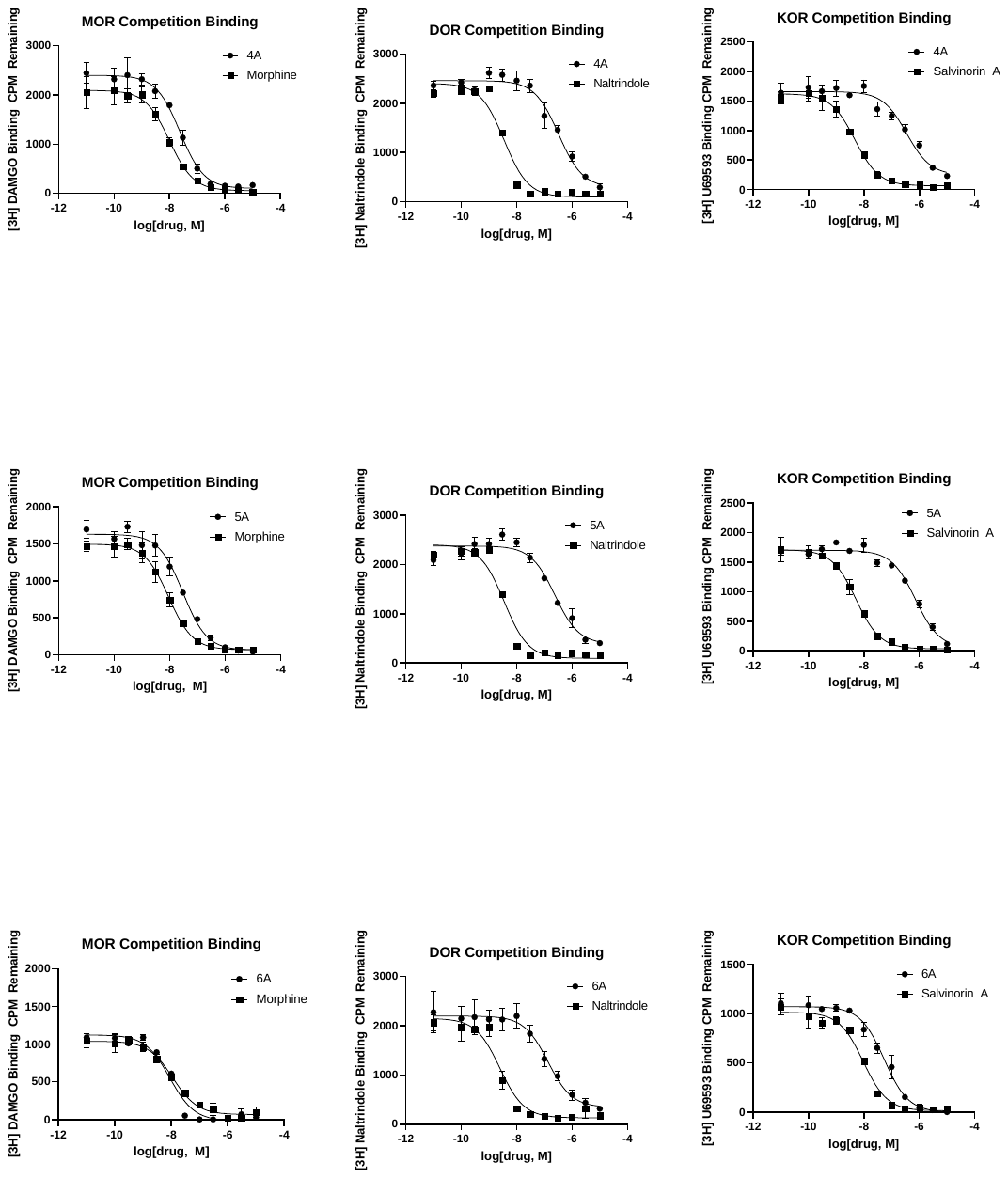

Figure S2: The intrinsic activity of ligands 1A and 3A were determined through the PRESTO-Tango assay. Each data point represents the average of three experiments. Ligands were screened at the Psychoactive Drug Screening Program (PDSP).

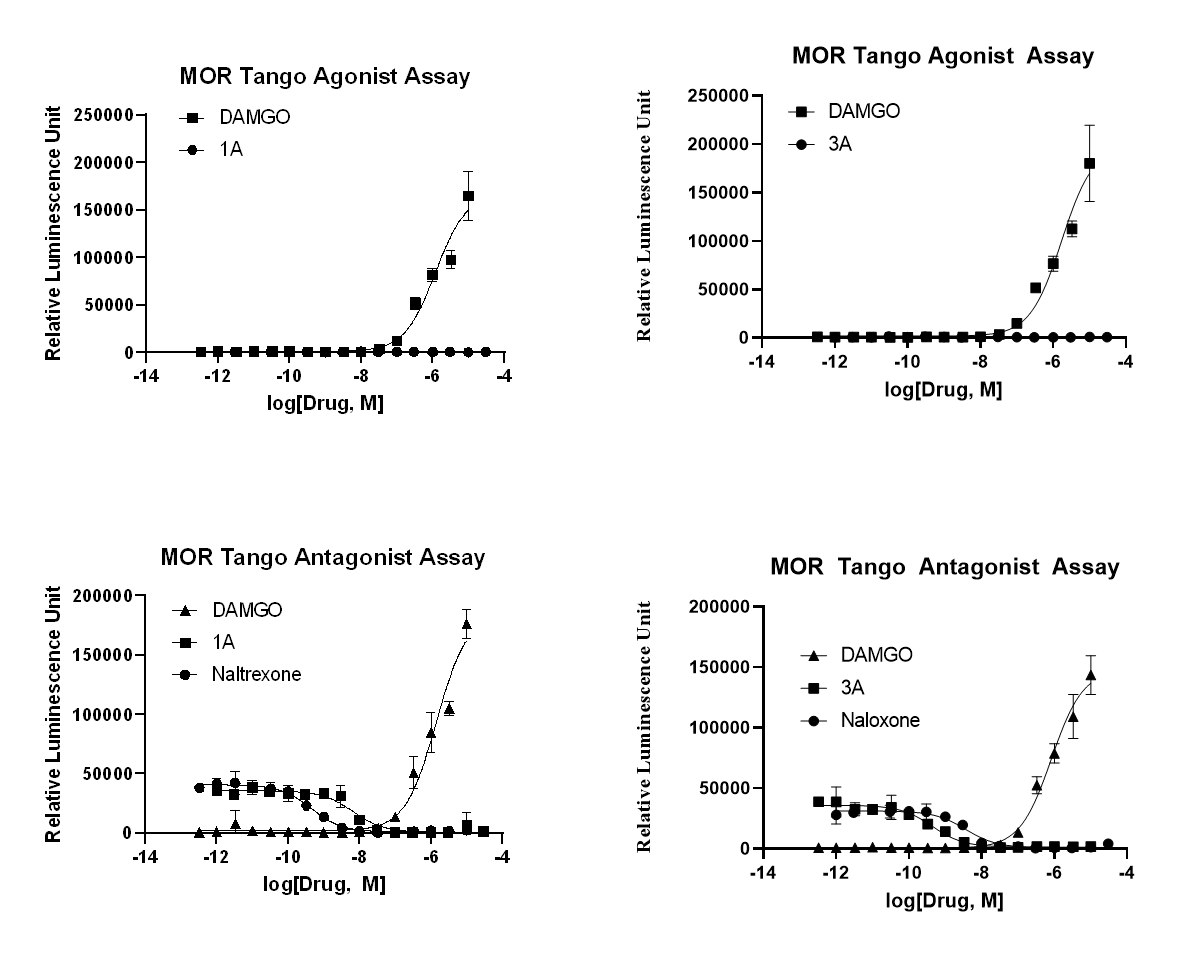

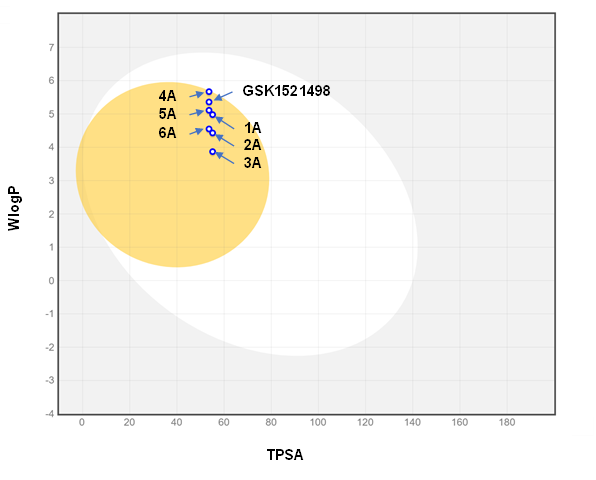

Figure S3: Brain Or IntestinaL (BOIL-ed Egg) Plot Demonstrating the predicted BBB permeability of -aminnoindane containing molecules entities 1A 🡪 6A. These ligands were chosen for predicted BBB-permeability studies based on their MOR binding potency and selectivity. Methylation of GSK1521498 yields 4A, a congener with increased lipophilicity compared to the lead compound. The lipophilicity of 4A was decreased through either triazole-benzamide substitution as in the order of 4A 🡪 1A or through iterative fluorine-hydrogen substitution in case order of 4A 🡪 5A 🡪 6A.

Figure S4. Semi Preparative HPLC chromatograms for the purification of [^11^C] 1A.

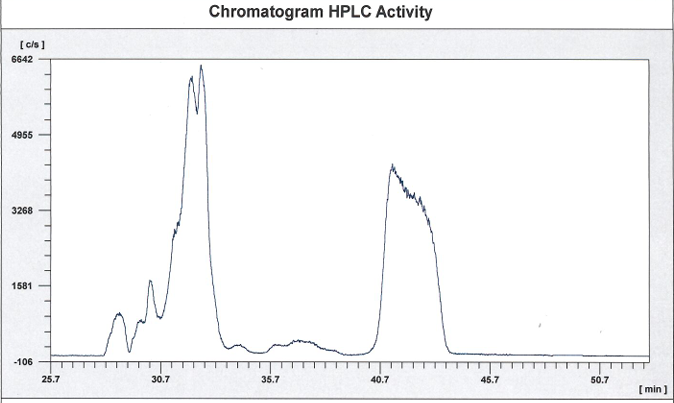

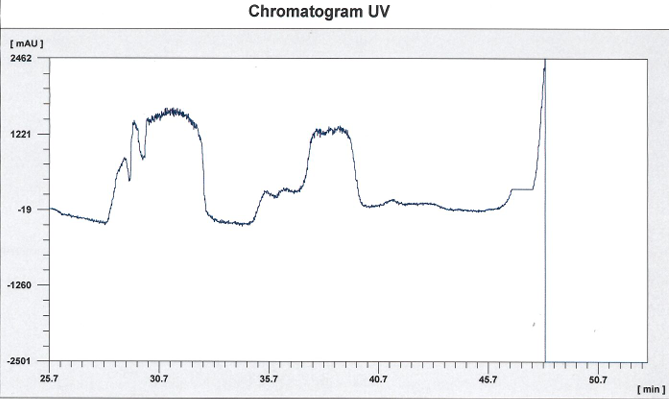

Figure S5. Analytical HPLC chromatogram for the quality control analysis of [^11^C] 1A.

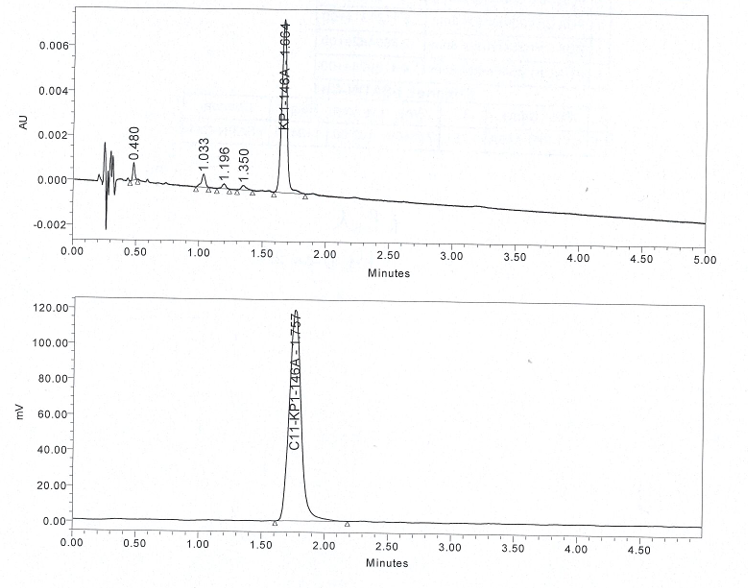

Figure S6. UPLC validation of 1A.

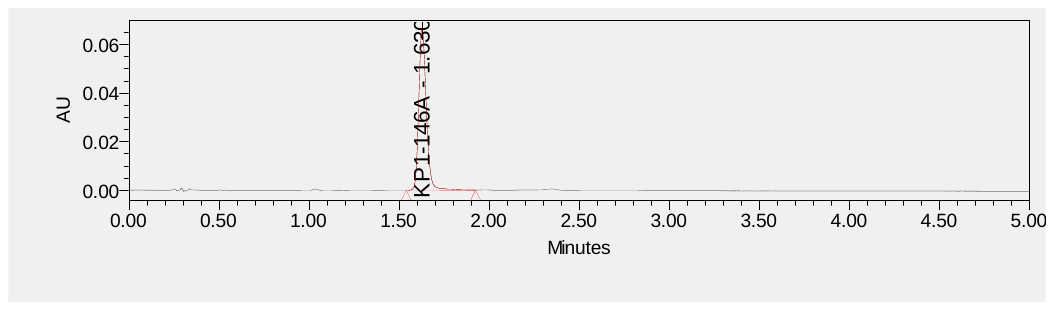

Synthetic Procedures:

Synthesis of Compounds **7A-C**

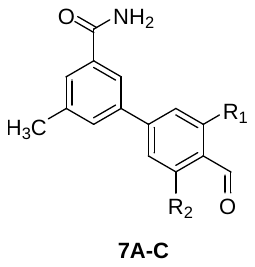

A 50-mL round-bottom flask was charged with 3-bromo-5-methylbenzamide (2.50 mmol, 1.0 eq), a commercially available formylphenyl boronic acid (2.75 mmol, 1.1 eq), and sodium bicarbonate (7.50 mmol, 3.0 eq). This was dissolved in a mixture of toluene, ethanol, and water (9.0 mL, 4.5 mL, and 5.0 mL). This mixture was evacuated and backfilled with nitrogen three times. [1,1′-Bis(diphenylphosphino)ferrocene]dichloropalladium(II) (36.6 mg, 2.0 mol %) was added and the resulting reaction mixture was heated to 80 °C for 18 hr. The reaction mixture was cooled to ambient temperature and quenched with water. The product was extracted with ethyl acetate (3 x 15 mL) and the organic layer was dried over sodium sulfate, filtered, and concentrated. The resulting solid was triturated with ether to yield a yellow solid.

**7A**: R_1_ = R_2_ = F ^1^H-NMR (DMSO) δ 10.25 (s, 1H), 8.11 (s, 2H), 7.87 (s, 1H), 7.81 (s, 1H), 7.76 (s, 1H), 7.73 (s, 1H), 7.48 (s, 1H), 2.44 (s, 3H). ^13^C-NMR (DMSO) δ 185.0, 167.8, 164.4 (d, J = 7.1 Hz), 161.8 (d, J = 6.9 Hz), 148.5, 139.3, 136.7, 135.5, 130.9, 130.1, 123.6, 111.3, 111.0, 21.4. HRMS (ESI) *m/z* [M + H] ^+^: 276.0836 (calculated), 276.0859(found).

**7B**: R_1_ = F, R_2_ = H, 83% yellow solid: ^1^H NMR (400 MHz, DMSO) δ 10.07 (s, 1H), 8.10-8.09 (m,2H), 7.94 (t, 1H, J = 7.88 Hz), 7.85-7.84 (m, 1H), 7.80-7.79 (m, 3H), 2.45 (s, 3H). ^13^C -NMR (DMSO) δ187.9, 168.0, 165.5, 162.9, 148.3, 148.2, 139.2, 137.8 (d, J = 1.4 Hz), 135.6, 131.0, 130.4, 129.5, 123.8, 123.2, 123.1, 115.2, 115.0, 21.4. HRMS (ESI) *m/z* [M + H] ^+^: 257.0484 (calculated), 257.0852 (found).

**7C**: R_1_ = R_2_ = H, 63% yellow solid: ^1^H NMR (400 MHz, DMSO) δ 10.03 (s, 1H), 8.09-7.97 (m,6H), 7.77 (s, 2H), 7.43 (s, 1H), 2.45 (s, 3H). ^13^C-NMR (DMSO) 193.2, 168.2, 145.9, 139.2, 139.0, 135.7, 135.6, 130.9, 130.6, 128.8, 128.4, 128.0, 123.8, 49.2, 21.5. HRMS (ESI) *m/z* [M + H] ^+^: 240.1020 (calculated), 240.1019 (found).

Synthesis of Compound **8A-10D**

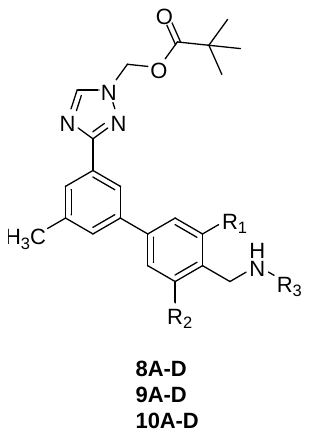

A 20-mL vial was charged with **11A-11C** (0.800 mmol, 1.0 eq) and a commercially available amine (1.20 mmol, 1.5 eq). This was dissolved in 1:1 methylene chloride and methanol (9.60 mL). Acetic acid (480 μL) was added, and the resulting mixture was stirred at ambient for 1 hr. Subsequently, sodium cyanoborohydride (2.40 mmol, 3.0 eq) was added, and the mixture was stirred for 18 hr at ambient temperature. The mixture was concentrated, redissolved in methylene chloride, and washed with saturated brine. The organic layer was dried with sodium sulfate, filtered, and concentrated. The residue was purified on silica with a gradient CH_2_Cl_2_ 🡪 10% MeOH/CH_2_Cl_2_. Product containing fractions were combined and concentrated to yield a clear residue.

**8A:** R_1_ = R_2_ = F, R_3_ = 2-aminoindane 23% white solid: ^1^H-NMR (CD_2_Cl_2_) δ 8.41 (s, 1H), 8.15 (s, 1H), 8.01 (s, 1H), 7.43 (S, 1H), 7.26-7.21 (m, 4H), 7.17-7.15 (m, 2H), 6.12 (s, 2H), 4.04 (s, 2H), 3.68 (q, 1H, J = 6.73 Hz), 3.22 (dd, 2H, J = 8.56 Hz, 7.08 Hz), 2.87 (dd, 2H, J = 9.32 Hz, J = 6.28 Hz), 2.50 (s, 3H), 1.23 (s, 9H). ^13^C-NMR (CD_2_Cl_2_) δ 178.0, 163.2 (d, J = 9.6 Hz), 162.8, 160.7 (d, J = 9.7 Hz), 146.2, 141.6, 139.3, 139.1, 131.1, 128.8, 127.1, 126.4, 124.7, 122.3, 110.1, 109.8, 69.4, 58.5, 39.9, 39.0, 38.8, 26.8, 21.4. HRMS (ESI) *m/z* [M + H] ^+^: 531.2567 (calculated), 531.2571 (found).

**8B**: R_1_ = R_2_ = F, R_3_ = cyclohexane 29% white solid: ^1^H-NMR (CD_2_Cl_2_) δ 8.42 (s, 1H), 8.16 (s, 1H), 8.00 (s, 1H), 7.46 (s, 1H), 7.24 (d, 2H, J = 8.4 Hz), 6.13 (s, 2H), 3.96 (s, 2H), 2.50 (s, 3H), 2.23 (s, 1H), 1.95 (d, 2H, J = 11.3 Hz), 1.77 (d, 2H, J = 9.2 Hz), 1.65 (d, 1H, J = 10.9 Hz), 1.24 (s, 9H). ^13^C-NMR (CD_2_Cl_2_) δ 178.0, 163.2 (d, J = 9.8 Hz), 162.8, 160.71 (d, J = 9.9 Hz), 146.2, 142.1, 139.19, 131.0, 128.8, 127.1, 122.3, 110.0, 109.8, 69.4, 55.6, 38.8, 37.7, 33.2, 26.8, 26.1, 24.9, 21.4.

HRMS (ESI) *m/z* [M + H] ^+^: 497.2723 (calculated), 497.2727 (found).

**8C**: R_1_ = R_2_ = F, R_3_ = methylenecyclohexane 34% white solid: ^1^H-NMR (CD_2_Cl_2_) δ 8.42 (s, 1H), 8.17 (s,1H), 8.01 (s, 1H), 7.47 (s, 1H), 7.25 (d, 2H, J = 8.5 Hz), 6.13 (s, 2H), 3.91 (s, 2H), 2.51-2.48 (m, 5H), 1.80-1.72 (m, 7H), 1.51-1.44 (m, 1H), 1.32-1.26 (m, 2H), 1.24 (s, 9H), 0.99-0.90 (m, 2H). ^13^C-NMR (CD_2_Cl_2_) δ 178.0, 163.2 (d, J = 9.8 Hz), 162.8, 160.7 (d, J = 9.8 Hz), 146.2, 139.2, 131.0, 128.8, 127.1, 122.3, 114.7, 110.0, 109.7, 69.4, 60.4, 55.6, 40.8, 38.8, 37.9, 31.4, 26.8, 26.1, 21.4. HRMS (ESI) *m/z* [M + H] ^+^: 511.2880 (calculated), 511.2888 (found).

**8D:** R_1_ = R_2_ = F, R_3_ = 4,4-dimethylcyclohexane 31% white solid: ^1^H-NMR (CD_2_Cl_2_) δ 8.40 (s, 1H), 8.13 (s, 1H), 7.99 (s, 1H), 7.42 (s, 1H), 7.21 (d, 2H, J = 8.49 Hz), 6.11 (s, 2H), 3.98 (s, 2H), 2.98 (br s, 1H), 2.49 (s, 3H), 2.45-2.39 (m, 1H), 2.05 (s, 1H), 1.77 (d, 2H, J = 10.31 Hz), 1.43-1.37 (m, 4H), 1.21 (s, 9H), 0.92 (s, 3H), 0.91 (s, 3H). ^13^C-NMR (CD_2_Cl_2_) δ 177.9, 163.2, 163.1, 162.8, 160.8, 160.6, 146.2, 142.2, 142.1, 142.0, 139.2, 131.0, 128.8, 127.1, 122.3, 110.1, 109.8, 69.4, 60.3, 55.8, 38.8, 37.7, 32.1, 30.1, 28.8, 26.8, 24.8, 21.4, 14.2. HRMS (ESI) *m/z* [M + H] ^+^: 525.3036 (calculated), 525.3029 (found).

**9A:** R_1_ = F, R_2_ = H: 29% clear residue: ^1^H-NMR (CD_2_Cl_2_) δ 8.41 (s, 1H), 8.20 (s, 1H), 8.00 (s, 1H), 7.51 (q, 3H, J = 6.88 Hz), 7.42 (d, 1H, J=11.48 Hz), 7.26-7.24 (m, 2H), 7.20-7.16 (m, 2H), 6.12 (s, 2H), 3.98 (s, 2H), 3.72 (s, 1H), 3.22 (dd, 2H, J = 6.96 Hz, J = 8.72 Hz), 2.86 (dd, 2H, J = 6.00 Hz, J = 9.64 Hz), 1.24 (s, 9H). ^13^C-NMR (CD_2_Cl_2_) δ 177.7, 162.8, 162.6, 160.2, 146.1, 141.9, 141.6, 141.54, 139.9, 139.1, 131.2, 130.7, 130.6, 128.9, 126.7, 126.6, 126.4, 126.2, 124.5, 122.64, 122.63, 122.10, 113.8, 113.5, 69.5, 59.0, 45.21, 45.19, 40.0, 38.7, 26.6, 21.2, 14.0. HRMS (ESI) *m/z* [M + H] ^+^: 513.2661 (calculated), 513.2659 (found).

**9B:** R_1_ = F, R_2_ = H, R_3_ = cyclohexane 31% clear residue: ^1^H-NMR (CD_2_Cl_2_) δ 8.41 (s, 1H), 8.19 (s, 1H), 7.99 (s, 1H), 7.51-7.48 (m, 3H), 7.40 (d, 1H, J = 11.48 Hz), 6.12 (s, 2H), 3.93 (s, 2H), 2.52 (s, 3H), 1.96 (d, 2H, J = 12.29 Hz), 1.80-1.76 (m, 2H), 1.66-1.64 (m, 2H), 1.37-1.11 (m, 10H). ^13^C-NMR (CD_2_Cl_2_) δ 177.7, 162.8, 162.6, 160.2, 146.2, 141.3, 141.2, 140.1, 140.0, 139.1, 131.2, 130.6, 130.5, 128.8, 127.3, 127.1, 126.4, 122.6, 122.5, 122.1, 113.7, 113.5, 69.5, 56.0, 43.83, 43.81, 38.7, 33.5, 26.6, 26.3, 25.0, 21.2. HRMS (ESI) *m/z* [M + H] ^+^: 479.2817 (calculated), 479.2808 (found).

**9C:** R_1_ = F, R_2_ = H, R_3_ = methylenecyclohexane 25% clear residue: ^1^H-NMR (CD_2_Cl_2_) δ 8.41 (s, 1H), 8.19 (s, 1H), 7.99 (s, 1H), 7.51-7.48 (m, 3H), 7.40 (d, 1H, J = 11.38 Hz), 6.12 (s, 2H), 3.88 (s, 2H), 2.52 (s, 3H), 2.50 (s, 2H), 1.83-1.67 (m, 6H), 1.59-1.49 (m, 2H), 1.35-1.20 (m, 10H), 1.01-0.92 (m, 2H). ^13^C-NMR (CD_2_Cl_2_) δ 177.7, 162.8, 162.65, 160.2, 146.1, 141.39, 141.31, 140.05, 140,03, 139.1, 131.1, 130.57, 130.51, 128.7, 126.9, 126.8, 126.3, 122.55, 122.53, 122.1, 113.7, 113.4, 69.5, 60.2, 56.1, 47.0, 46.9, 38.7, 38.1, 31.4, 26.7, 26.5, 26.1, 21.2, 14.0. HRMS (ESI) *m/z* [M + H] ^+^: 493.2974 (calculated), 493.2979 (found).

**9D:** R_1_ = F, R_2_ = H, R_3_ = 4,4-dimethylcyclohexane 29% clear residue: ^1^H-NMR (CD_2_Cl_2_) δ 8.41 (s, 1H), 8.19 (s, 1H), 7.99 (s, 1H), 7.51-7.47 (m, 3H), 7.40 (d, 1H, J = 11.51 Hz), 6.12 (s, 2H), 3.92 (s, 2H), 2.51 (s, 3H), 2.04 (s, 1H), 1.81-1.77 (m, 2H), 1.47-1.35 (m, 7H), 1.26-1,23 (m, 10H), 0.96 (s, 3H), 0.95 (s, 3H). ^13^C-NMR (CD_2_Cl_2_) δ 177.7, 170.7, 162.8, 162.6, 160.1, 146.1, 141.3, 141.2, 140.0, 139.1, 131.1, 130.6, 130.5, 128.7, 127.3, 127.1, 126.3, 122.6, 122.5, 122.0, 113.7, 113.5, 69.5, 60.2, 56.2, 44.0, 43.9, 38.7, 37.6, 31.7, 29.9, 29.2, 26.5, 24.8, 21.2, 20.7, 14.0. HRMS (ESI) *m/z* [M + H] ^+^: 507.3130 (calculated), 507.3127 (found).

**10A:** R_1_ = R_2_ = H, R_3_ = 2-aminoindane 29% clear residue: ^1^H-NMR (CD_2_Cl_2_) δ 8.42 (s, 1H), 8.20 (s, 1H), 7.98 (s, 1H), 7.69 (d, 2H, J = 8.19 Hz), 7.52-7.50 (m, 3H), 7.25-7.23 (m, 2H), 7.18-7.15 (m, 2H), 6.12 (s, 2H), 3.98 (s, 2H), 3.77 (s, 1H), 3.24 (dd, 2H, J = 7.24 Hz, J = 8.40 Hz), 2.96 (dd, 2H, J = 6.56 Hz, J = 9.04 Hz), 2.52 (s, 3H), 1.24 (s, 9H). ^13^C-NMR (CD_2_Cl_2_) δ 177.7, 162.9, 146.1, 141.4, 141.0, 139.7, 139.0, 138.1, 131.0, 129.0, 128.9, 127.1, 126.4, 125.9, 124.5, 122.1, 69.5, 60.2, 58.6, 51.1, 39.1, 38.7, 26.5, 22.1, 21.2, 20.7, 14.0. HRMS (ESI) *m/z* [M + H] ^+^: 495.2760 (calculated), 495.2753 (found).

**10B**: R_1_ = R_2_ = H, R_3_ = cyclohexane ^1^H-NMR (CD_2_Cl_2_) ) δ 8.42 (s, 1H), 8.22 (s, 1H), 7.98 (s, 1H), 7.68 (d, 2H, J =7.87), 7.53 (s. 1H), 7.47 (d, 2H, J = 7.85 Hz), 6.13 (s, 2H), 3.90 (s, 2H), 2.85 (br s, 1H), 2.52 (s, 3H), 1.99 (d, 2H, J = 11.40 Hz), 1.79 (d, 2HJ = 6.15 Hz), 1.76 (d, 2H, J = 9.9 Hz), 1.25 (s, 9H). ^13^C-NMR (CD_2_Cl_2_) δ 177.7, 162.9, 146.2, 141.3, 140.4, 139.2, 138.9, 131.0, 128.9, 128.6, 126.9, 125.8, 122.2, 69.5, 56.2, 50.3, 38.7, 33.4, 26.6, 26.3, 25.0, 21.3. HRMS (ESI) *m/z* [M + H] ^+^: 479.2817 (calculated), 479.2825 (found).

**10C:** R_1_ = R_2_ = H, R_3_ = methylenecyclohexane 34% clear residue: ^1^H-NMR (CD_2_Cl_2_) δ 8.41 (s, 1H), 8.18 (s, 1H), 7.96 (s, 1H), 7.67 (d, 2H, J = 8.11 Hz), 7.48 (d, 3H, J = 7.93 Hz), 6.12 (s, 2H), 3.90 (s, 2H), 2.00 (s, 2H), 1.83-1.71 (m, 6H), 1.62-1.59 (m, 1H), 1.36-1.26 (m, 3H), 1.23 (s, 9H), 0.99-0.95 (m, 3H). ^13^C-NMR (CD_2_Cl_2_) δ 177.7, 162.9, 146.1, 141.0, 139.8, 139.0, 137.4, 131.0, 129.1, 128.9, 128.1, 126.0, 122.2, 69.5, 54.8, 52.5, 38.7, 37.0, 31.2, 26.6, 26.5, 25.9, 21.2. HRMS (ESI) *m/z* [M + H] ^+^: 475.3073 (calculated), 475.3073 (found).

**10D:** 147D R_1_ = R_2_ = H, R_3_ = 4,4-dimethylcyclohexane, 33% clear residue: ^1^H-NMR (CD_2_Cl_2_) δ 8.40 (s, 1H), 8.15 (s, 1H), 7.95 (s, 1H), 7.71 (s, 4H), 7.42 (s, 1H), 6.10 (s, 2H), 4.05 (s, 2H), 2.83-2.77 (m, 1H), 2.61 (s, 1H), 2.02 (s, 1H), 1.98 (br s, 3H), 1.89-1.80 (m, 2H), 1.48 (d, 2H, J = 13.28 Hz), 1.22 (s, 9H), 0.97 (s, 3H), 0.88 (s, 3H). 13C-NMR (CD_2_Cl_2_) δ 177.7, 162.7, 146.2, 141.3, 140.5, 139.0, 131.0, 130.9, 130.6, 128.9, 127.4, 126.3, 122.2, 69.5, 55.5, 47.1, 40.8, 38.7, 37.0, 31.5, 29.5, 26.6, 24.9, 23.8, 21.2. HRMS (ESI) *m/z* [M + H] ^+^: 489.3230 (calculated), 489.3252 (found).

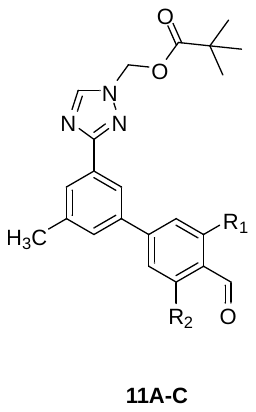

A 50-mL round-bottom flask was charged with **13** (3.90 mmol, 1.0 eq), a commercially available formylphenyl boronic acid (4.30 mmol, 1.1 eq), and sodium bicarbonate (11.72 mmol, 3.0 eq). This was dissolved in a mixture of toluene, ethanol, and water (13.0 mL, 6.5 mL, and 7.0 mL). This mixture was evacuated and backfilled with nitrogen three times. [1,1′-Bis(diphenylphosphino)ferrocene]dichloropalladium(II) (0.0781 mmol, 2.0 mol %) was added and the resulting reaction mixture was heated to 80 C for 18 hr. The reaction mixture was cooled to ambient temperature and quenched with water. The product was extracted with ethyl acetate (3 x 15 mL) and the organic layer was dried over sodium sulfate, filtered, and concentrated. The resulting solid was triturated with ether to yield a yellow solid.

**11A:** R_1_ = R_2_ = F, 40% white solid: ^1^H-NMR (DMSO) δ 10.23 (s, 1H), 8.80 (s, 1H), 8.12 (s, 1H), 7.94 (s, 1H), 7.73 (s, 1H), 7.65 (d, J=10.51 Hz, 2H), 6.18 (s, 2H), 2.45 (s, 3H), 1.12 (s, 9H).

^13^C-NMR (DMSO) δ 185.02, 177.17, 164.37, 161.95, 161.83, 148.74, 147.90, 139.85, 137.62, 131.68, 129.57, 128.20, 122.23, 112.88, 111.36, 111.11, 70.42, 38.74, 26.99, 21.38. HRMS (ESI) m/z [M + H] ^+^: 414.1624 (calculated), 414.1631 (found).

**11B:** R_1_ = H, R_2_ = F, 75% white sold: 1H-NMR (DMSO) d 10.26 (s, 1H), 8.82 (s, 1H), 8.15 (s, 1H), 7.97-7,93 (m, 2H), 7.80-7.75 (m, 2H), 7.72 (s, 1H), 6.21 (s, 2H), 2.46 (s, 3H), 1.14 (s, 9H). 13C-NMR (DMSO) d 187.88, 177.16, 165.50, 162.95, 162.04, 148.48, 148.39, 147.88, 139.77, 138.60, 131.63, 130.46, 129.53, 127.67, 123.74, 123.18, 123.10, 122.20¸ 115.20, 115.00, 70.41, 38.73, 26.98, 21.41. HRMS (ESI) m/z [M + H] ^+^: 396.1723 (calculated), 396.1747 (found).

**11C:** R_1_ = R_2_ = H, 86% yellow solid: ^1^H-NMR (DMSO) δ 10.08 (s, 1H), 8.81 (s, 1H), 8.15 (s, 1H), 8.04 (d, J=8.22 Hz, 2H), 7.97-7.93 (m, 3H), 6.21 (s, 2H), 2.48 (s, 3H), 1.14 (s, 9H). ^13^C-NMR (DMSO) δ 193.21, 177.16, 162.15, 147.87, 145.94, 139.88, 139.69, 135.75, 131.58, 130.69, 129.47, 127.91, 127.10, 122.20, 70.41, 38.74, 26.99, 21.48. HRMS (ESI) *m/z* [M + H] ^+^: 378.1813 (calculated), 378.1818 (found).

Synthesis of Compound **12**

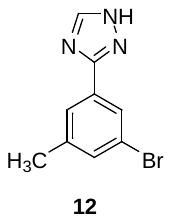

A 15-mL round bottom flask was charged with 3-bromo-5-methylbenzamide (4.67 mmol, 1.0 eq) and N,N-Dimethylformamide dimethyl acetal (2.50 mL). This mixture heated to 80 C for 1 hr and was subsequently cooled to ambient temperature. The mixture was poured into crushed ice (30 mL) and stirred for 1 hr. The precipitated solid was collected by filtration and dried. The intermediate N-acyl formimadine was dissolved in acetic acid (6.15 mL) and hydrazine monohydrate (4.67 mmol, 1.0 eq) was added. This heated to 90 C for 1 hr. The reaction was cooled, and dumped into crushed ice (30 mL) and stirred for 1 hr. This mixture was quenched with saturated sodium carbonate and the product was extracted with ethyl acetate (3 x 20 mL). The combined organic extracts were dried over sodium sulfate, filtered, and concentrated. The resulting white solid was used without further purification.

**12:** 69% white solid. ^1^H-NMR (DMSO) δ 14.26 (br s, 1H), 8.51 (s, 1H), 7.96 (s, 1H), 7.85 (s, 1H), 7.47 (s, 1H), 2.38 (s, 1H). ^13^C-NMR (DMSO) δ 164.72, 141.43, 136.83, 132.72, 128.58, 125.96, 125.92, 122.32, 21.11. HRMS (ESI) *m/z* [M + H] ^+^: 237.9993 (calculated), 237.9980 (found).

Synthesis of Compound **13**

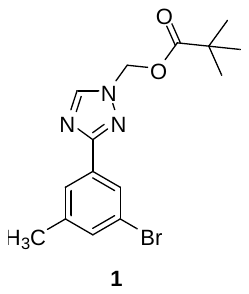

A 15-mL round bottom flask was charged with **12** (16.50 mmol, 1.0 eq), potassium carbonate (24.76 mmol, 1.50 eq) and anhydrous acetonitrile (20 mL). Subsequently, chloromethyl pivalate (24.76 mmol, 1.50 eq) was added. This mixture heated to 80 C for 1.5 hr and was subsequently cooled to ambient temperature. The mixture was poured into crushed ice (30 mL) and stirred for 1 hr. The product was extracted with methylene chloride (3 x 15 mL). The combined organic extracts were dried over sodium sulfate, filtered, and concentrated. The resulting residue was purified on silica with a gradient 10% EtOAc/hexanes 🡪100% EtOAc to yield a white solid.

**13:** 72% white solid**:** ^1^H-NMR (DMSO) δ 8.81 (s, 1H), 7.93 (s, 1H), 7.84 (s, 1H), 7.52 (s, 1H), 6.19 (s, 2H), 2.39 (s, 3H), 1.14 (s, 9H). ^13^C-NMR (DMSO) δ 177.13, 161.07, 148.02, 141.62, 133.17, 132.75, 126.02, 125.97, 122.37, 70.38, 38.74, 26.98, 21.07. HRMS (ESI) *m/z* [M + H] ^+^: 352.0659 (calculated), 352.0661 (found).

Synthesis of Compound **14A-14B**

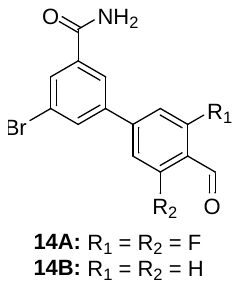

A 50-mL round-bottom flask was charged with 3,5-dbromo-benzamide (3.59 mmol, 1.0 eq), 4-formylphenylboronic acid (3.59 mmol, 1.0 eq), and sodium bicarbonate (10.76 mmol, 3.0 eq). This was dissolved in a mixture of toluene, ethanol, and water (12.0 mL, 6.0 mL, and 8.0 mL). This mixture was evacuated and backfilled with nitrogen three times. [1,1′-Bis(diphenylphosphino)ferrocene]dichloropalladium(II) (0.0718 mmol, 2.0 mol %) was added and the resulting reaction mixture was heated to 80 C for 18 hr. The reaction mixture was cooled to ambient temperature and quenched with water. The product was extracted with ethyl acetate (3 x 15 m) and the organic layer was dried over sodium sulfate, filtered, concentrated and dry loaded onto a silica column. The crude material was purified with a gradient 10% EtOAc/CH_2_Cl_2_ 🡪100% EtOAc to yield a white solid.

**14A**: R_1_ = R_2_ = F 34% white solid: ^1^H-NMR (DMSO) δ 10.25 (s, 1H), 8.28 (s, 1H), 8.24-8.23 (m, 2H), 8.11 (s, 1H), 7.80 (d, 2H, J = 10.5 Hz), 7.67 (s, 1H). ^13^C-NMR (DMSO) δ 185.0, 166.3, 164.3 (d, J = 7.2 Hz), 161.8 (d, J = 7.2 Hz), 146.6, 139.1, 137.5, 132.8, 131.8, 125.6, 123.2, 113.4, 111.8, 111.5. HRMS (ESI) *m/z* [M + H] ^+^: 339.9780 (calculated), 339.9777 (found).

**14B:** R_1_ = R_2_ = H 36% white solid: ^1^H-NMR (DMSO) δ 10.09 (s, 1H), 8.25 (s, 2H), 8.14 (s, 1H), 8.10 (s, 1H), 8.03 (s, 4H), 7.63 (s, 1H). ^13^C-NMR (DMSO) δ 193.3, 166.5, 144.0, 141.6, 137.5, 136.2, 132.6, 130.6, 125.7, 123.0. HRMS (ESI) *m/z* [M + H] ^+^: 303.9968 (calculated), 303.9979 (found).

Synthesis of Compounds **15A-15B.**

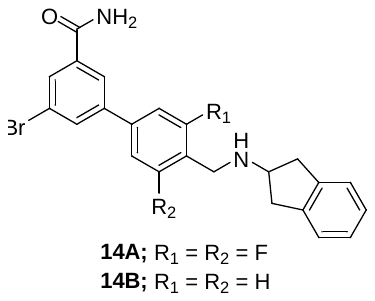

A 20-mL vial was charged with **14A or 14B** (0.329 mmol, 1.0 eq) and a 2-aminoindance (0.493 mmol, 1.5 eq). This was dissolved in 1:1 methylene chloride and methanol (4.00 mL). Acetic acid (200 μL) was added, and the resulting mixture was stirred at ambient for 1 hr. Subsequently, sodium cyanoborohydride (0.986 mmol, 3.0 eq) was added, and the mixture was stirred for 18 hr at ambient temperature. The mixture was concentrated, redissolved in methylene chloride, and washed with saturated brine. The organic layer was dried with sodium sulfate, filtered, and concentrated. The residue was purified on silica with a gradient CH_2_Cl_2_ 🡪10% MeOH/CH_2_Cl_2_. Product containing fractions were combined and concentrated to yield a clear residue.

**14A**: R_1_ = R_2_ = F 68% white solid: ^1^H-NMR (DMSO) δ 8.21 (s, 2H), 8.14 (s, 1H), 8.05 (s, 1H), 7.63-7.59 (m, 3H), 7.19-7.17 (m, 2H), 7.11-7.09 (m, 2H), 3.84 (s, 2H), 3.51 (quin, 1H, J = 6.5 Hz), 3.08 (dd, 2H, J = 7.0 Hz, 6.9 Hz), 2.72 (dd, 2H, J = 8.6 Hz, 6.1 Hz), 1.82 (s, 1H). ^13^C-NMR (DMSO) δ 166.5, 163.2 (d, J = 9.9 Hz), 160.8 (d, J = 10.0 Hz), 142.3, 140.2, 139.7, 137.4, 132.3, 130.7, 126.6, 125.1, 124.9, 123.0, 116.4, 110.7, 110.3, 58.7, 38.7. HRMS (ESI) *m/z* [M + H] ^+^: 457.0722 (calculated), 457.0719 (found).

**14B**: R_1_ = R_2_ = H 74% white solid: ^1^H-NMR (DMSO) δ 8.21 (s, 1H), 8.16 (s, 1H), 8.01 (d, 2H, J = 1.4 Hz), 7.73 (d, 2H, J = 8.2 Hz), 7.58 (s, 1H), 7.51 (d, 2H, J = 8.2 Hz), 7.20-7.18 (m, 2H), 7.13-7.10 (m, 2H), 3.87 (s, 2H), 3.59 (quin, 1H, J = 6.9 Hz), 3.10 (dd, 2H, J = 7.2 Hz, 6.8 Hz), 2.79 (dd, 2H, J = 6.6 Hz, 6.5 Hz), 1.91 (s, 1H). ^13^C-NMR (DMSO) δ 172.6, 166.7, 142.8, 142.1, 137.4, 136.9, 132.0, 129.4, 127.2, 126.7, 125.1, 124.9, 122.8, 58.9, 51.2, 21.6. HRMS (ESI) *m/z* [M + H] ^+^: 421.0911 (calculated), 421.0909 (found).

Synthesis of Compound **15A-15B**

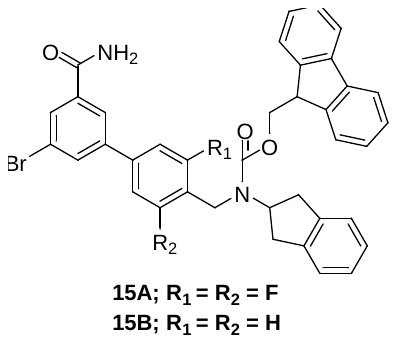

A 5-mL round bottom flask was charged with **14A or 14B** (0.155 mmol, 1.0 eq), sodium carbonate (0.310 mmol, 2.0 eq) and a 1:1 mixture of acetonitrile:water (1.50 mL). Subsequently, Fmoc-Cl (0.186 mmol, 1.20 eq) was added and the reaction mixture stirred at ambient for 1.5 hr. The reaction mixture was diluted with water (5 mL) and the product was extracted with methylene chloride (3 x 3 mL). The organic layer was dried over sodium sulfate, filtered, and concentrated. The crude residue was purified on silica with a gradient CH_2_Cl_2_ 🡪10% MeOH/CH_2_Cl_2._ Product containing fractions were combined and concentrated to yield a white crystalline solid.

**14A**: R_1_ = R_2_ = F 88% white solid HRMS (ESI) *m/z* [M + H] ^+^: 679.1403 (calculated), 679.1410 (found).

**14B**: R_1_ = R_2_ = H 84% white solid HRMS (ESI) *m/z* [M + H] ^+^: 643.1519 (calculated), 643.1518 (found).

**Table S2 Purity of Final Compounds:**

| Compound | Purity | Compound | Purity |
| --- | --- | --- | --- |
| **1A** | > 99.9% | **4A** | = 95.2% |
| **1B** | = 95.8% | **4B** | = 96.2% |
| **1C** | = 95.7% | **4C** | = 98.7% |
| **1D** | = 99.2% | **4D** | = 96.5% |
| **2A** | > 99.9% | **5A** | = 99.1% |
| **2B** | = 96.9% | **5B** | = 97.4% |
| **2C** | = 95.6% | **5C** | = 96.2% |
| **2D** | = 95.7% | **5D** | > 99.9% |
| **3A** | = 99.0% | **6A** | = 96.5% |
| **3B** | = 95.6% | **6B** | = 97.9% |
| **3C** | = 95.7% | **6C** | > 99.9% |
| **3D** | = 98.0% | **6D** | = 96.7% |

^1^H and ^13^C NMR Spectra of Final Compounds

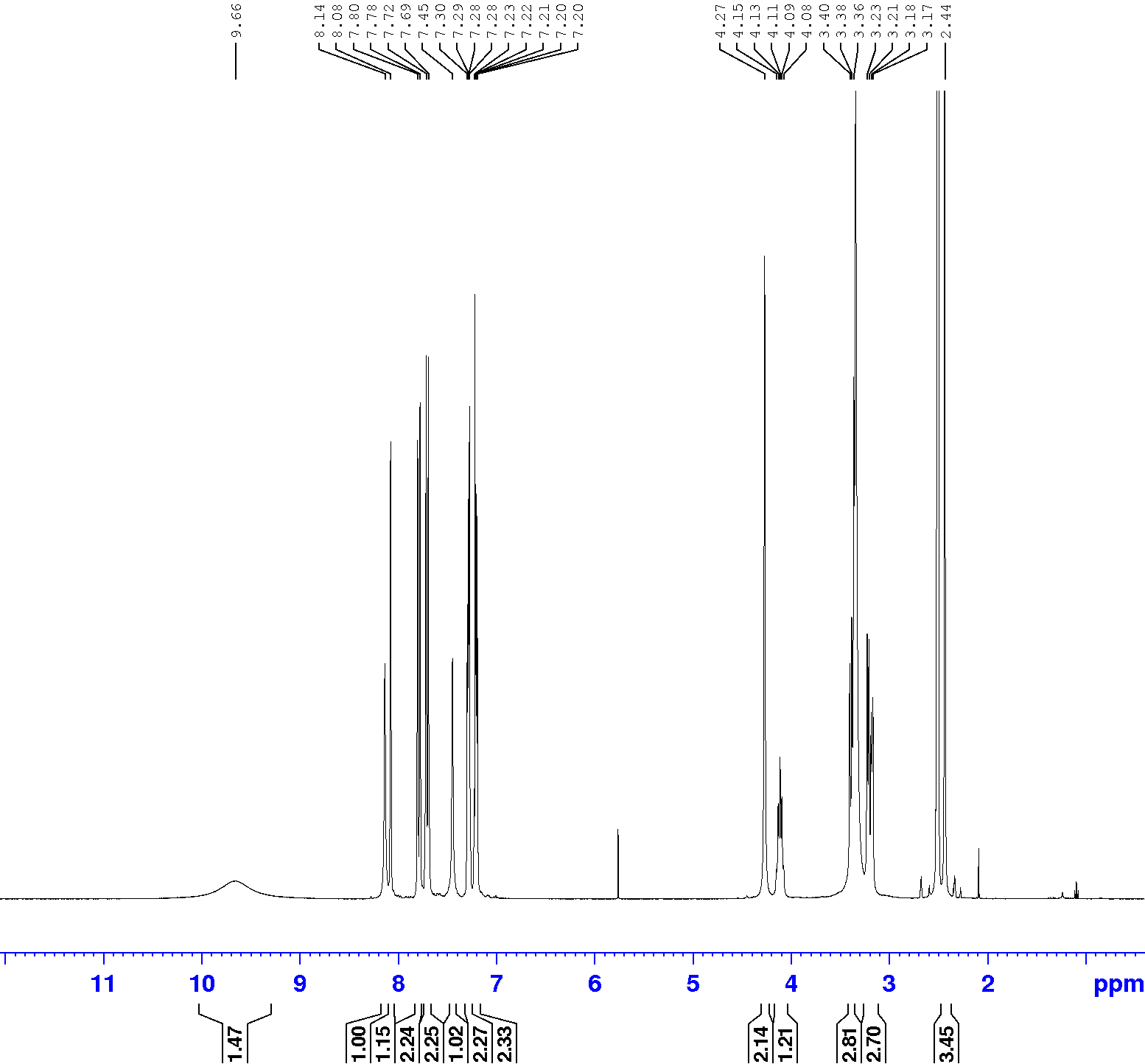

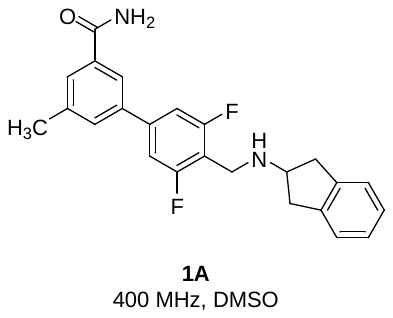

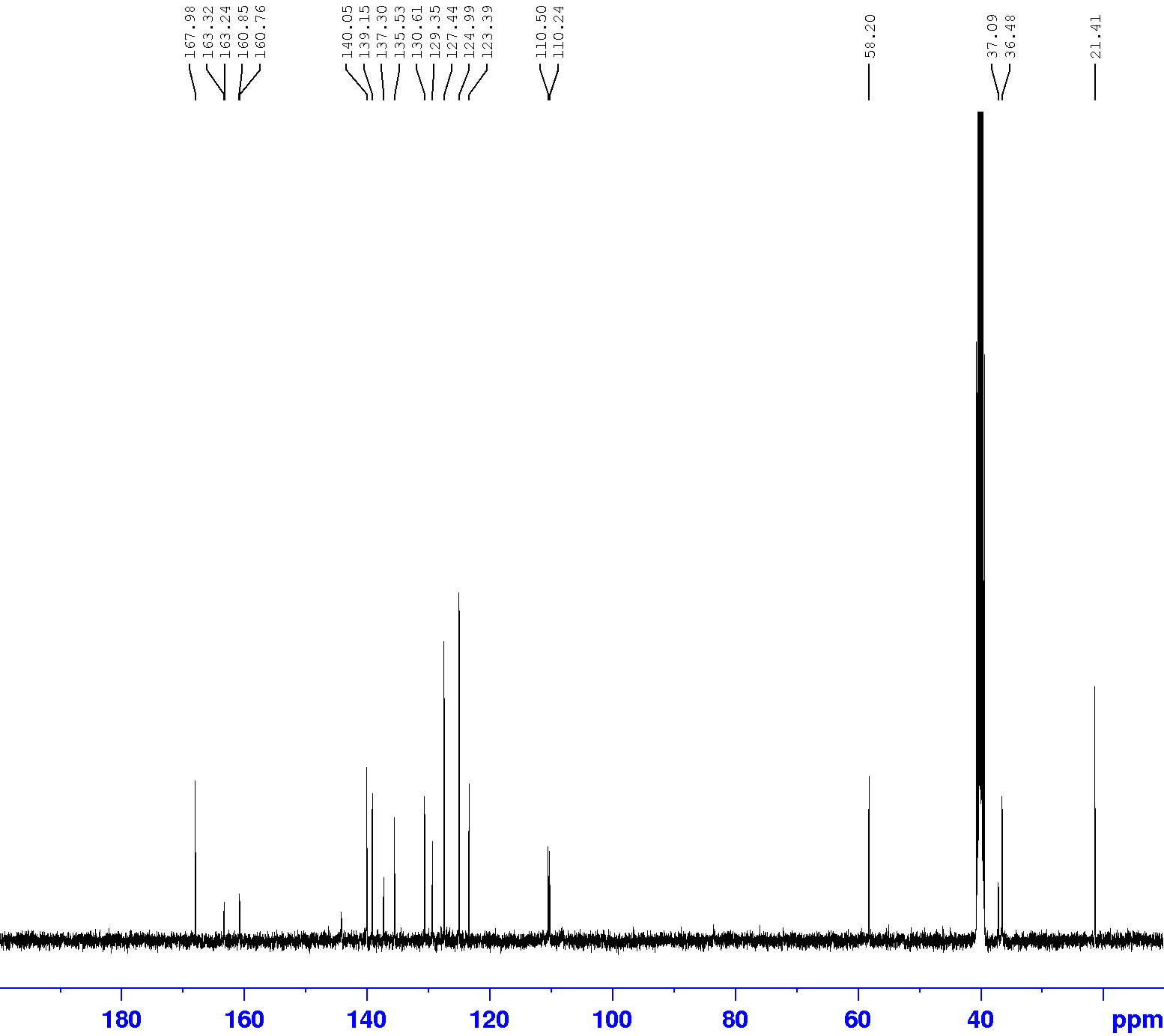

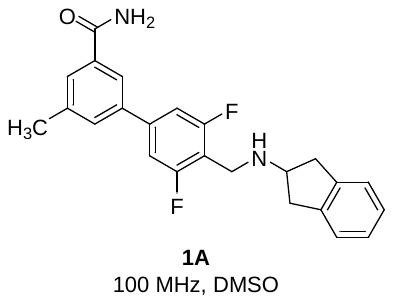

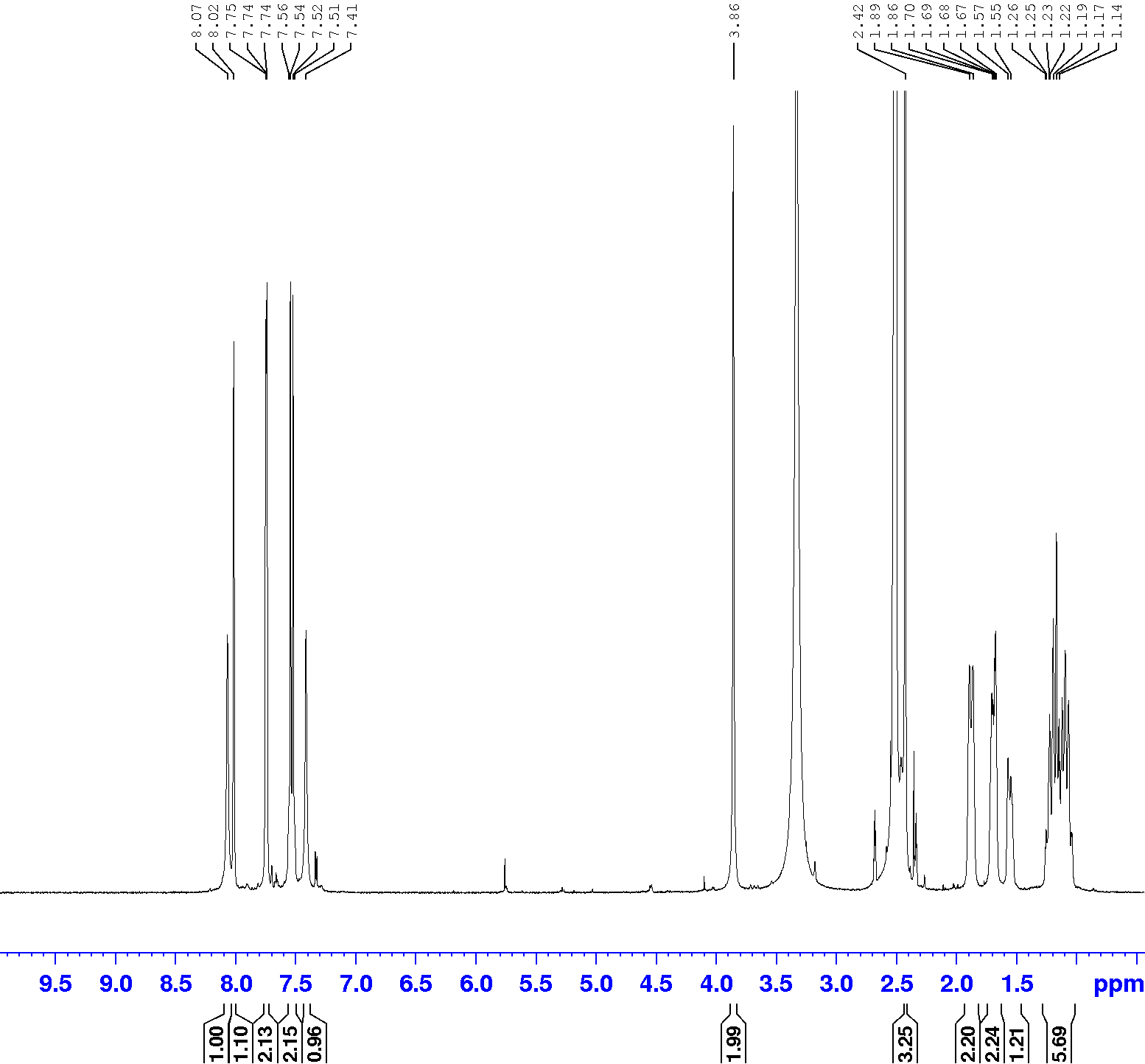

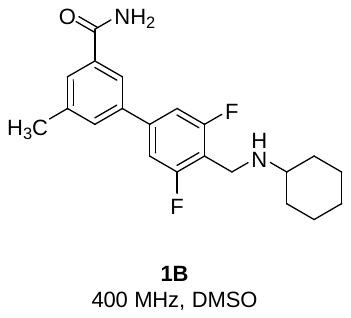

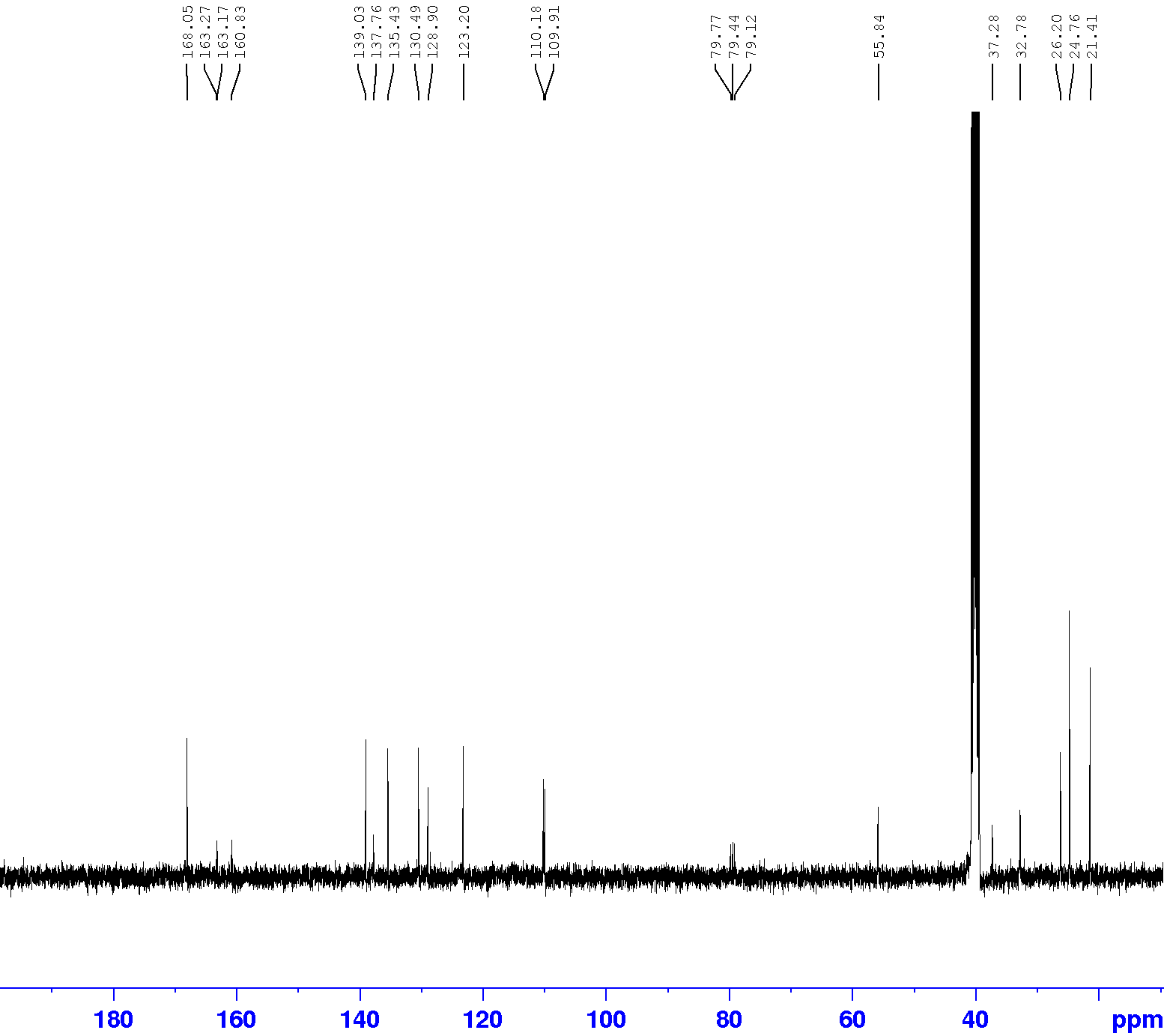

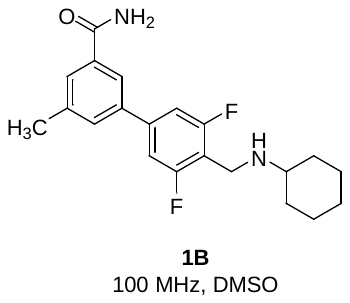

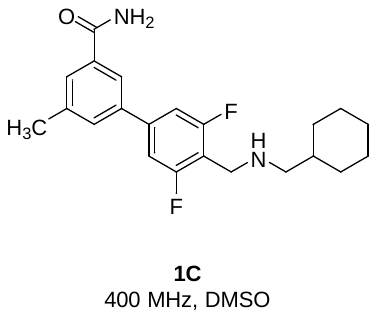

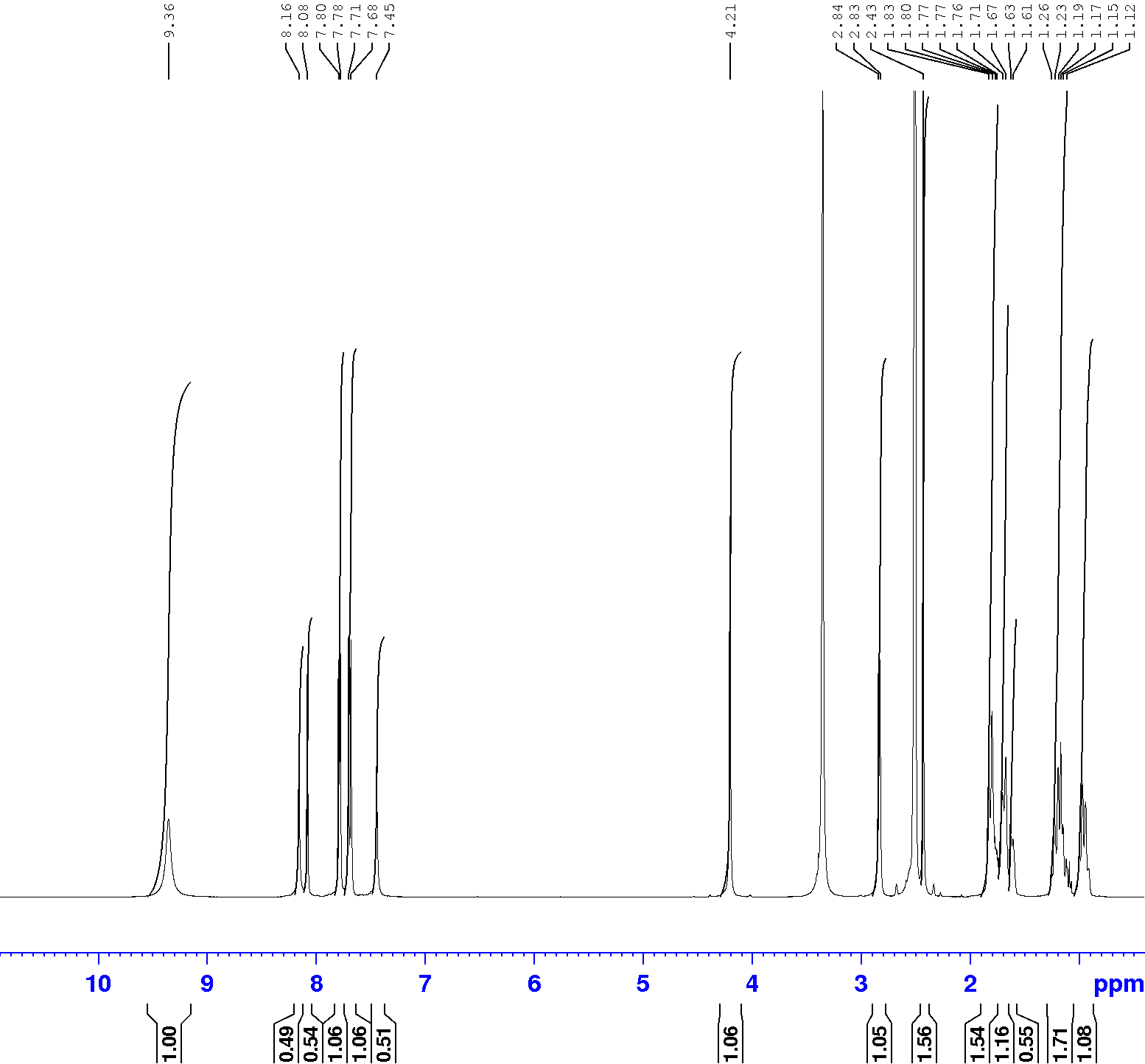

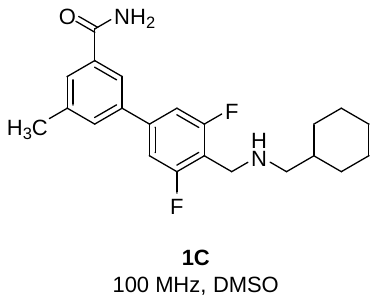

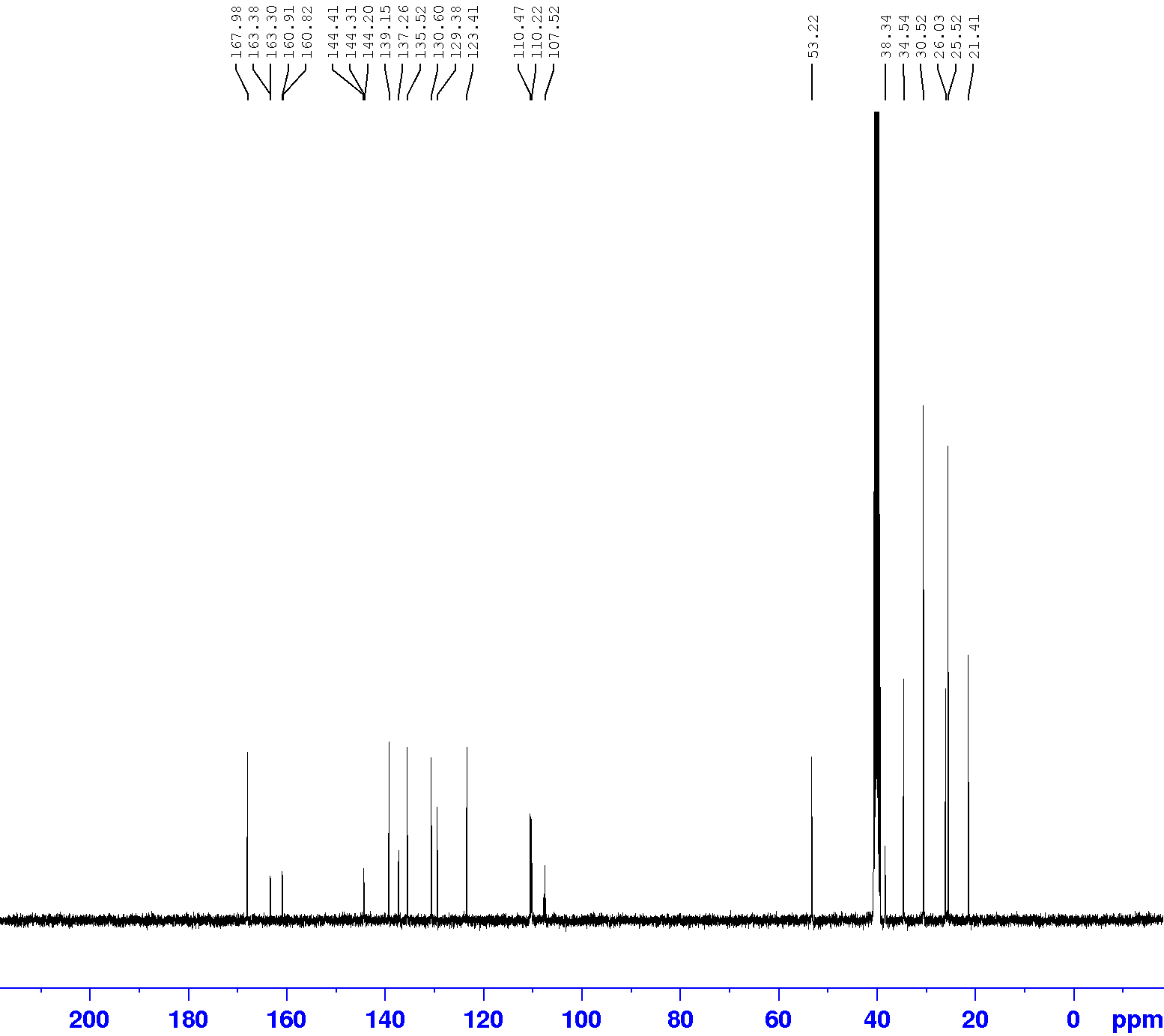

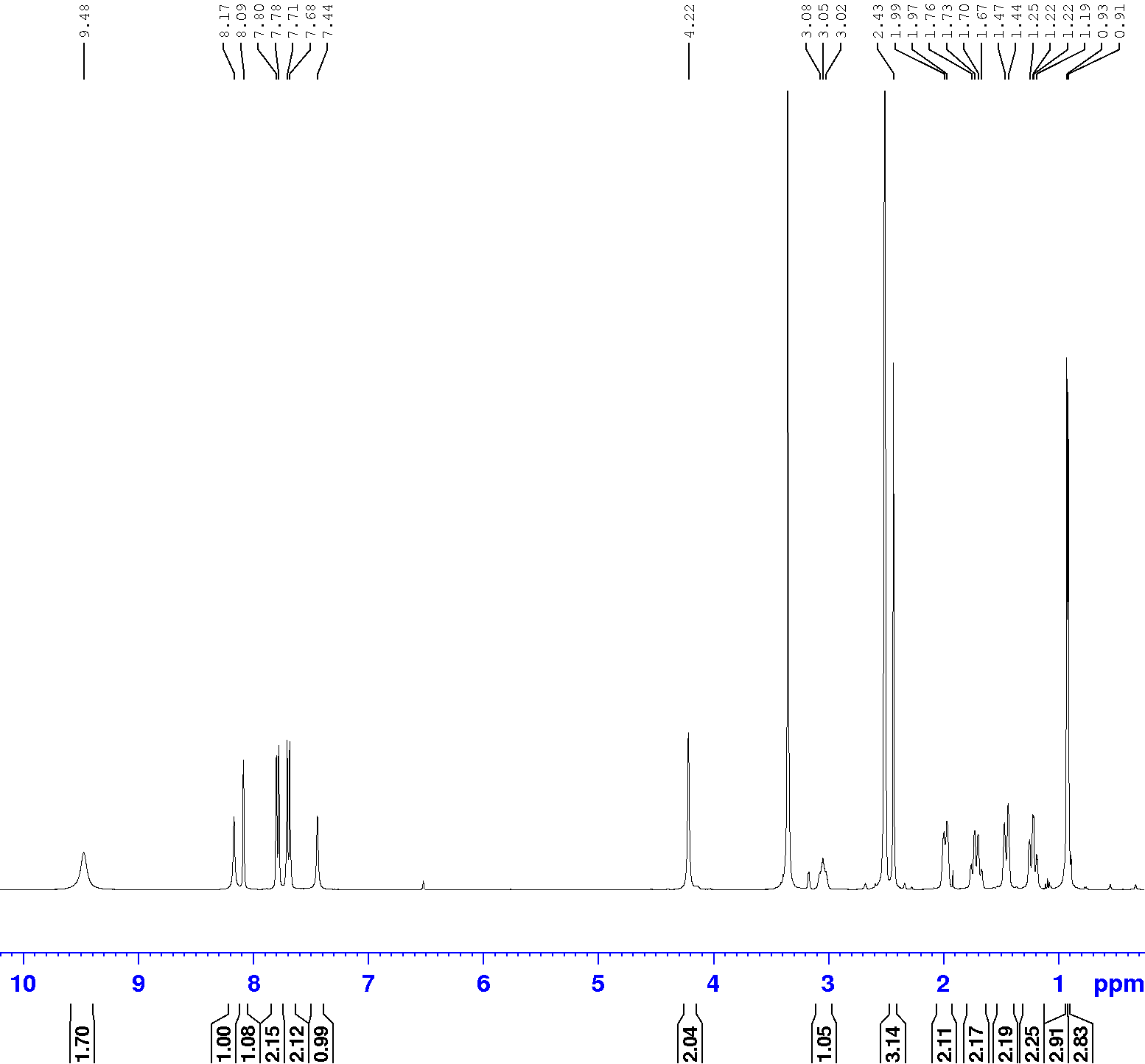

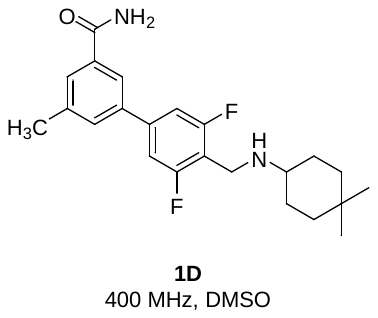

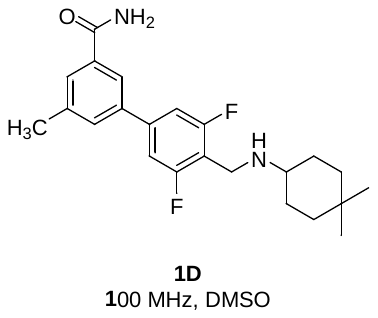

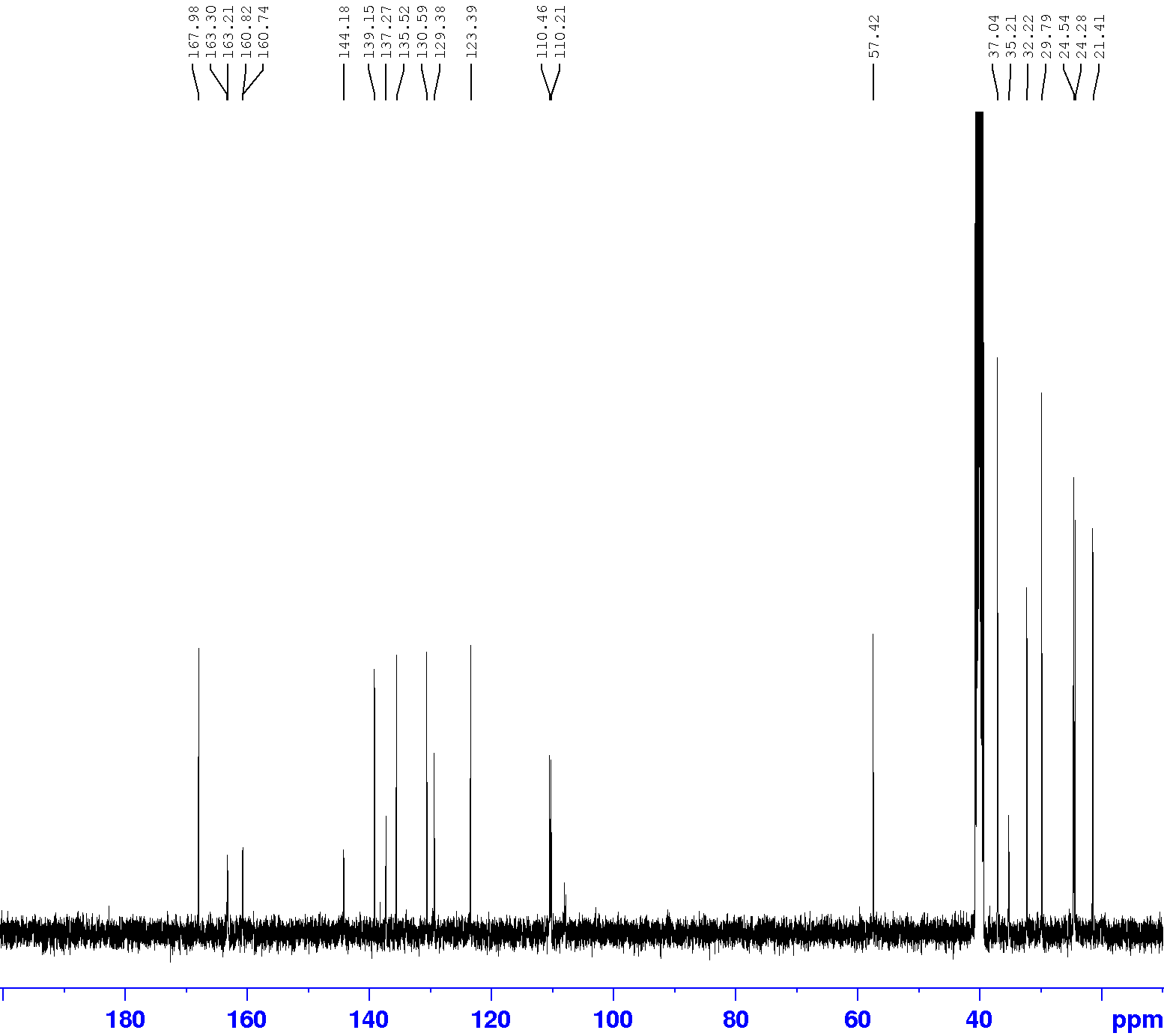

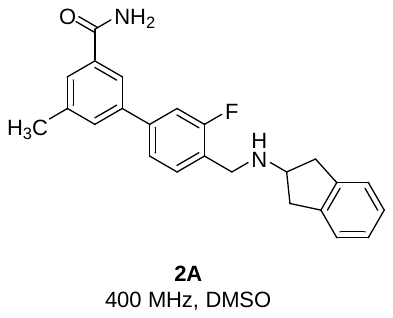

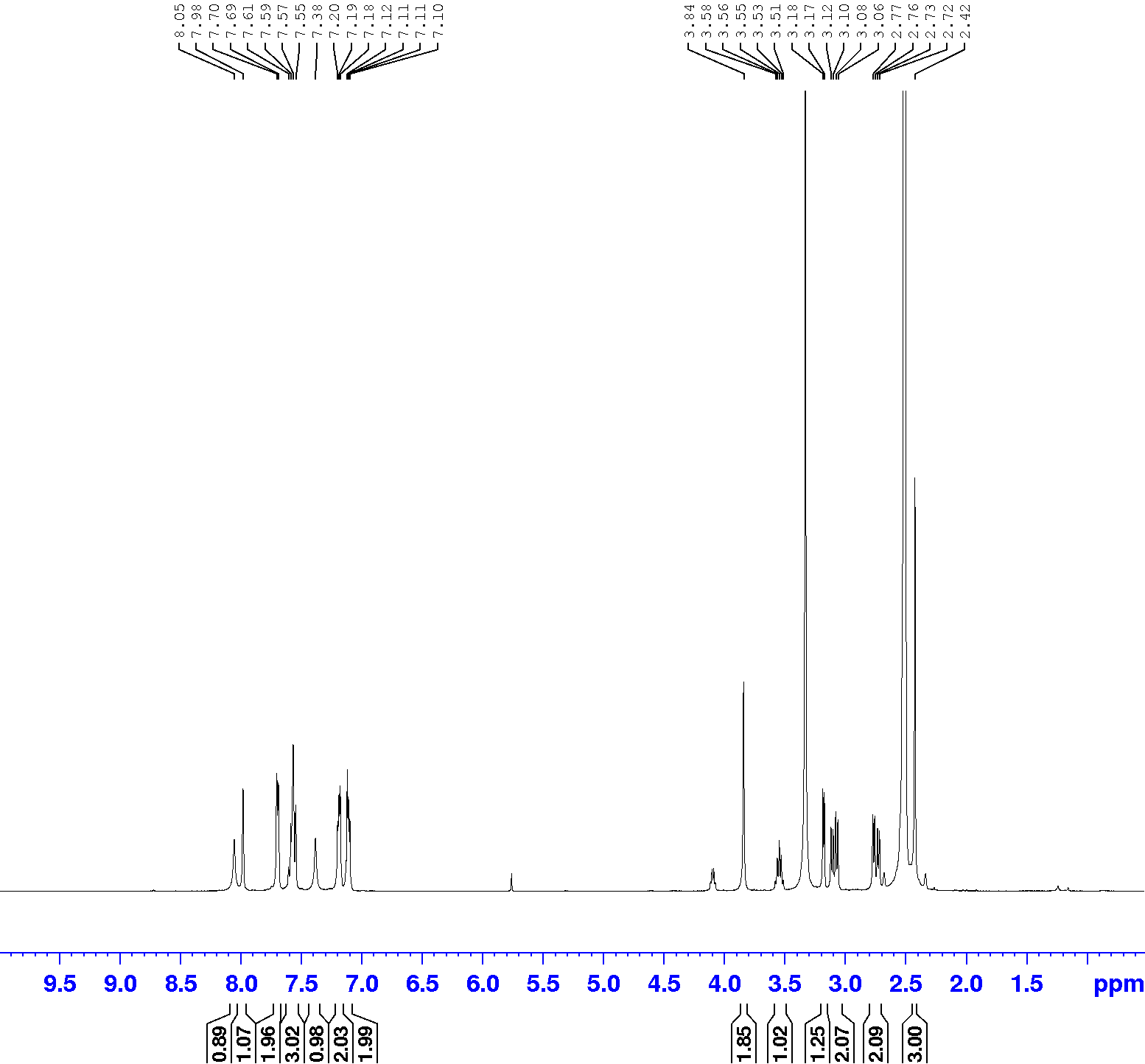

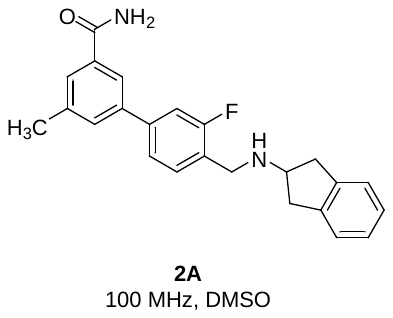

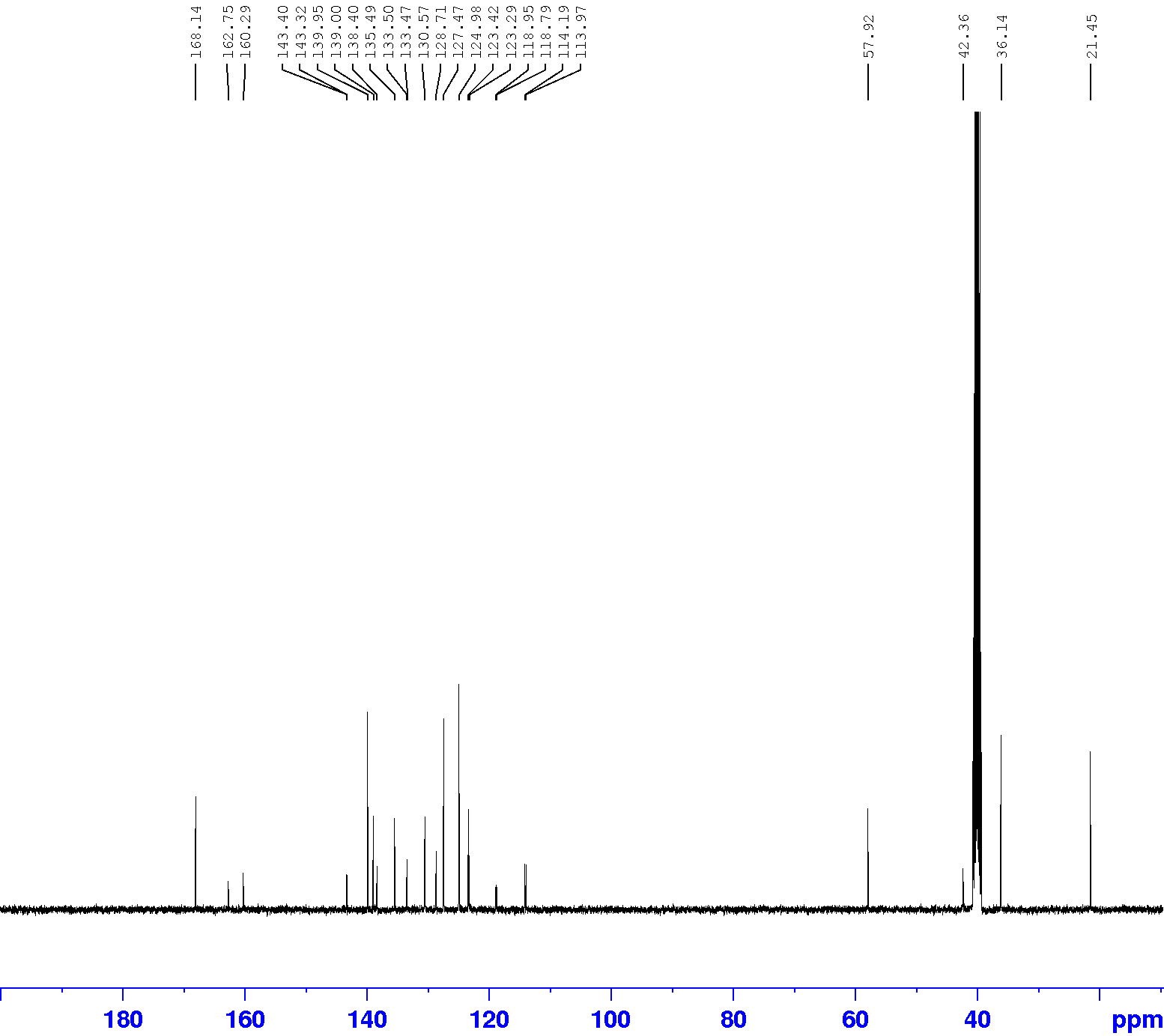

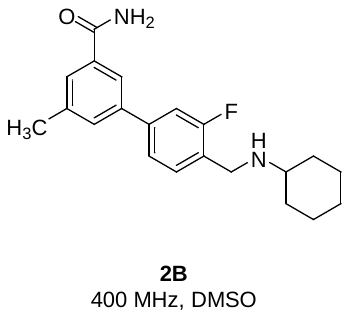

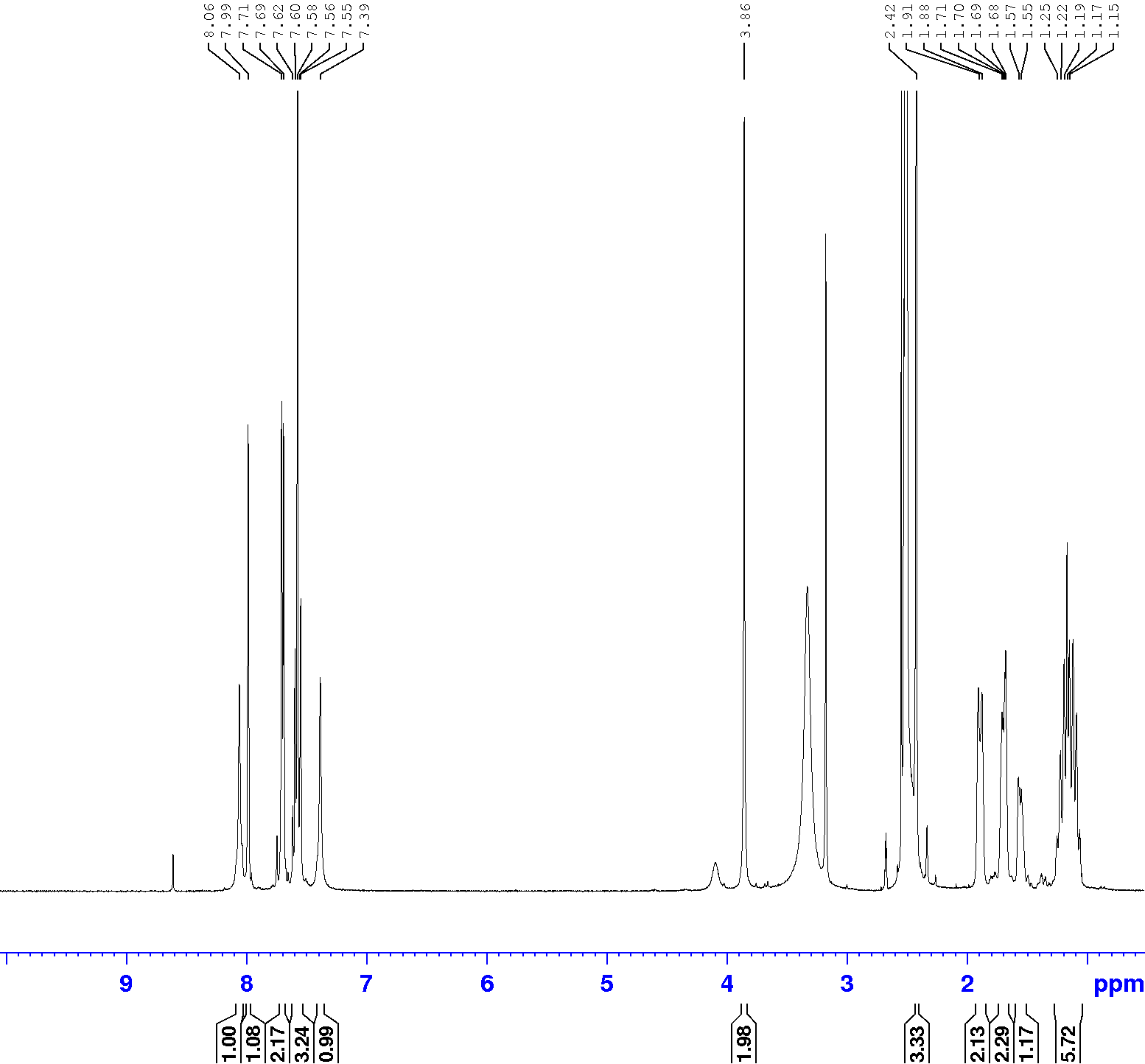

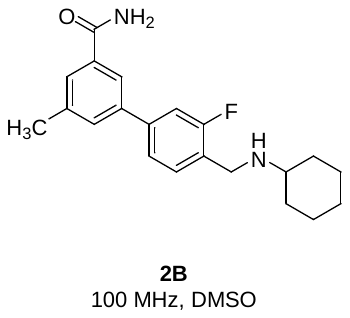

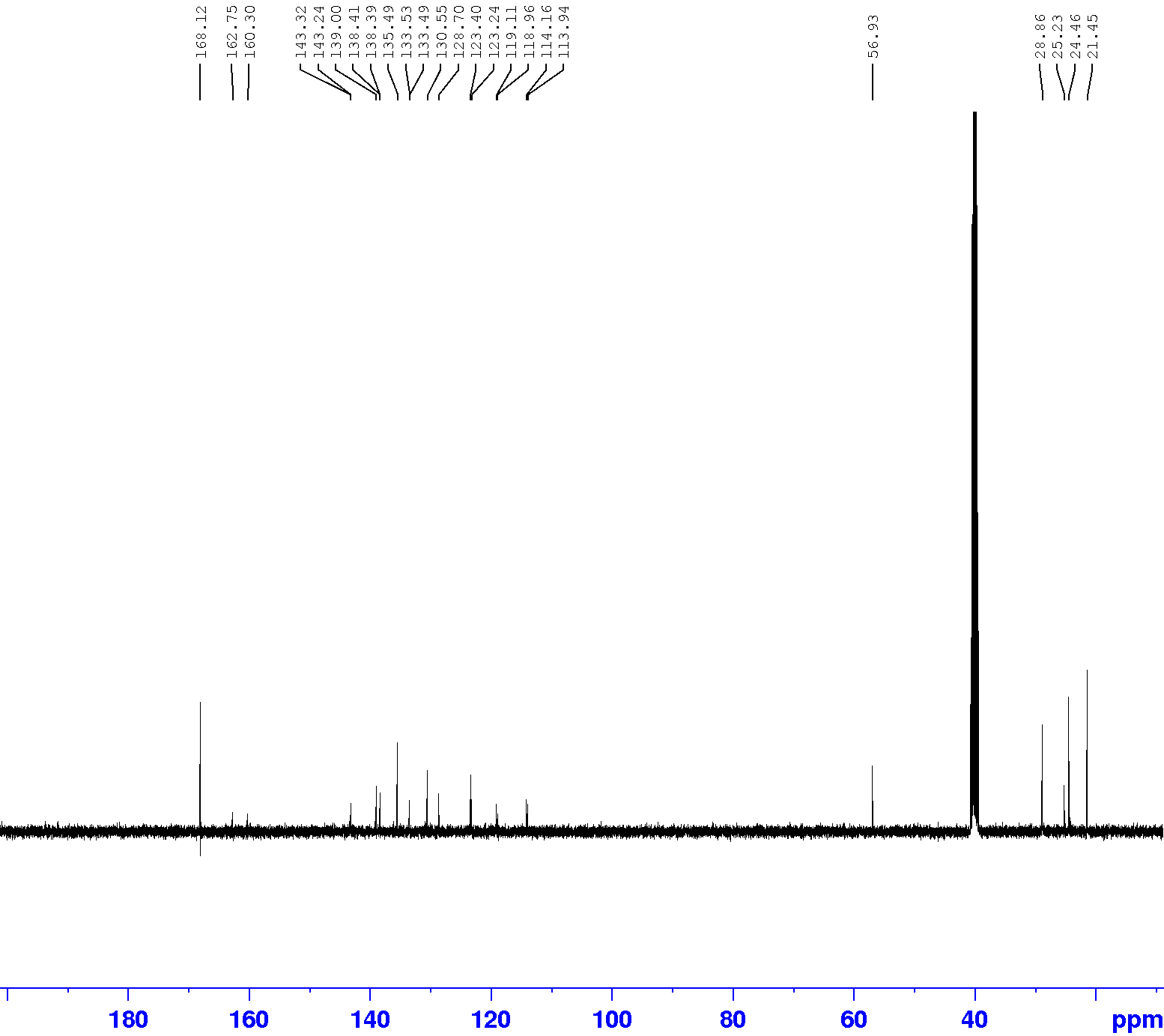

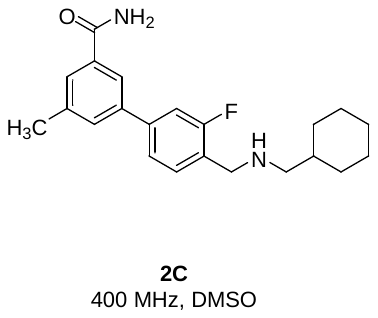

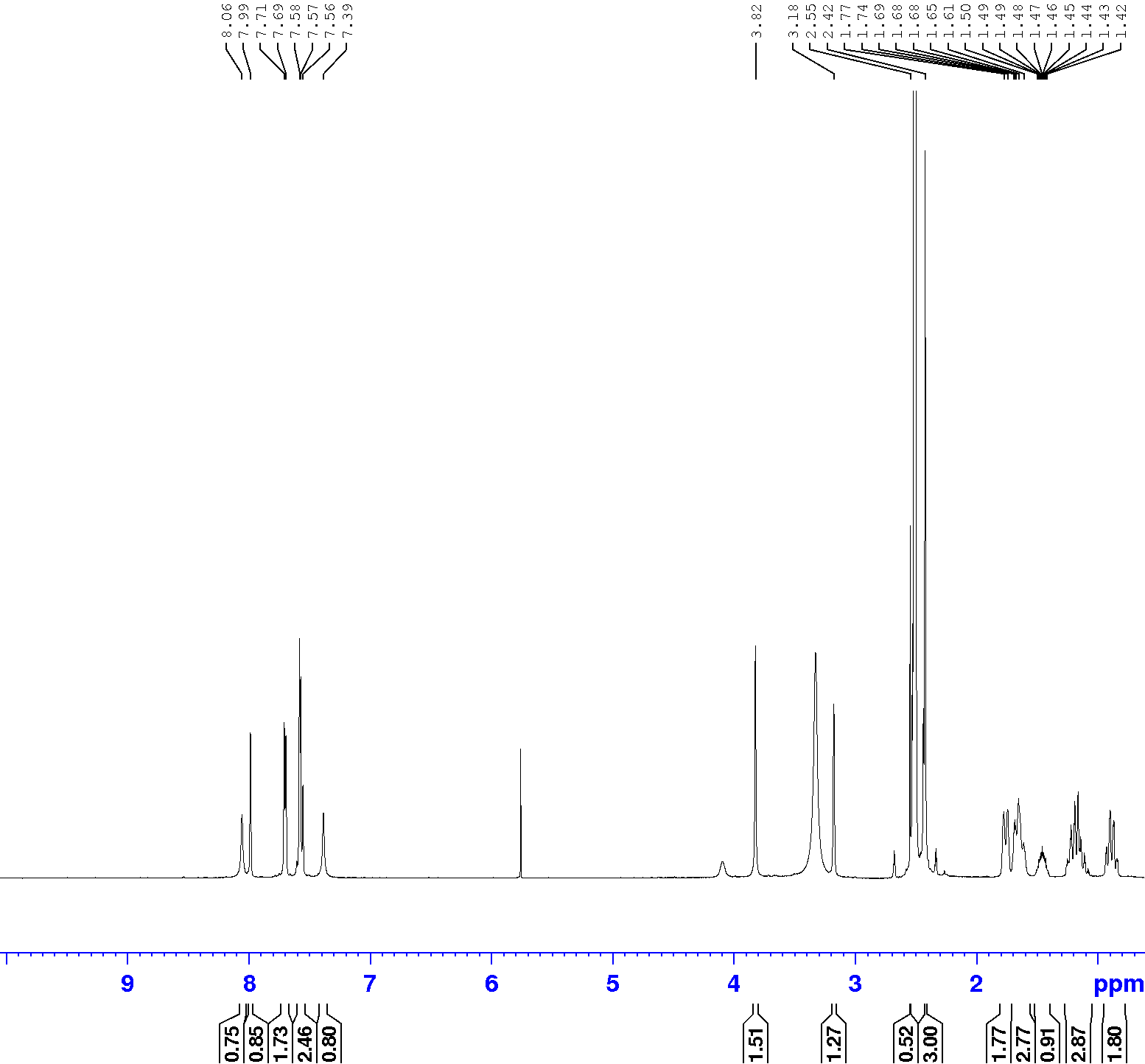

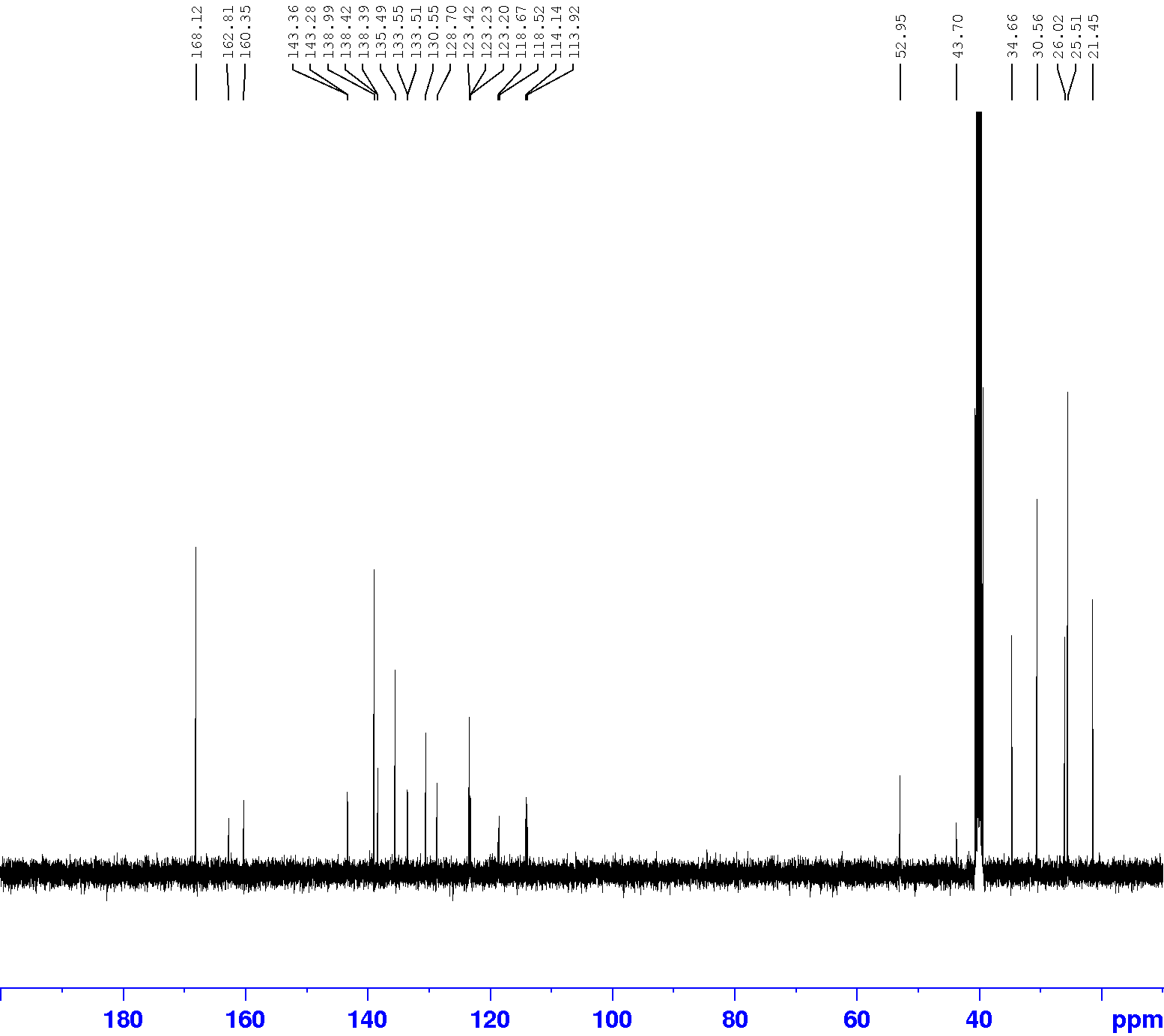

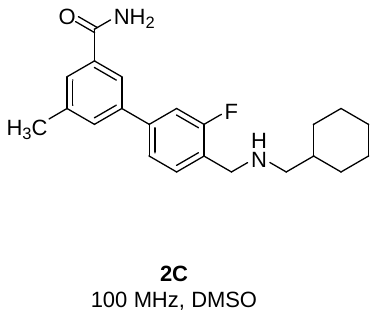

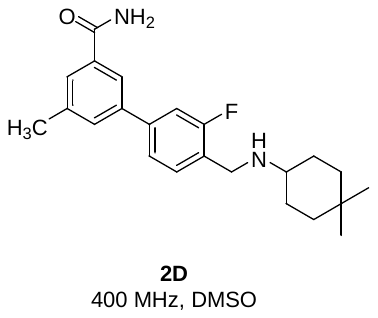

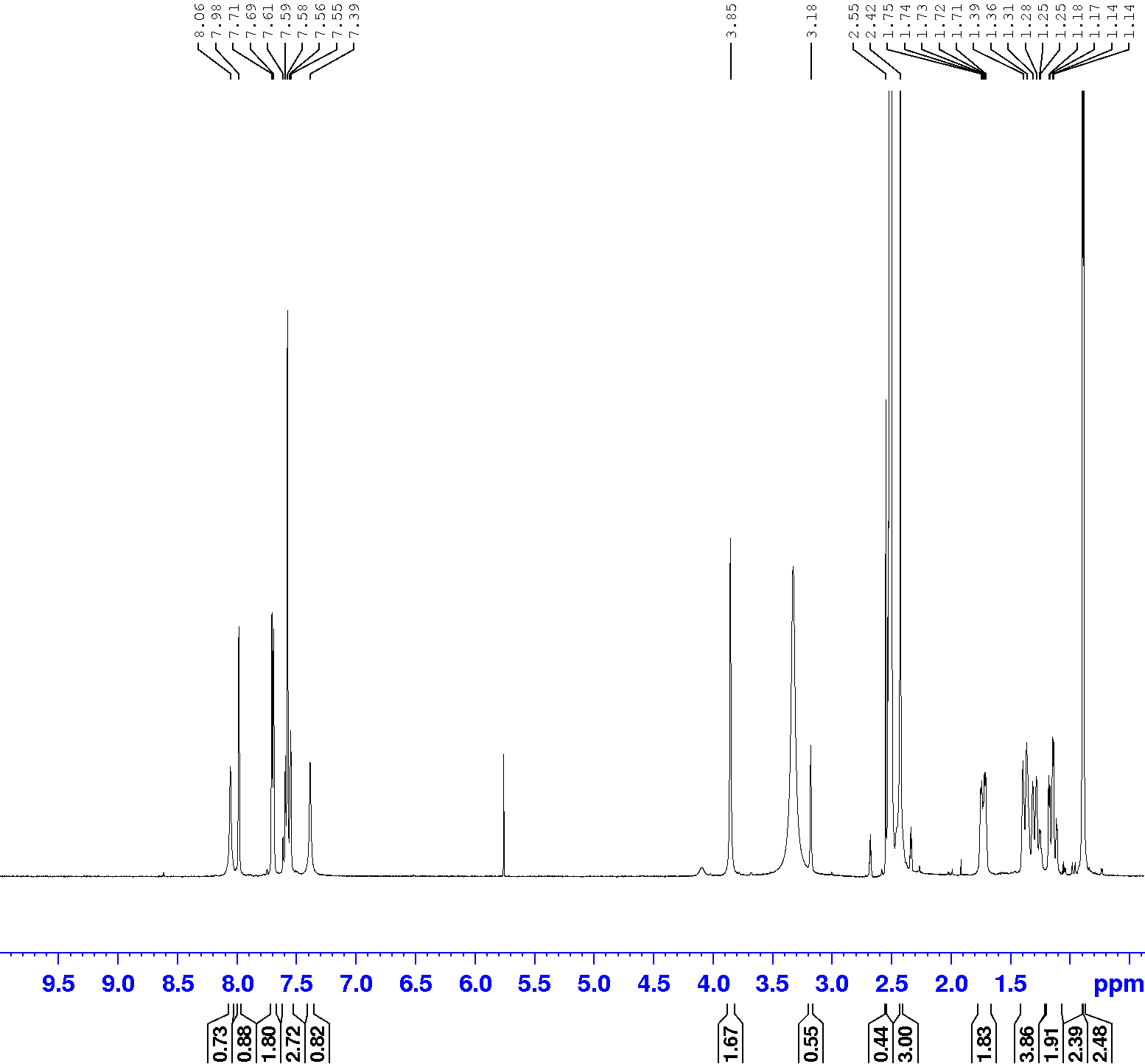

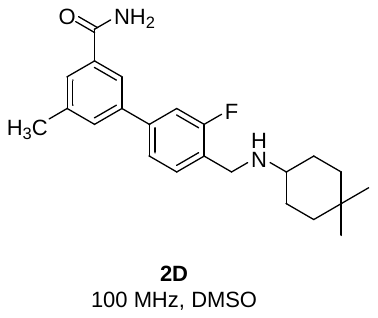

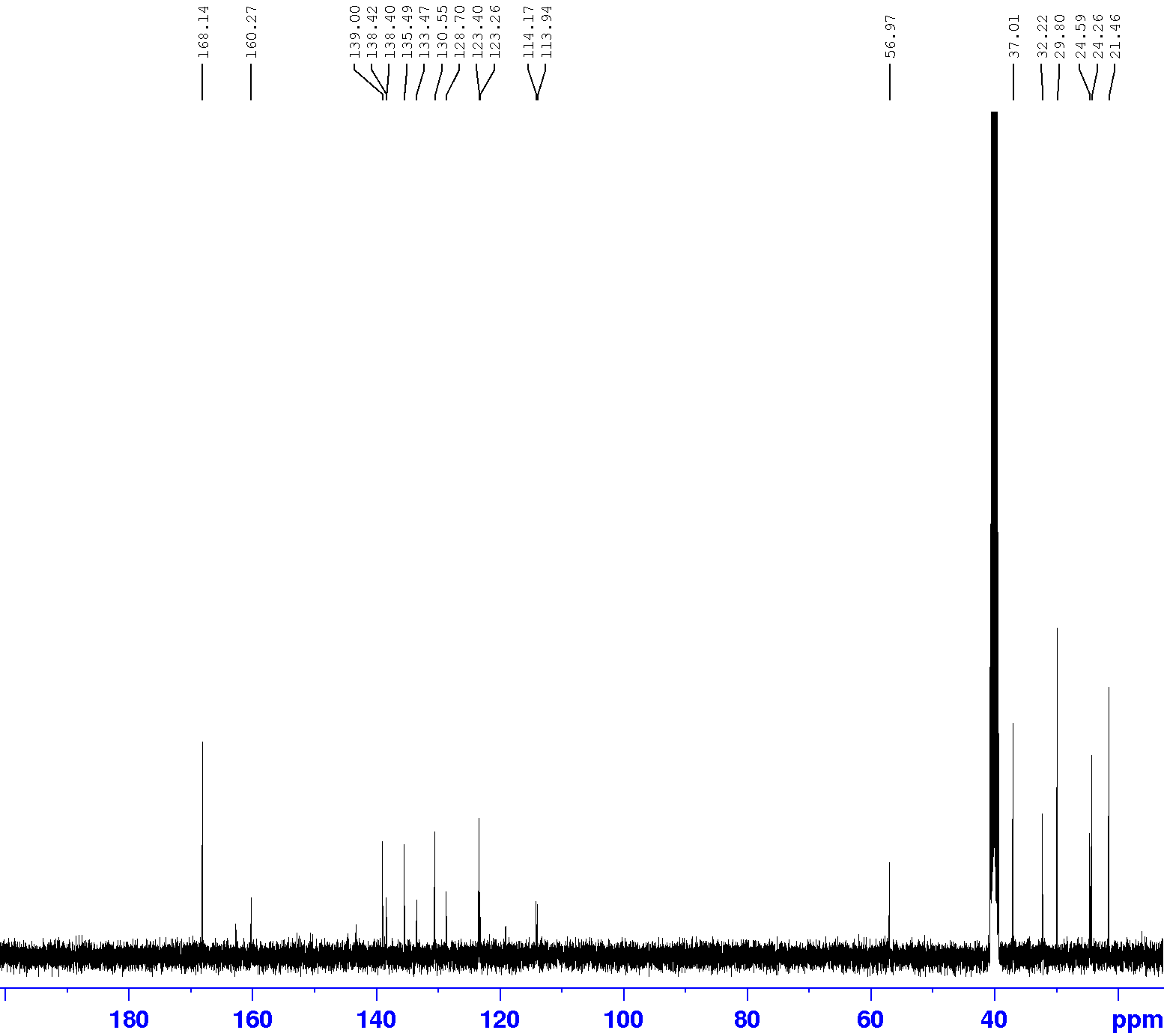

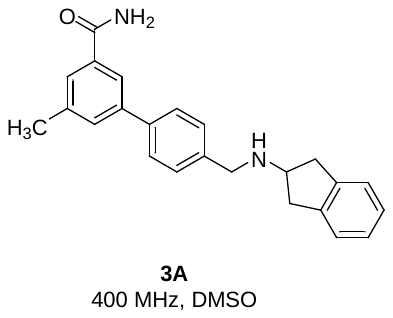

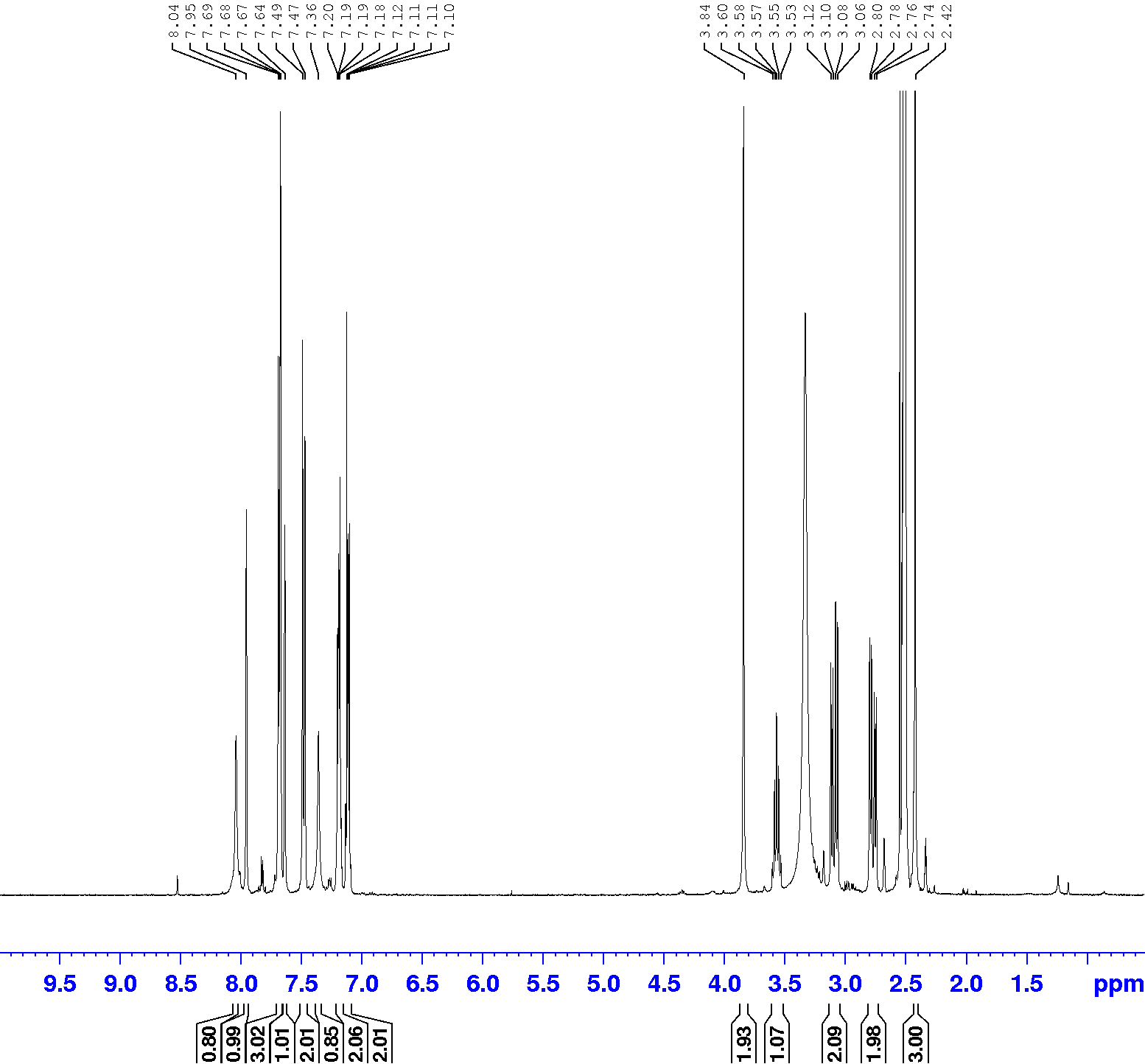

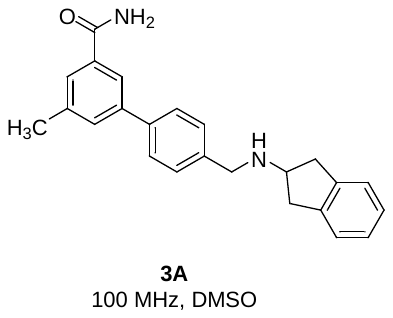

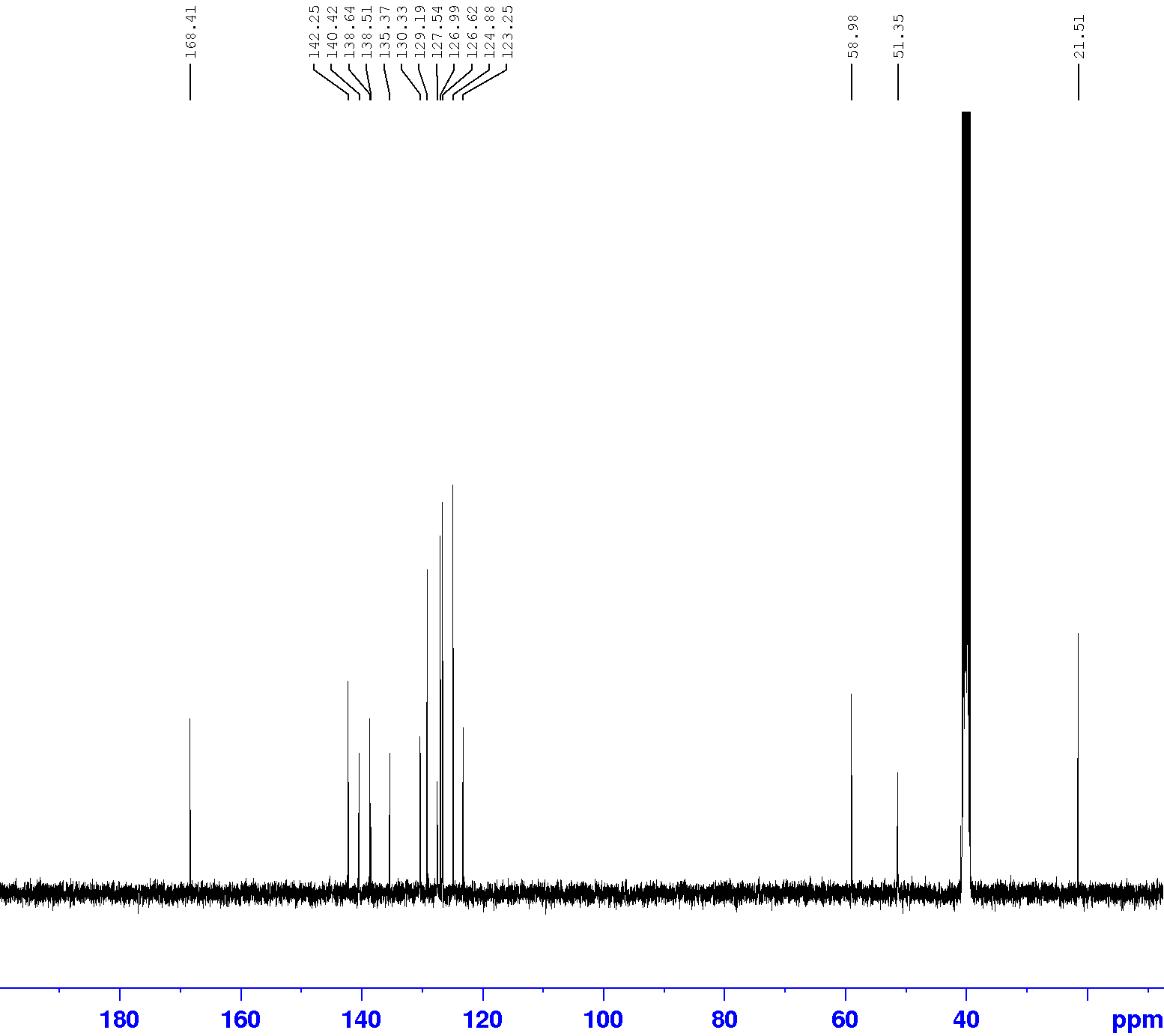

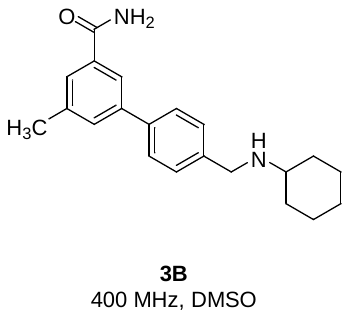

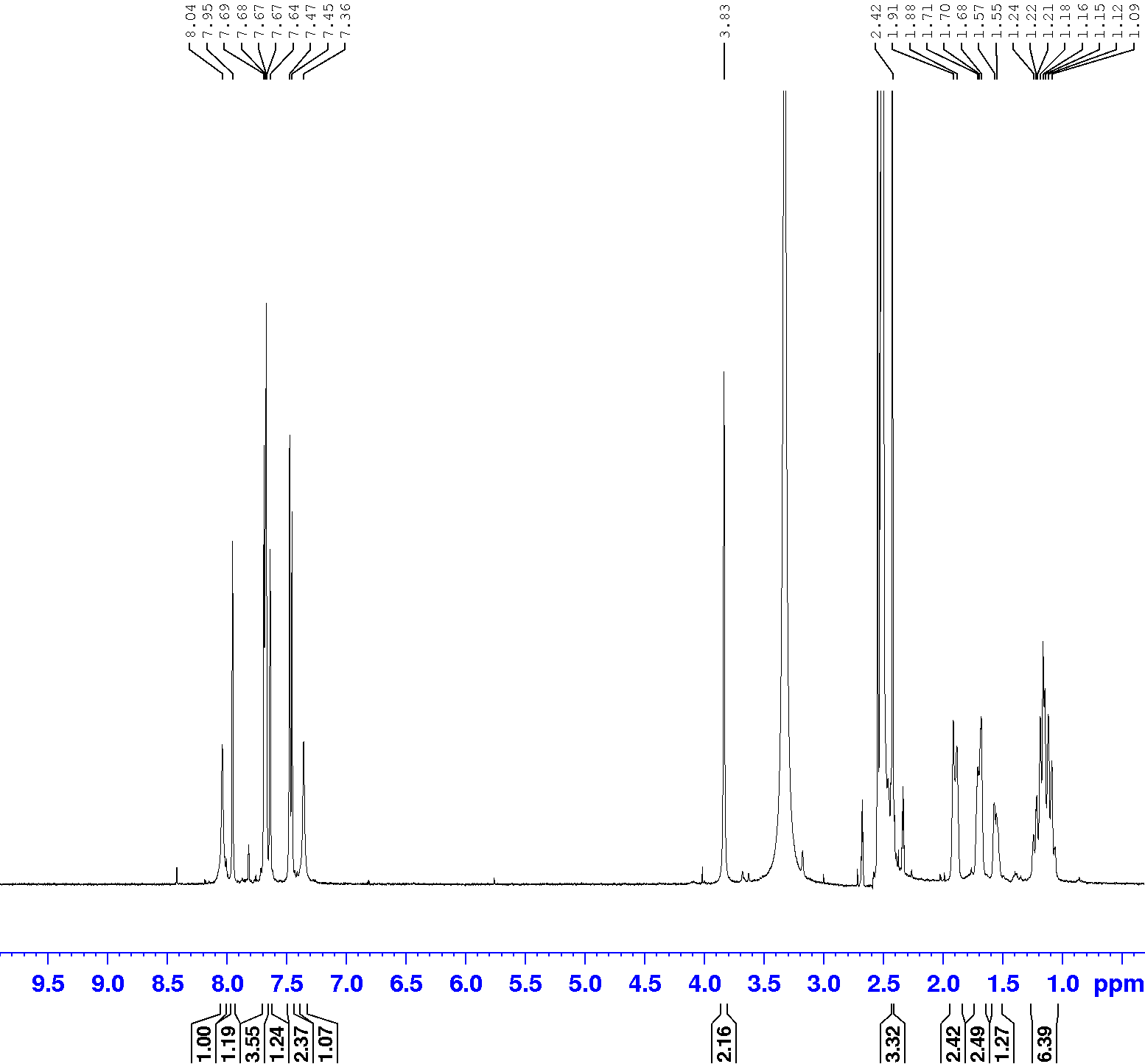

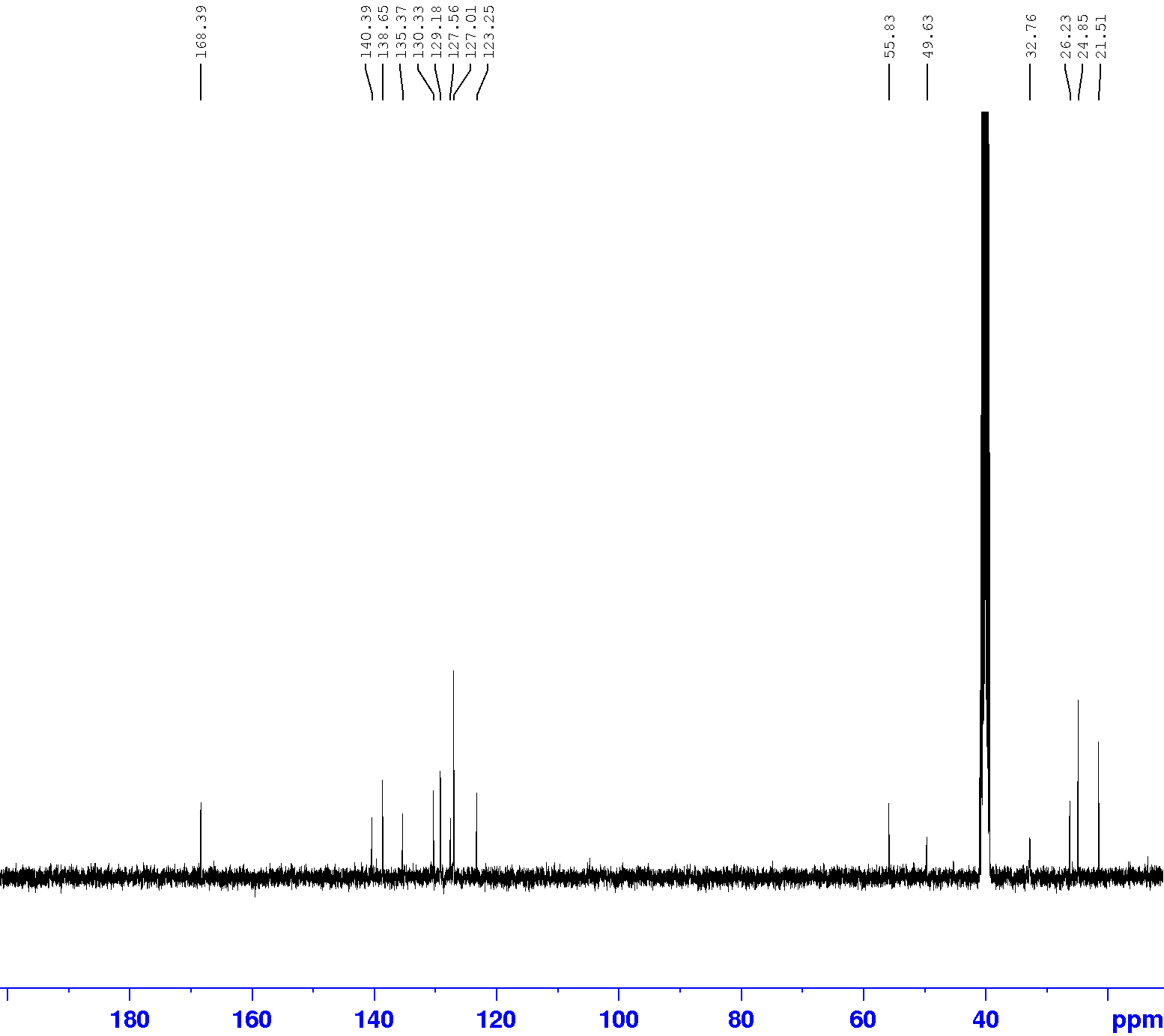

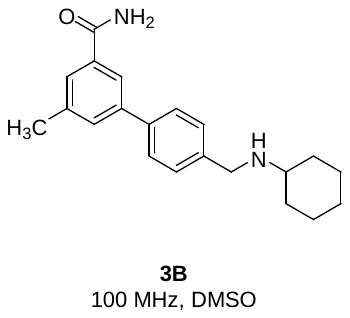

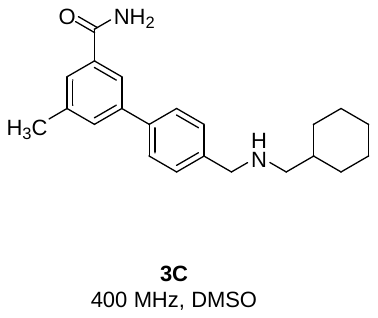

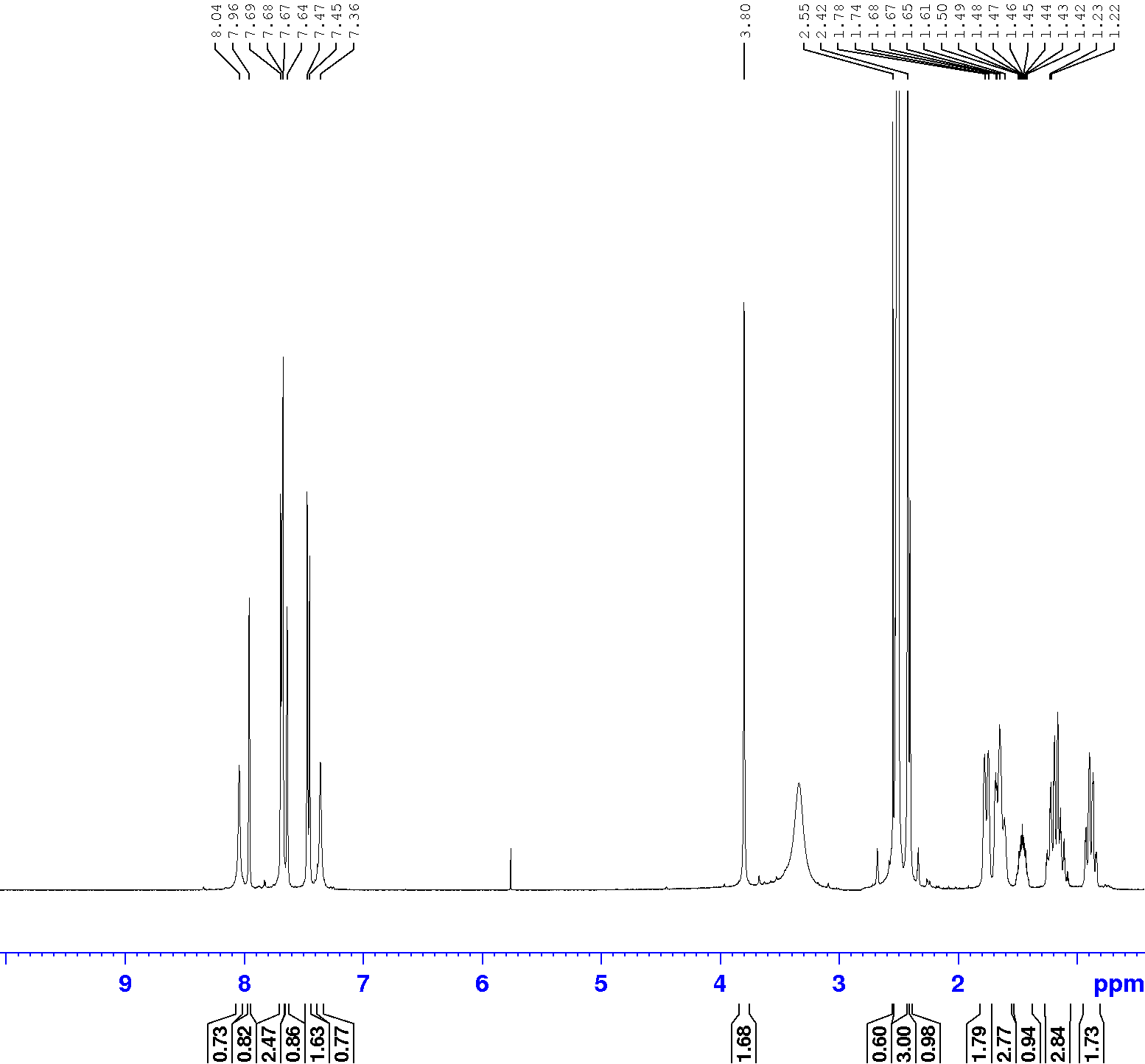

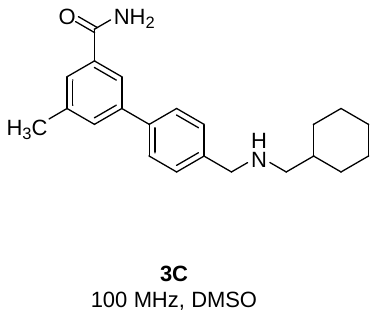

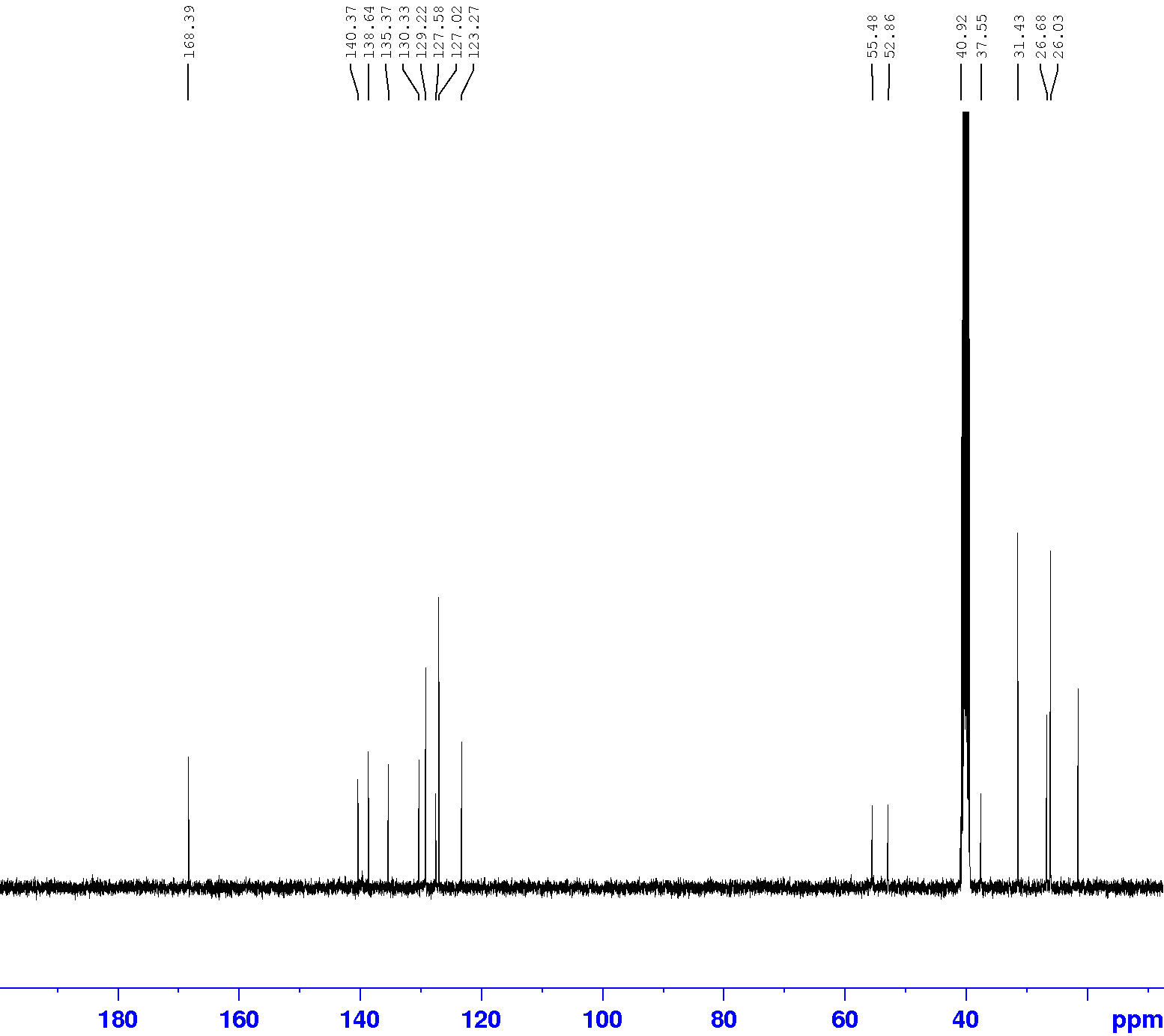

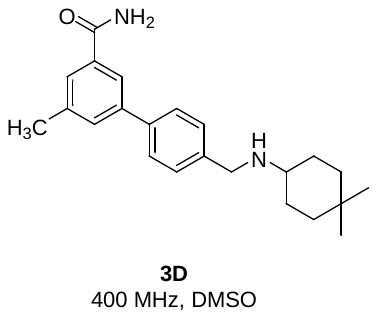

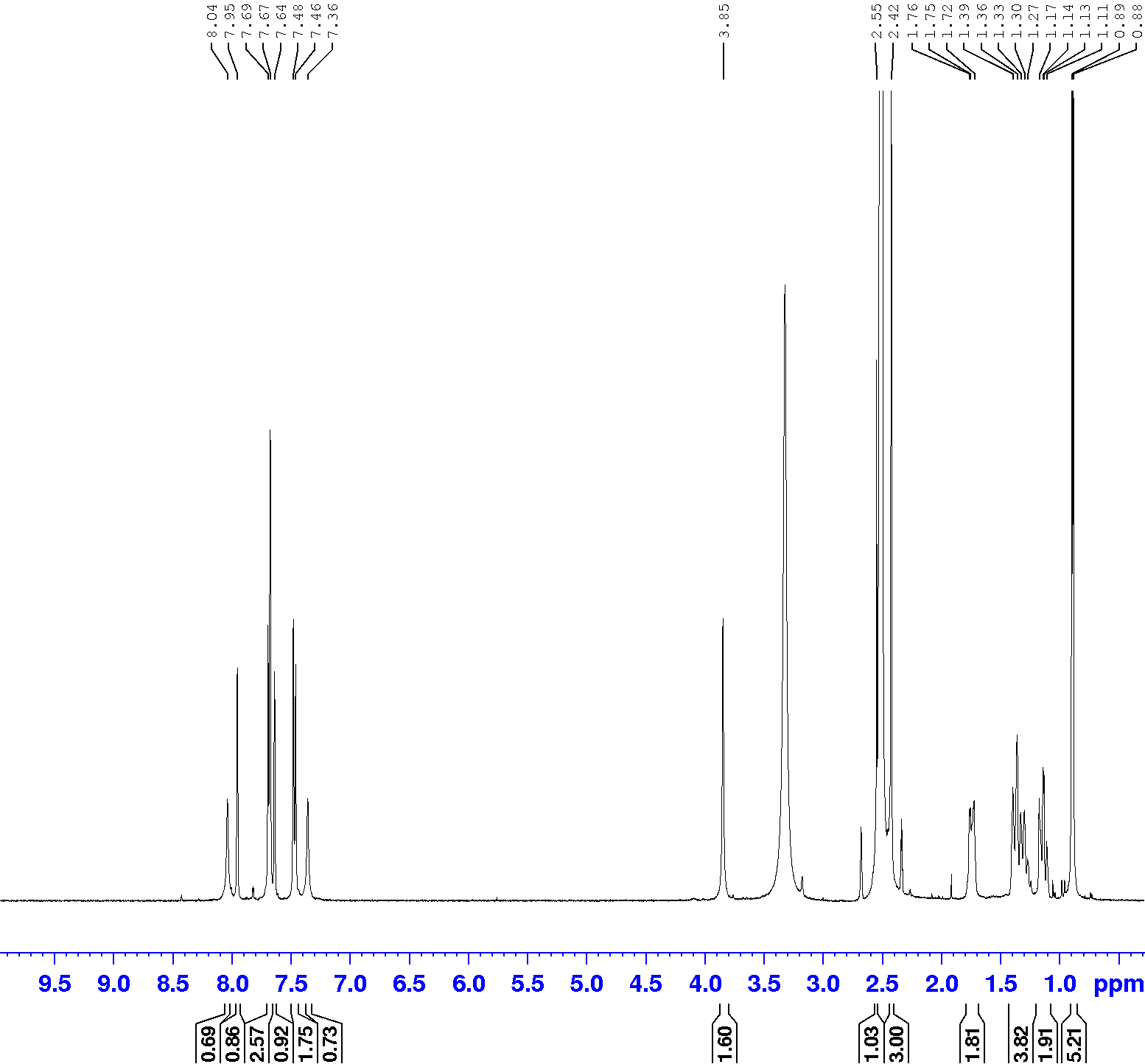

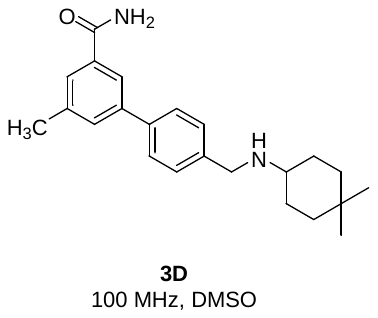

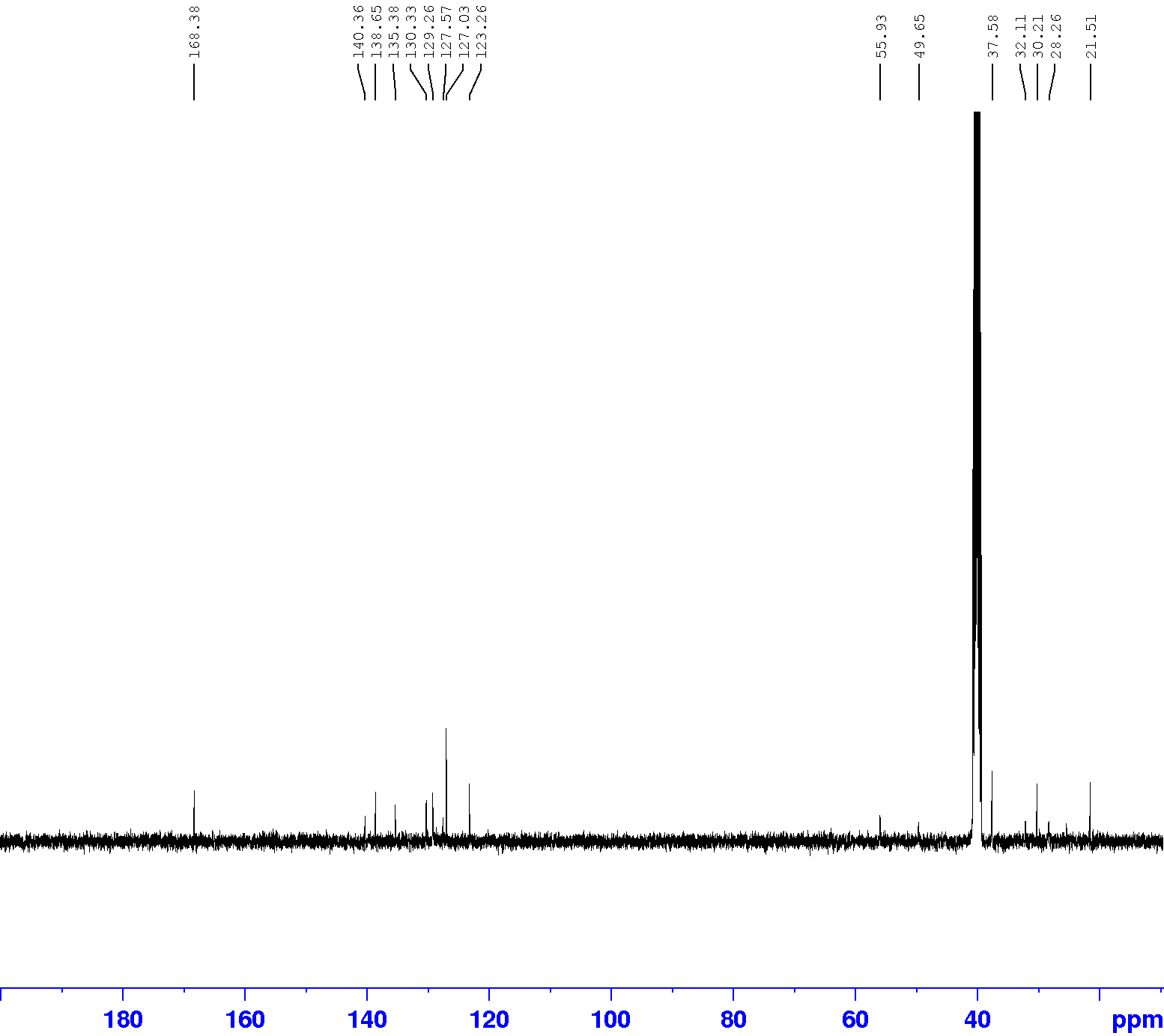

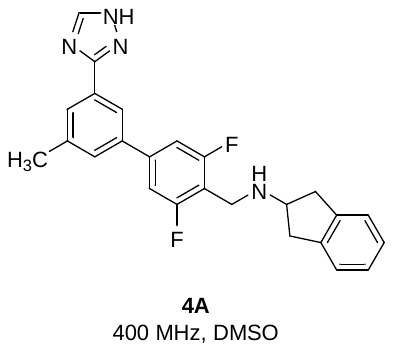

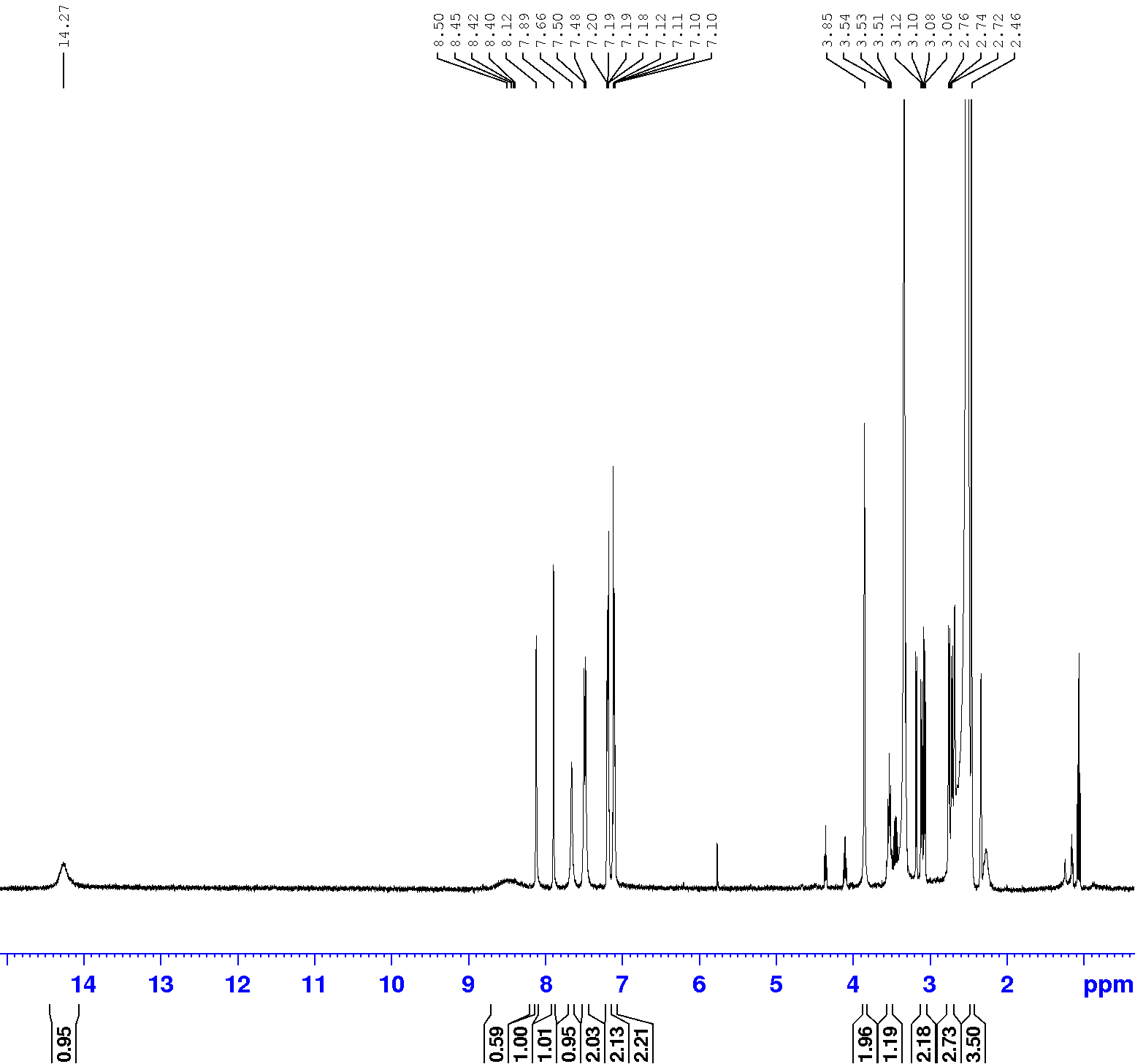

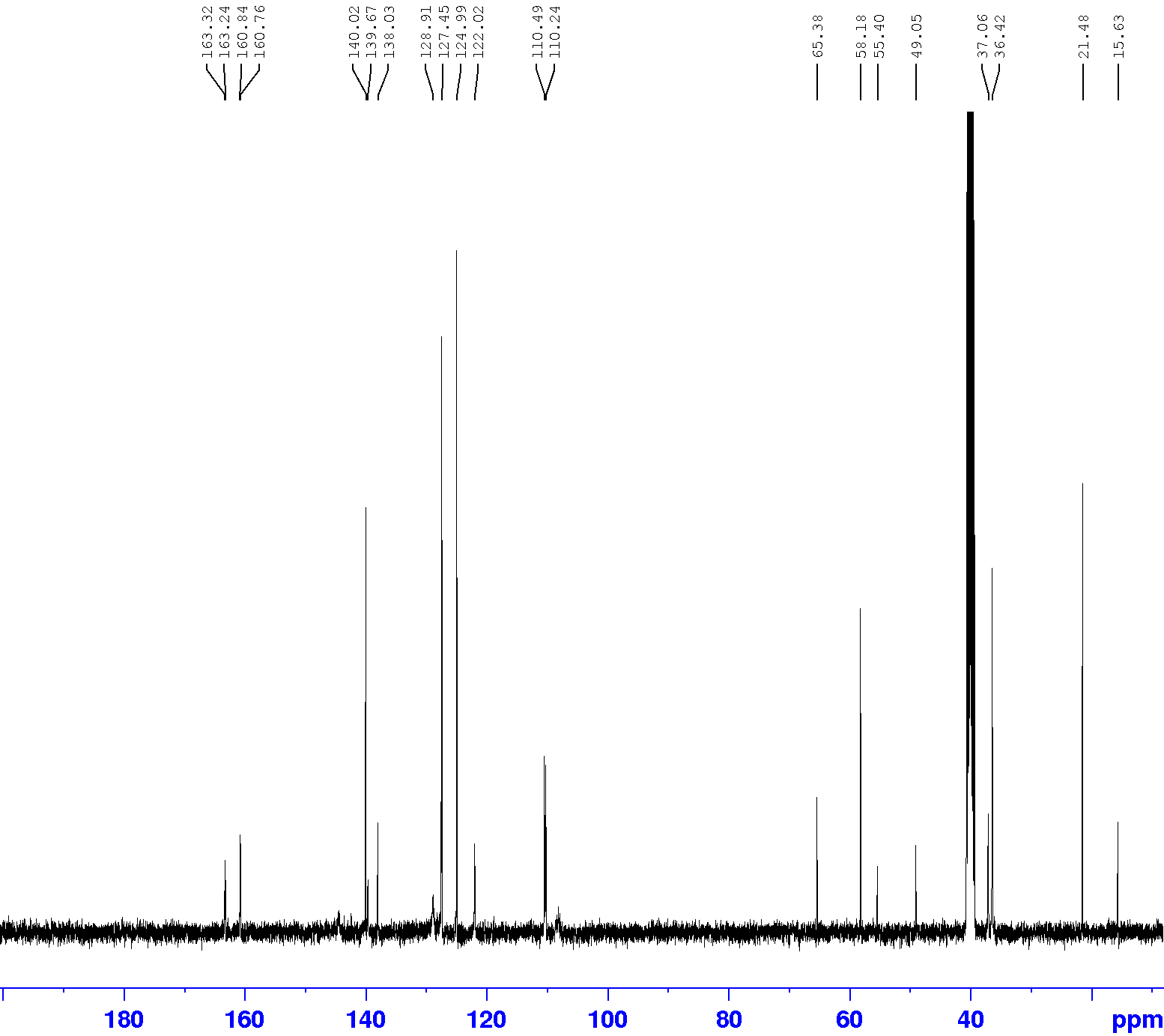

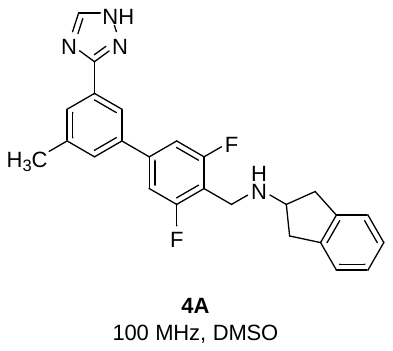

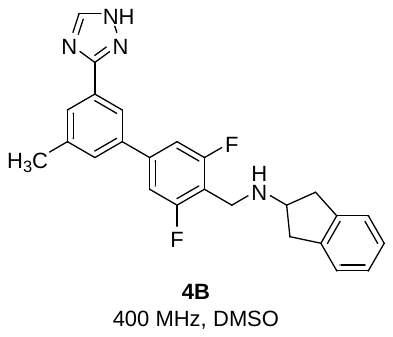

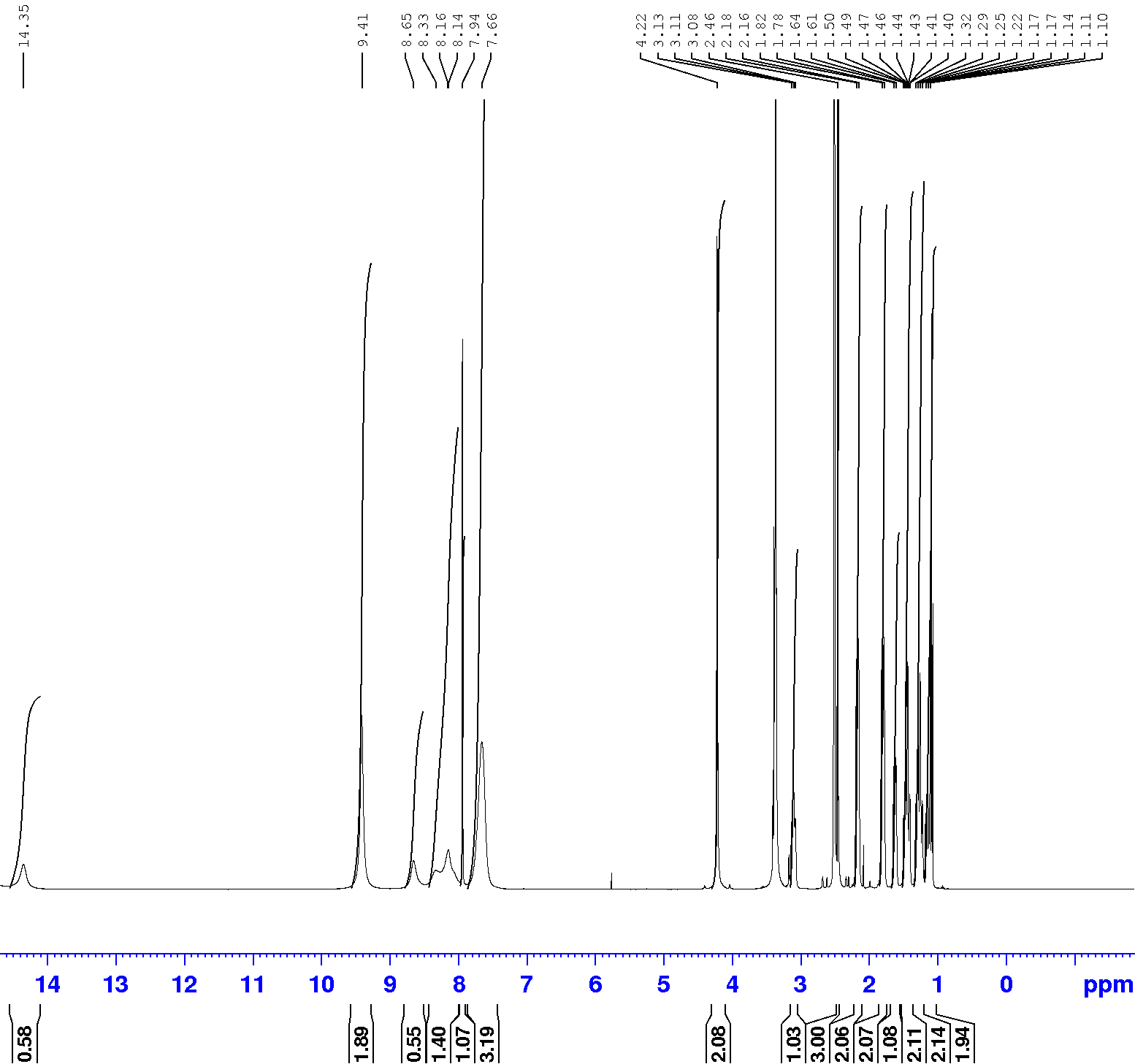

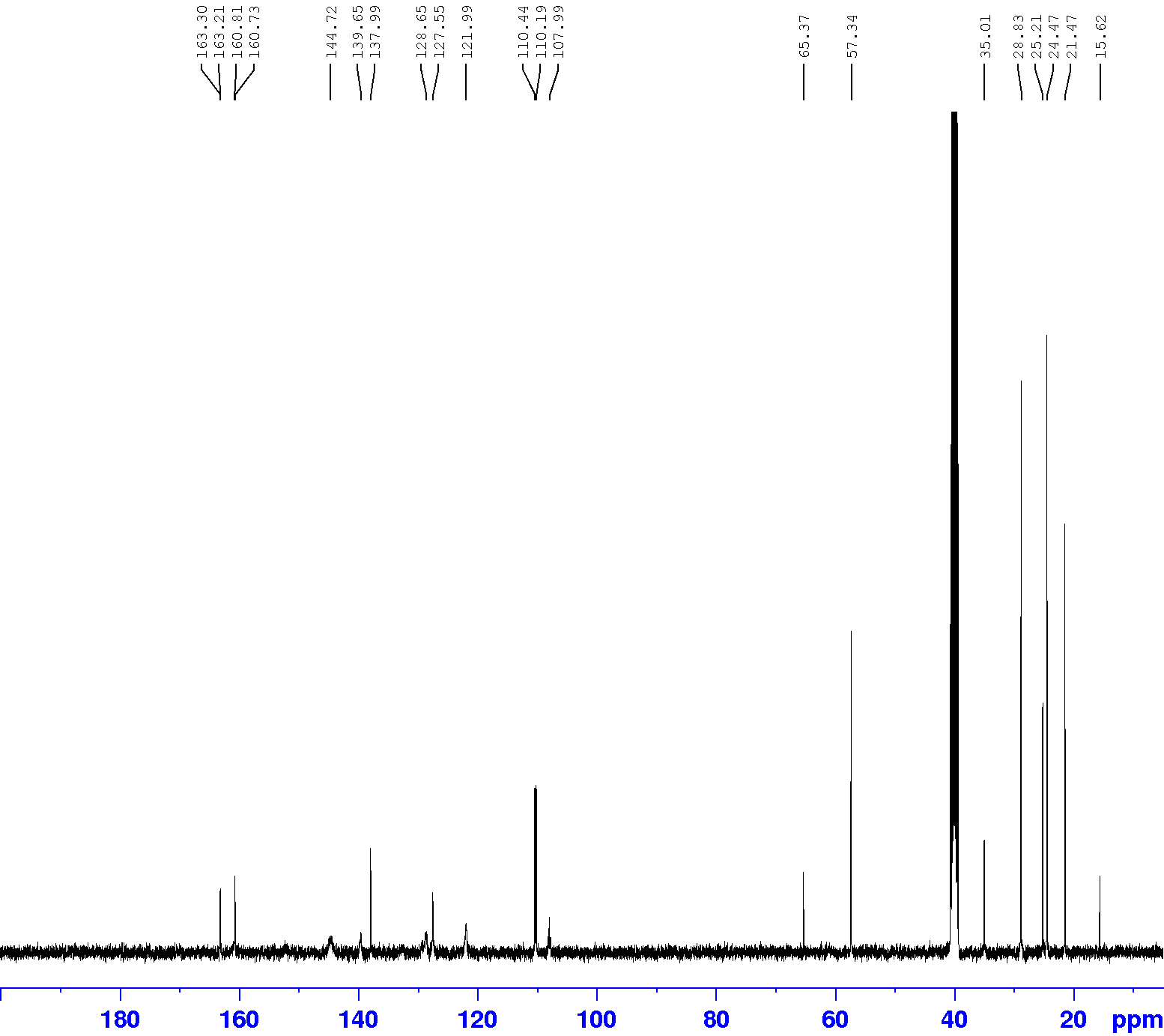

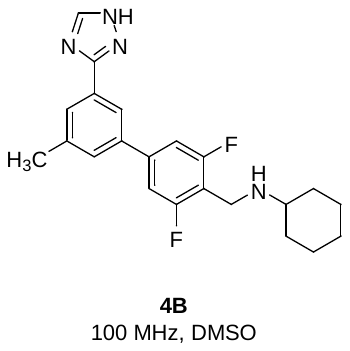

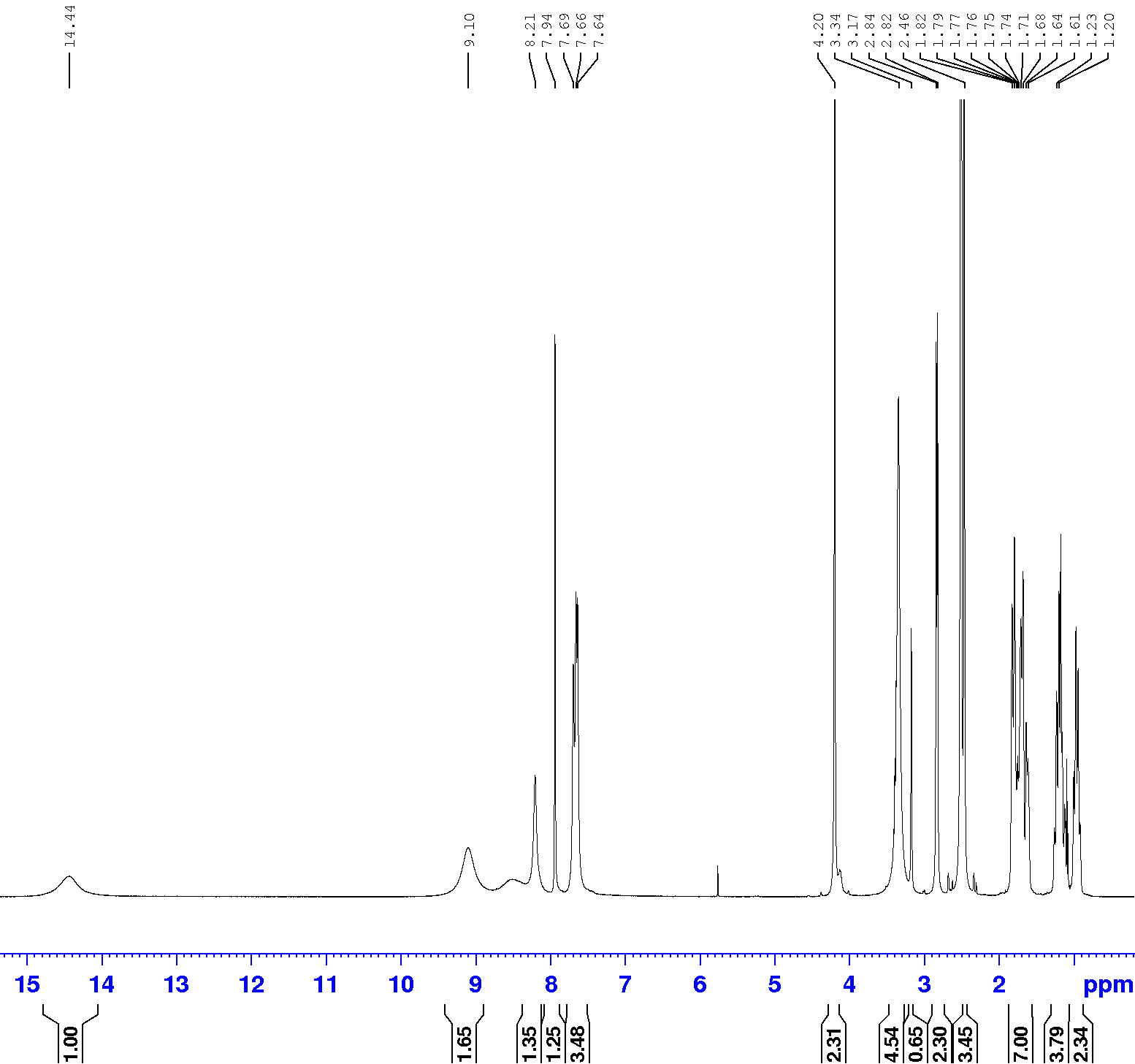

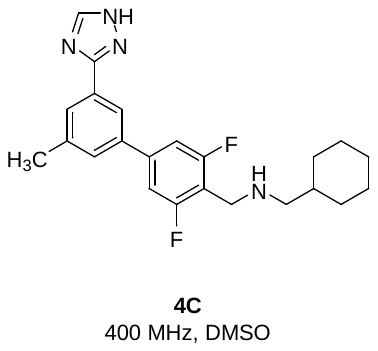

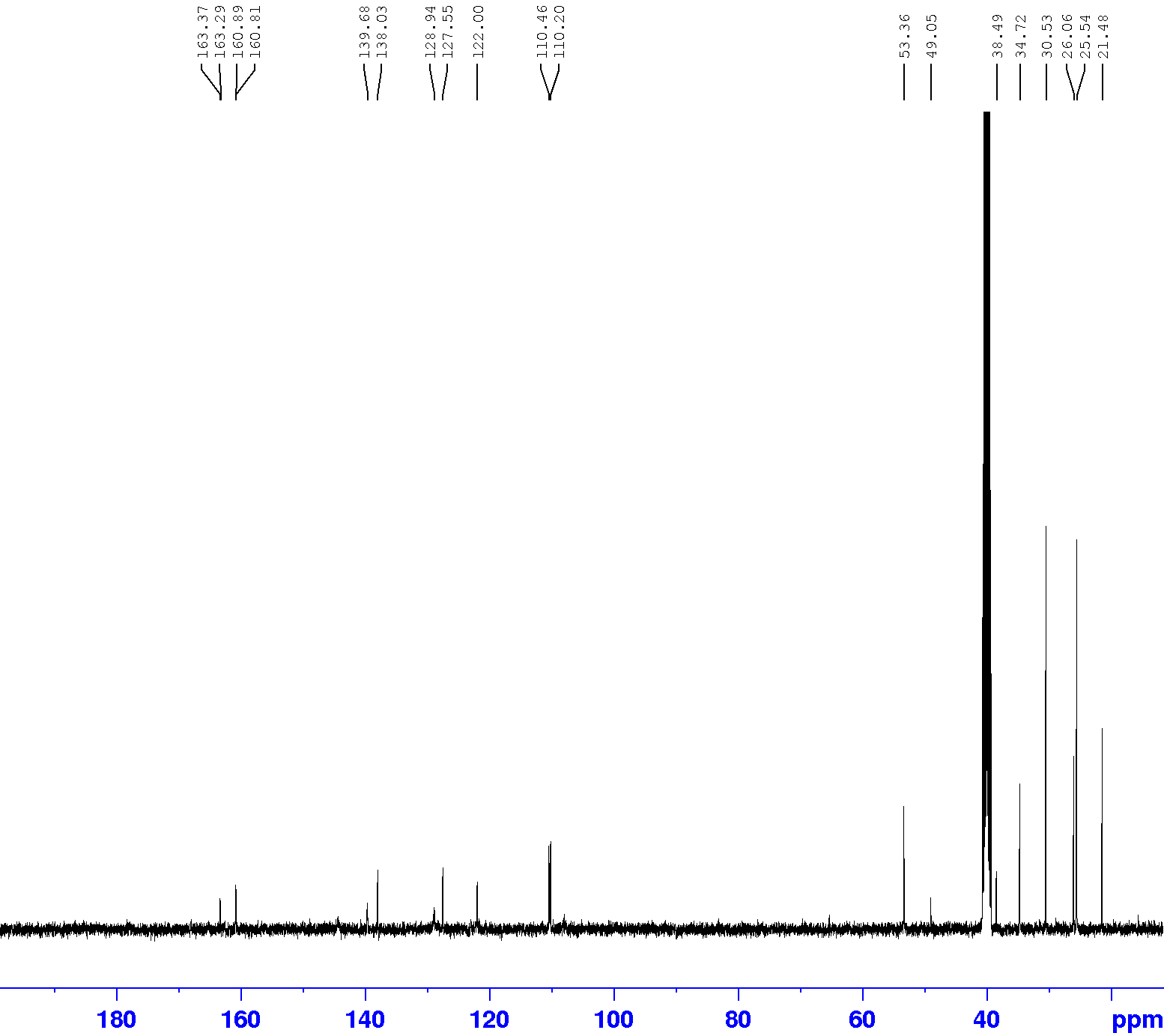

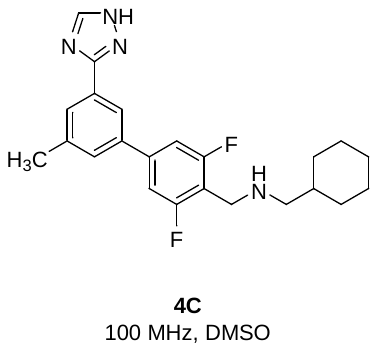

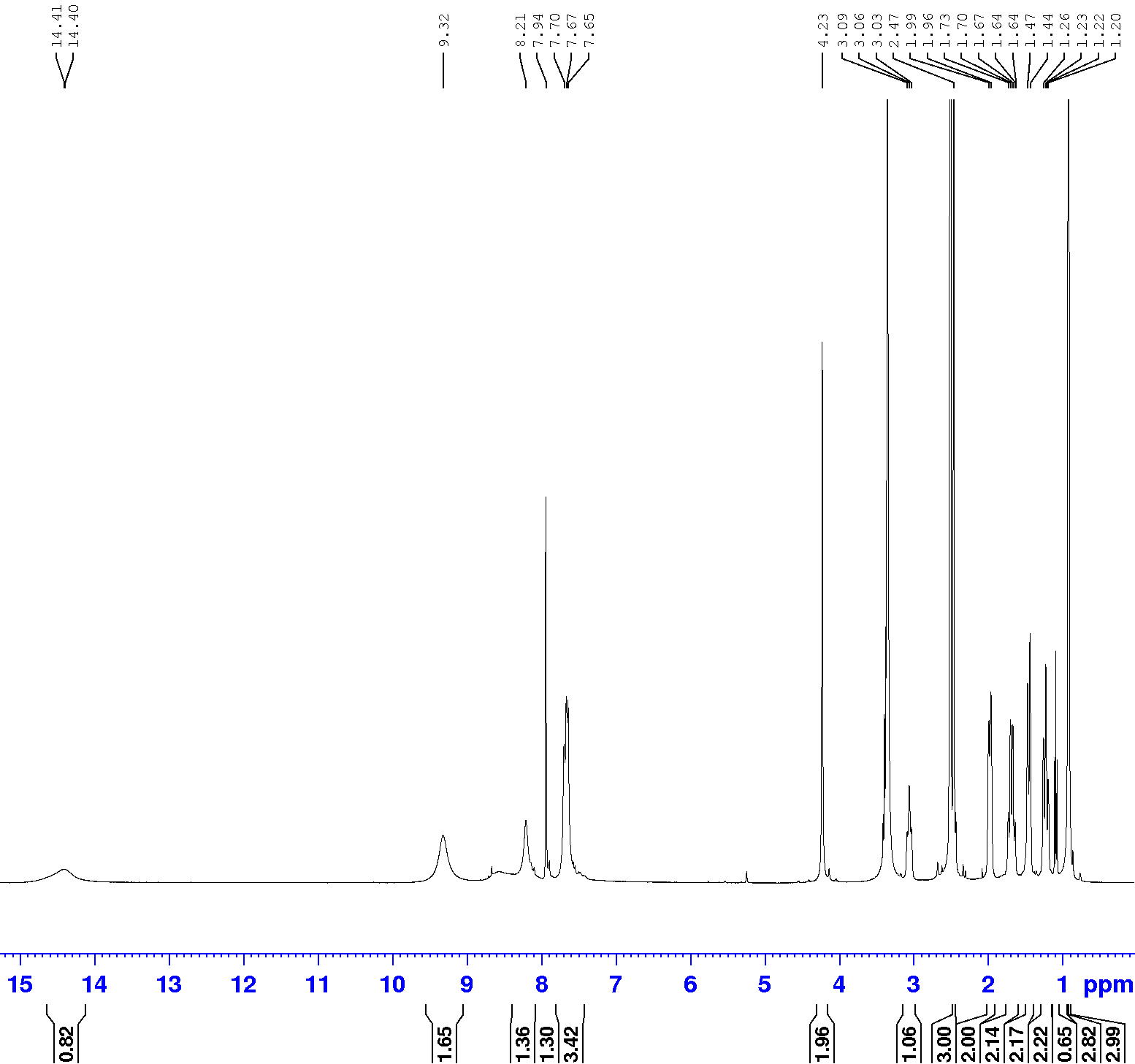

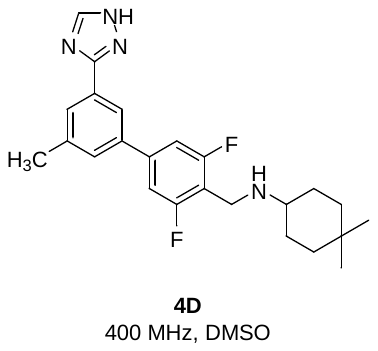

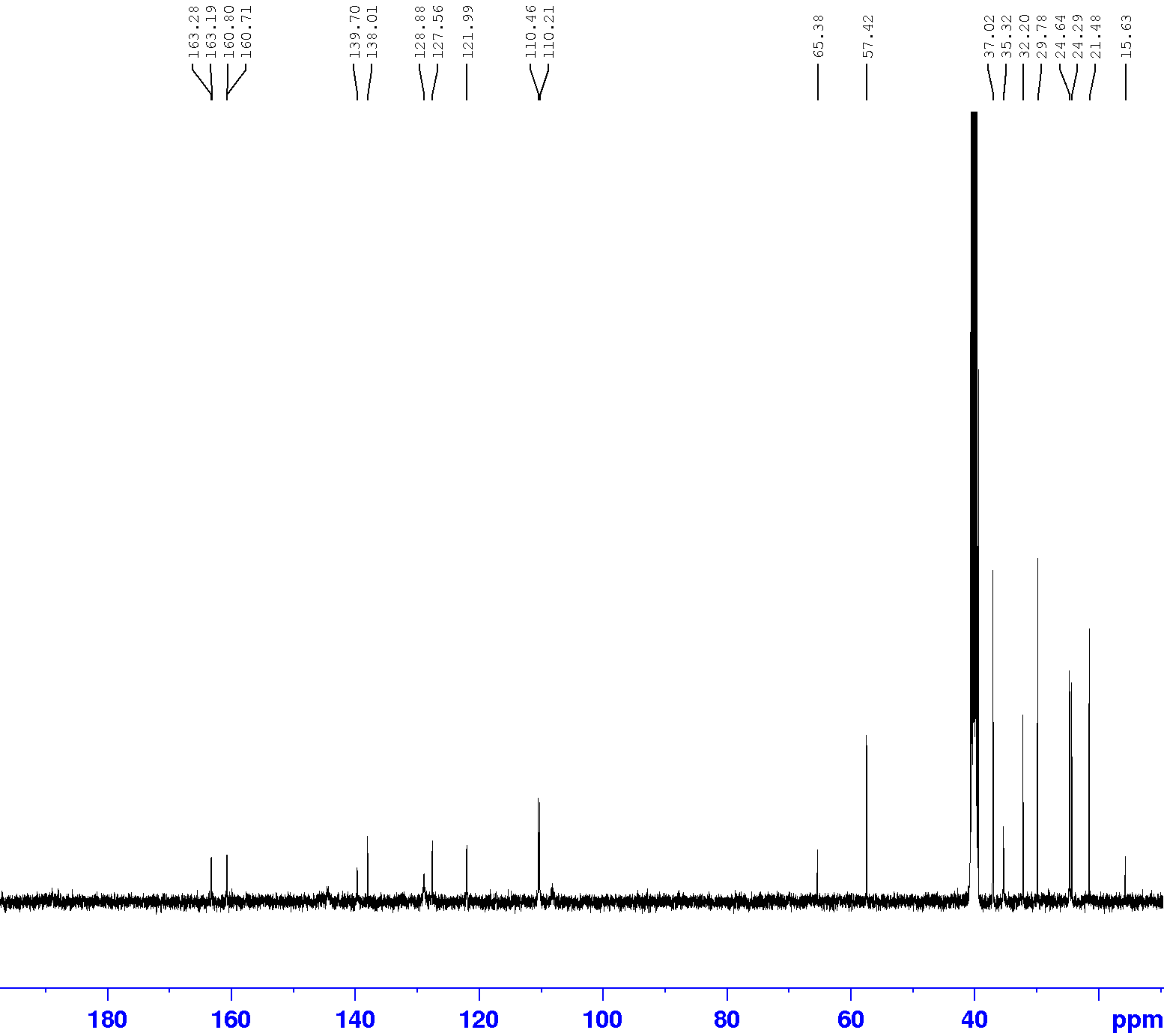

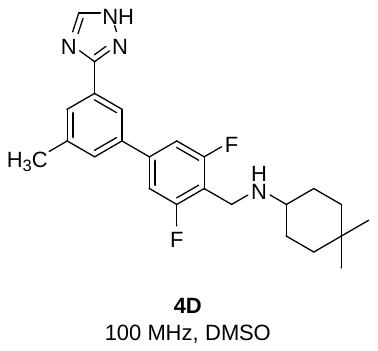

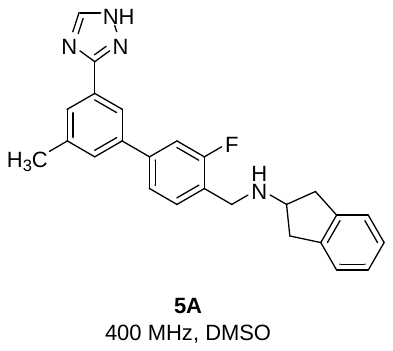

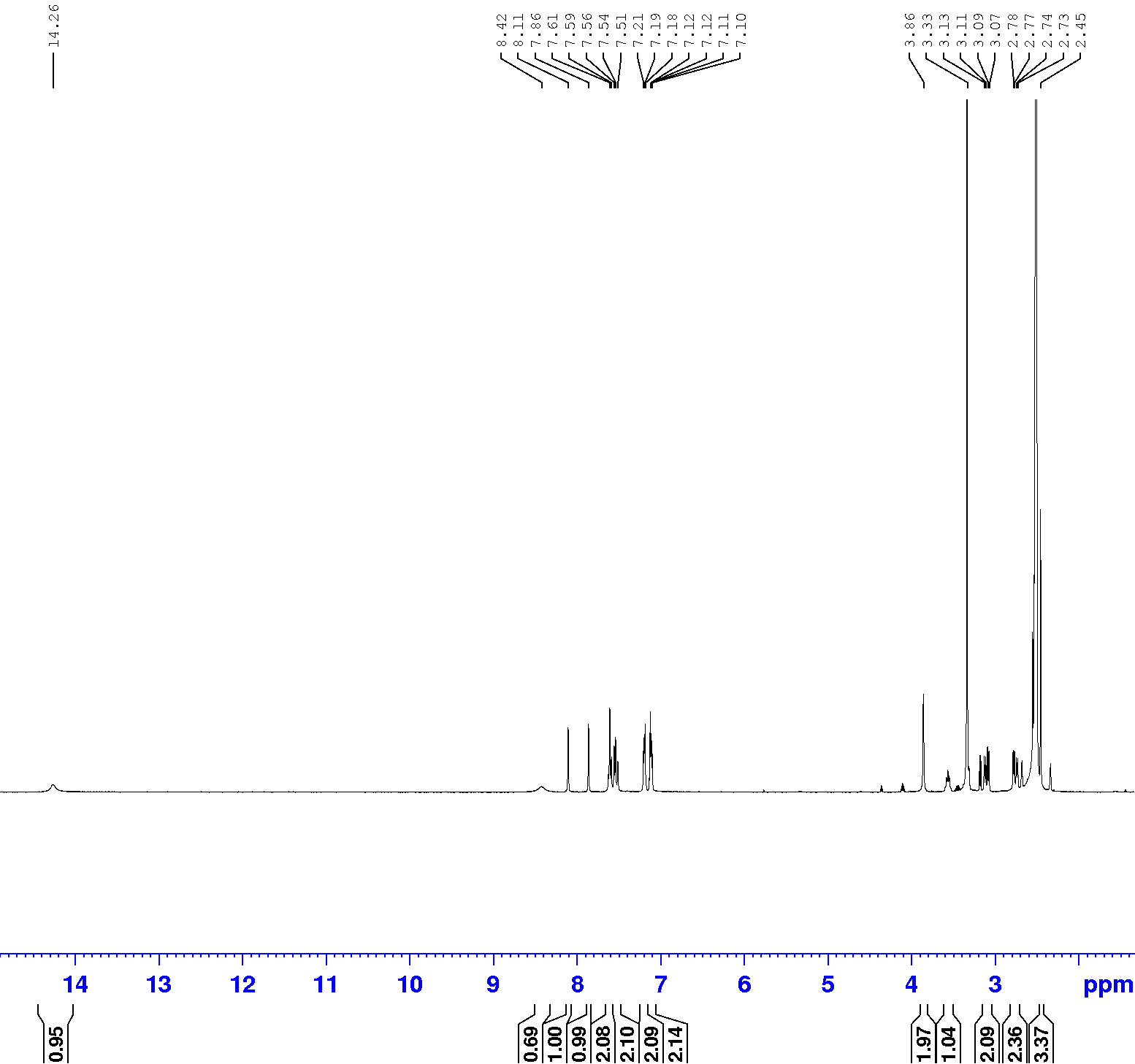

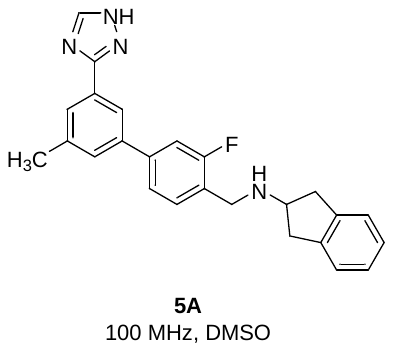

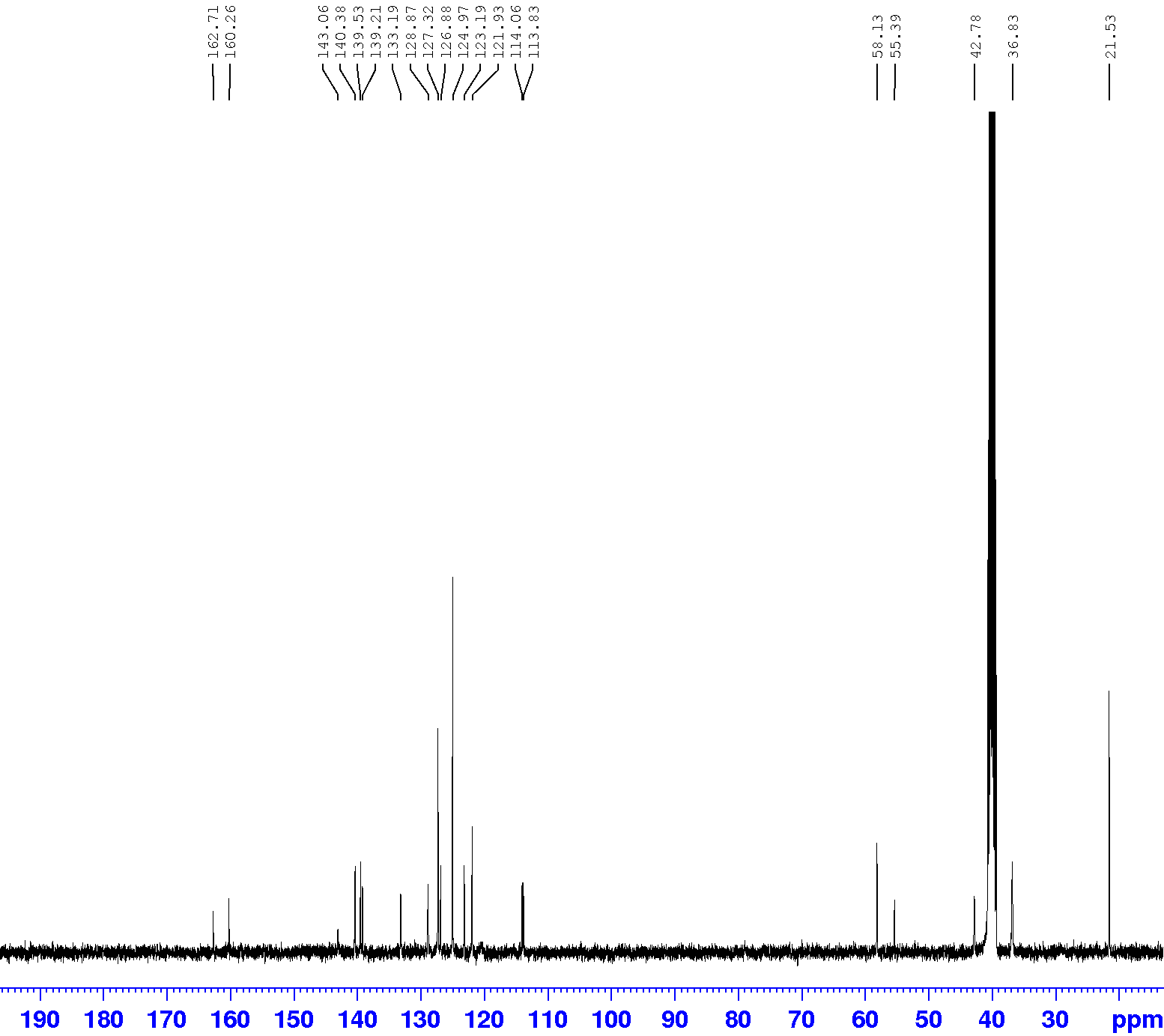

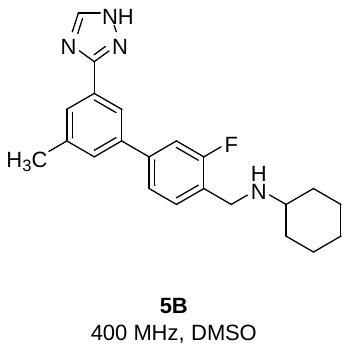

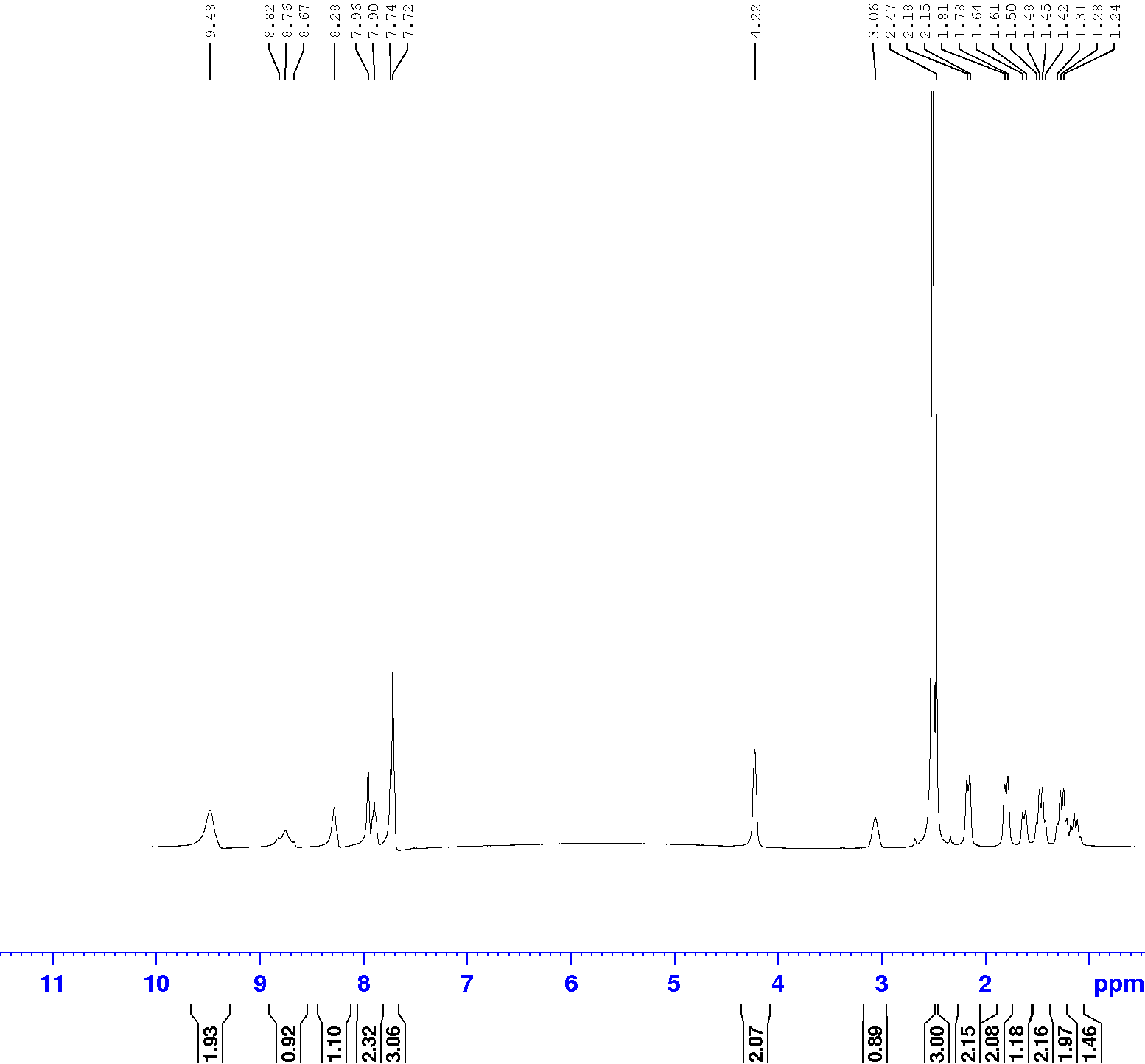

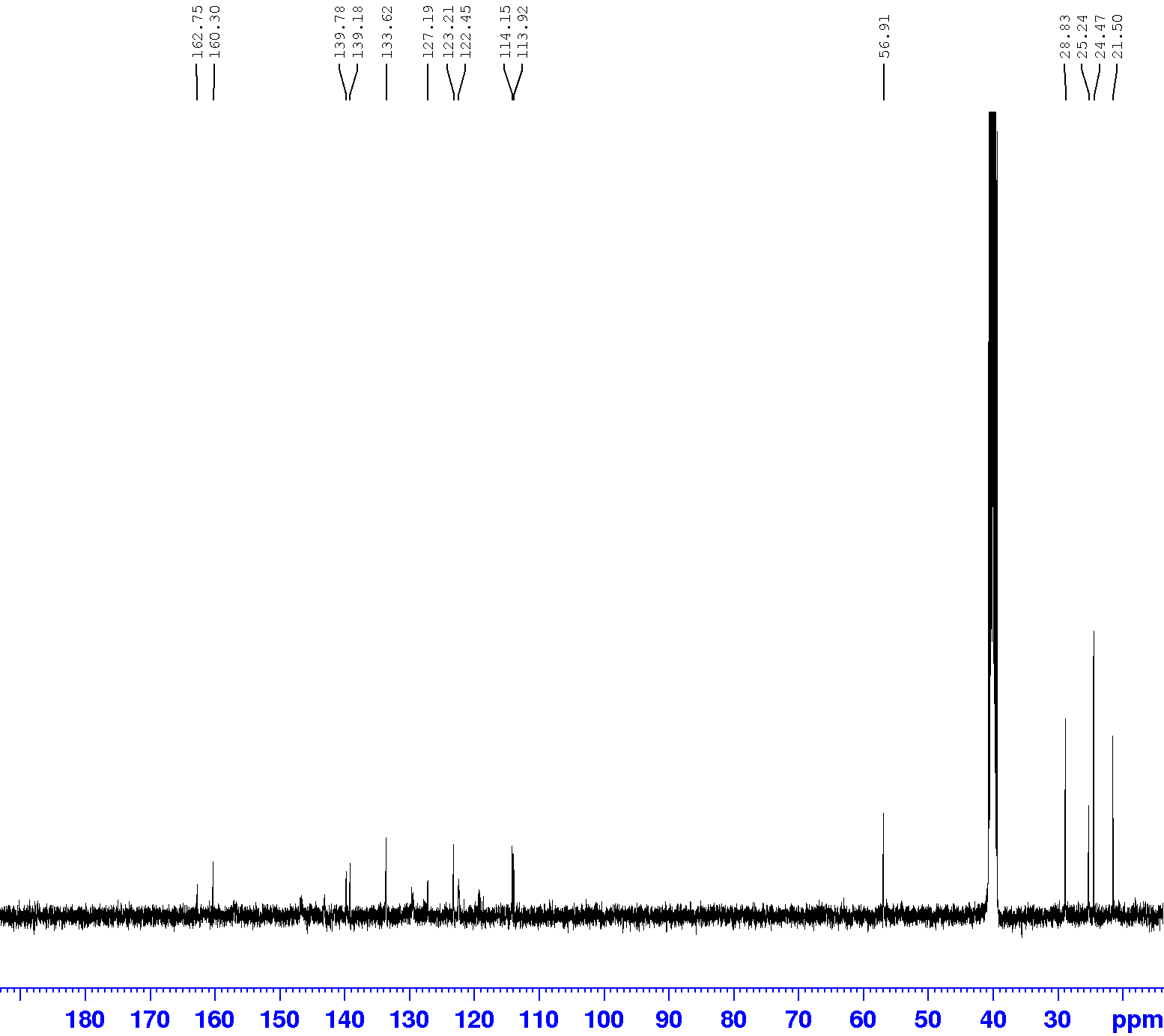

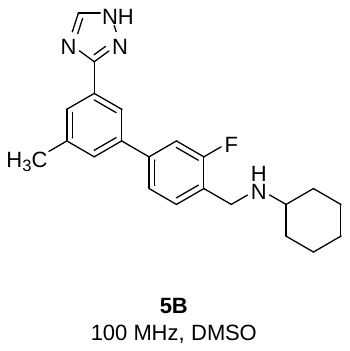

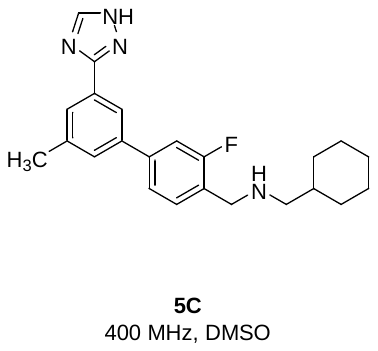

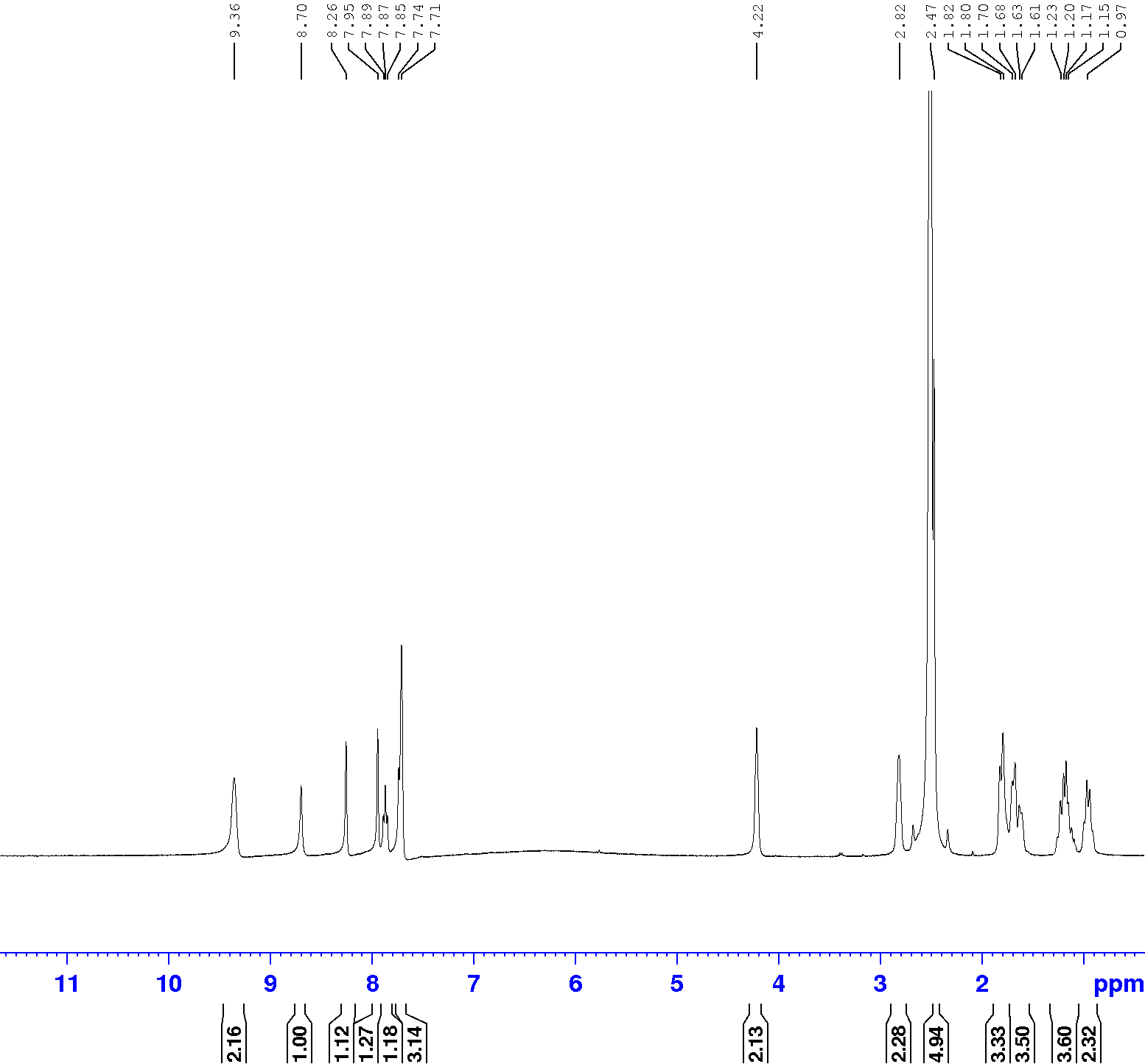

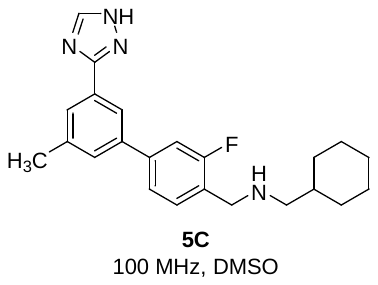

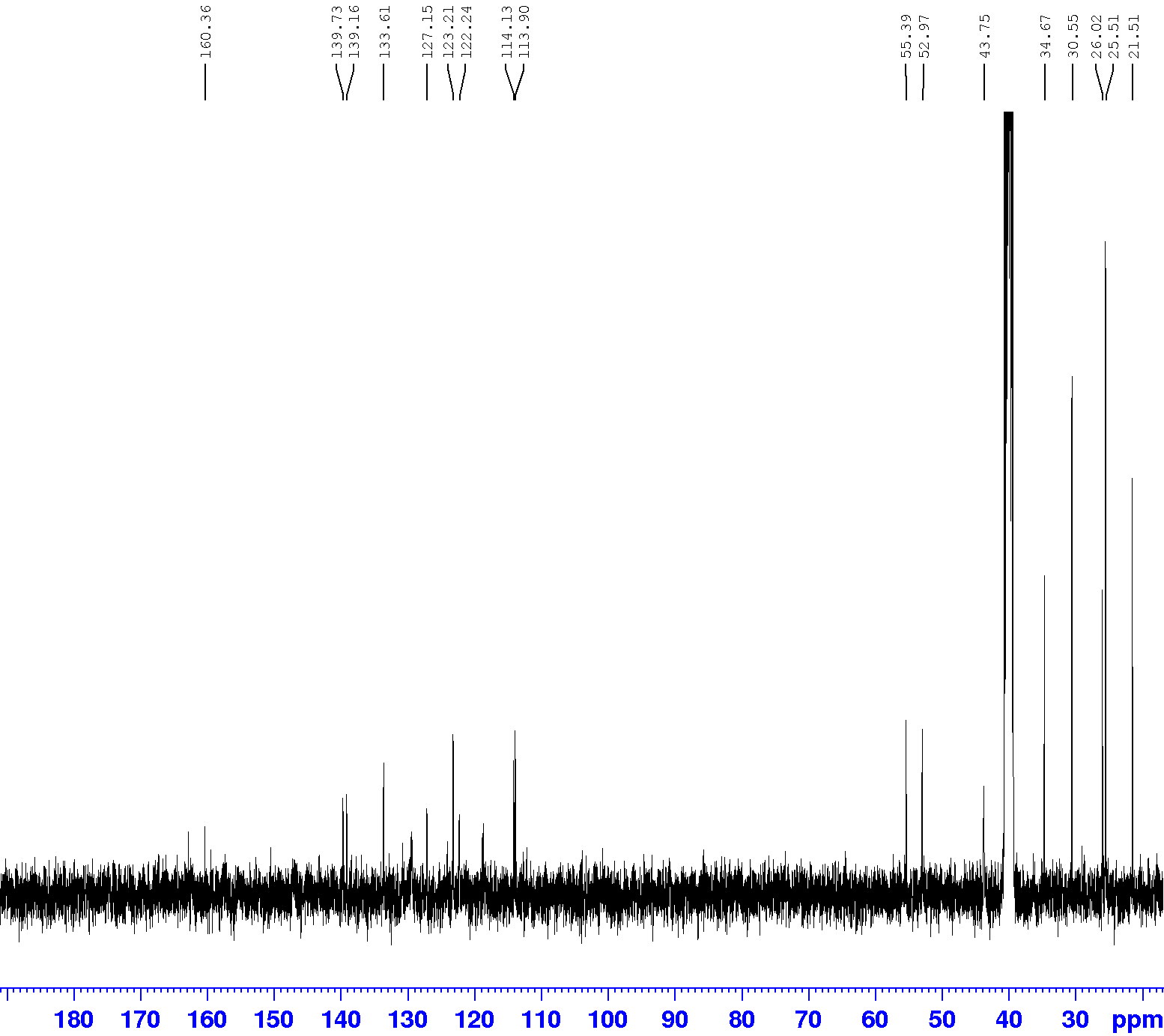

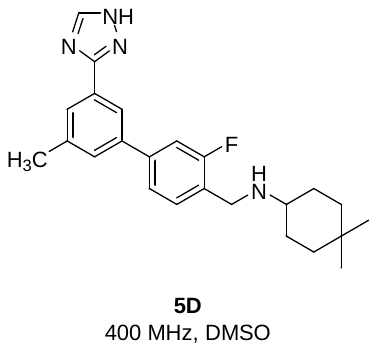

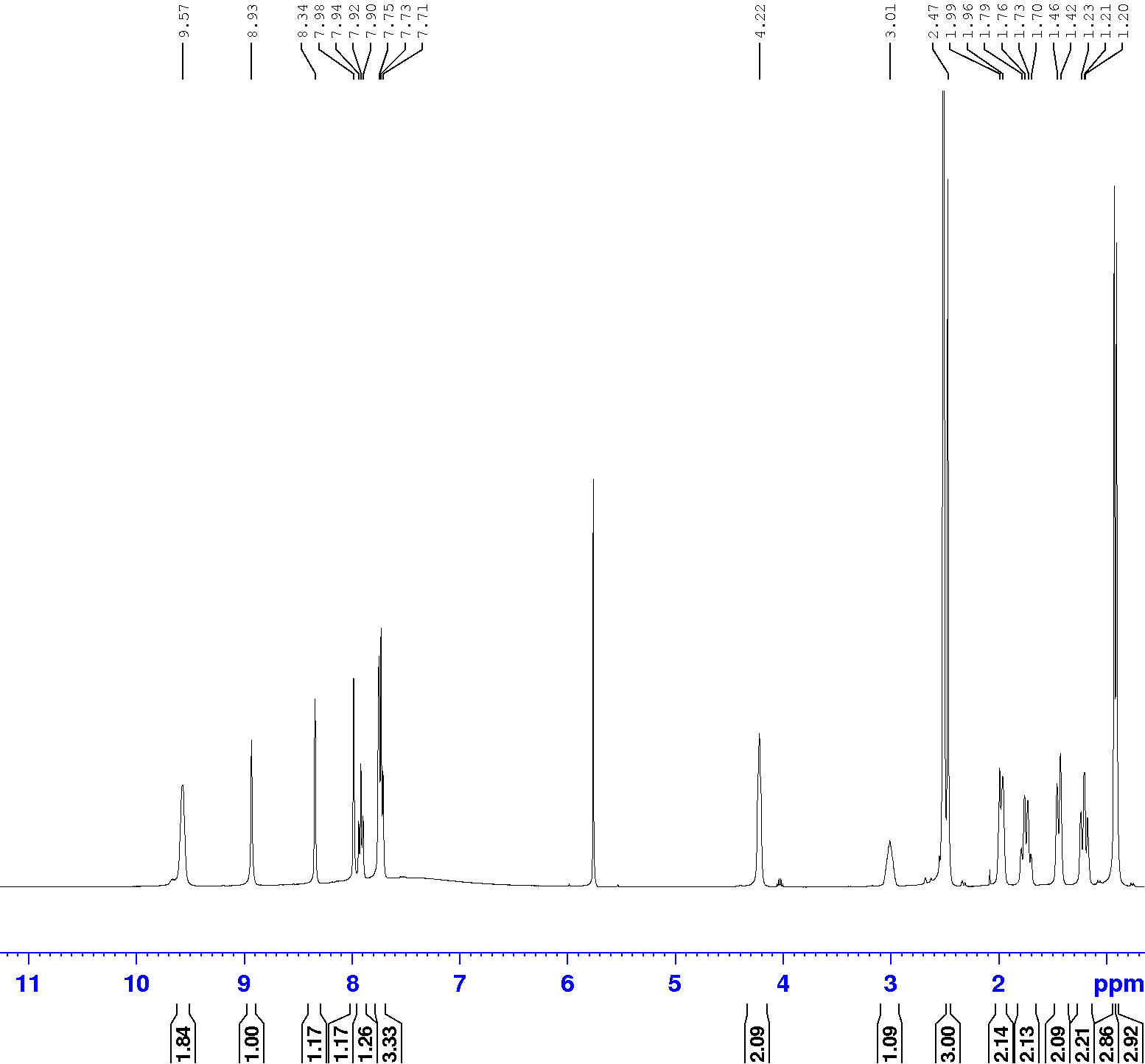

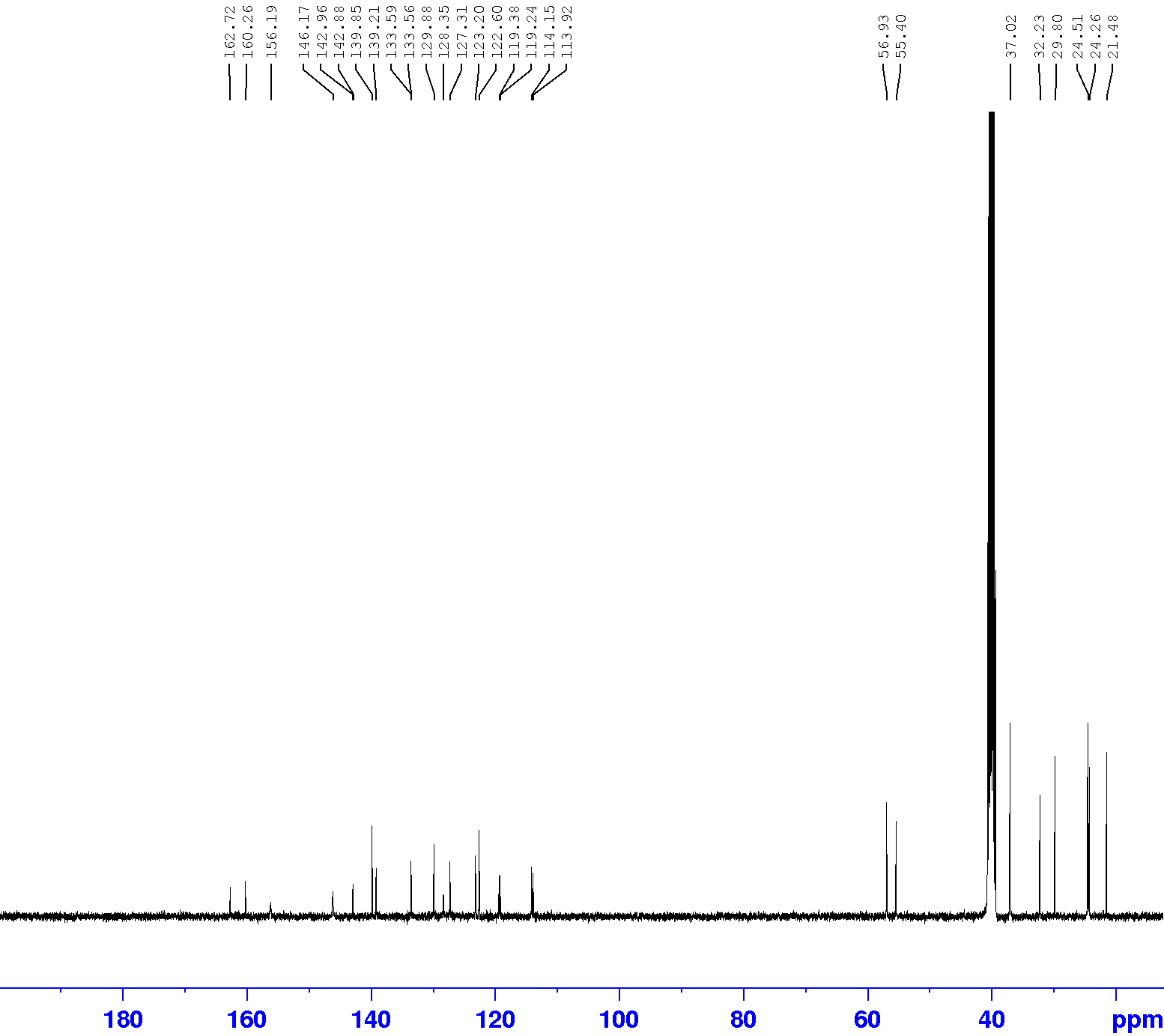

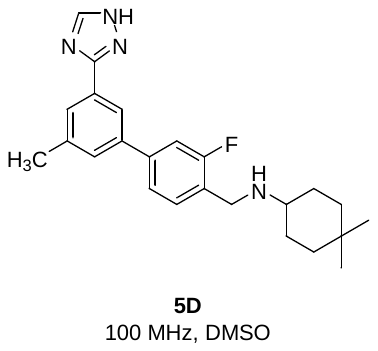

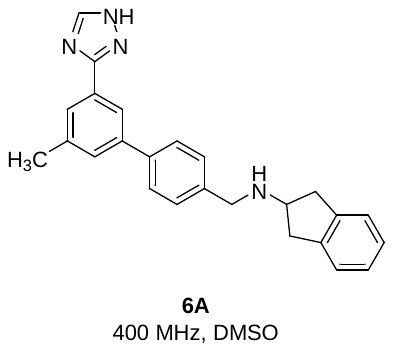

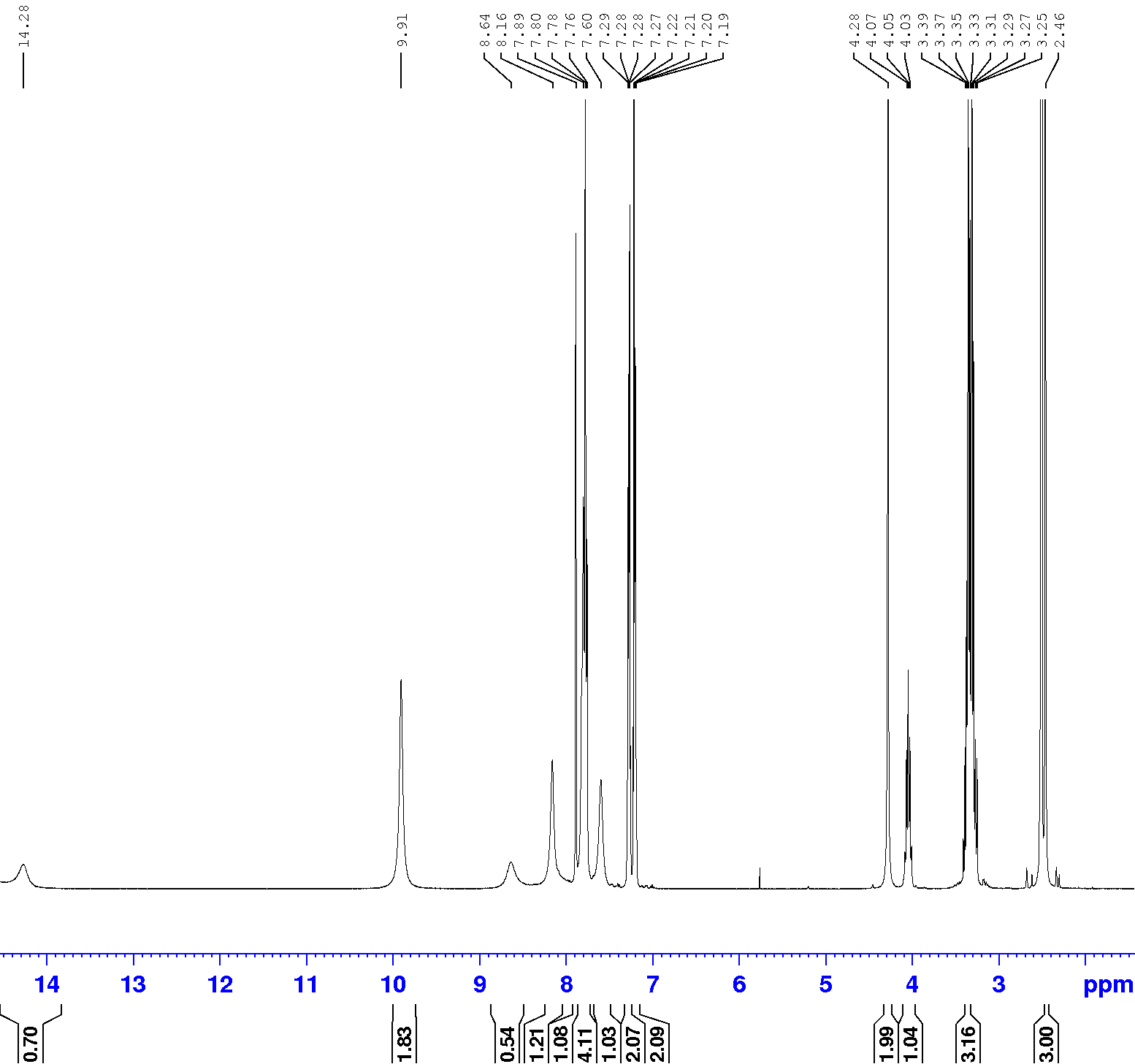

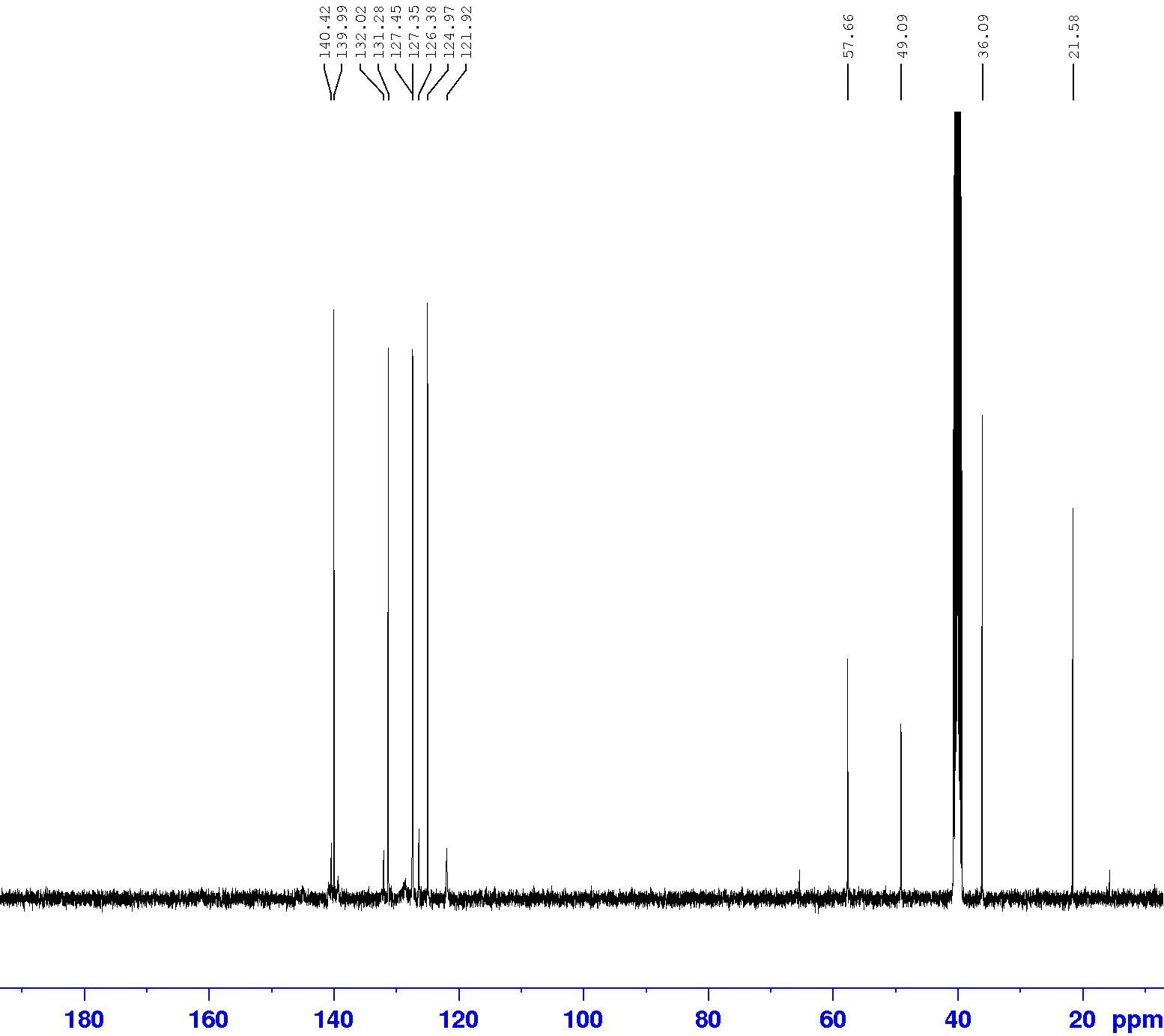

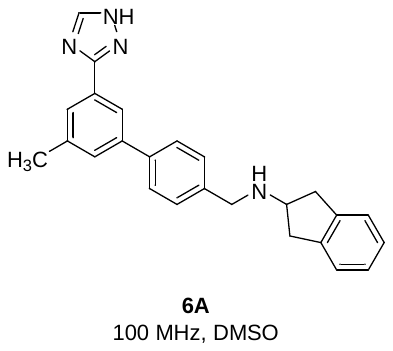

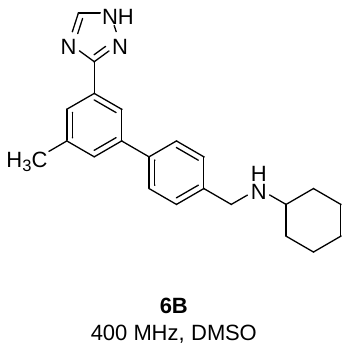

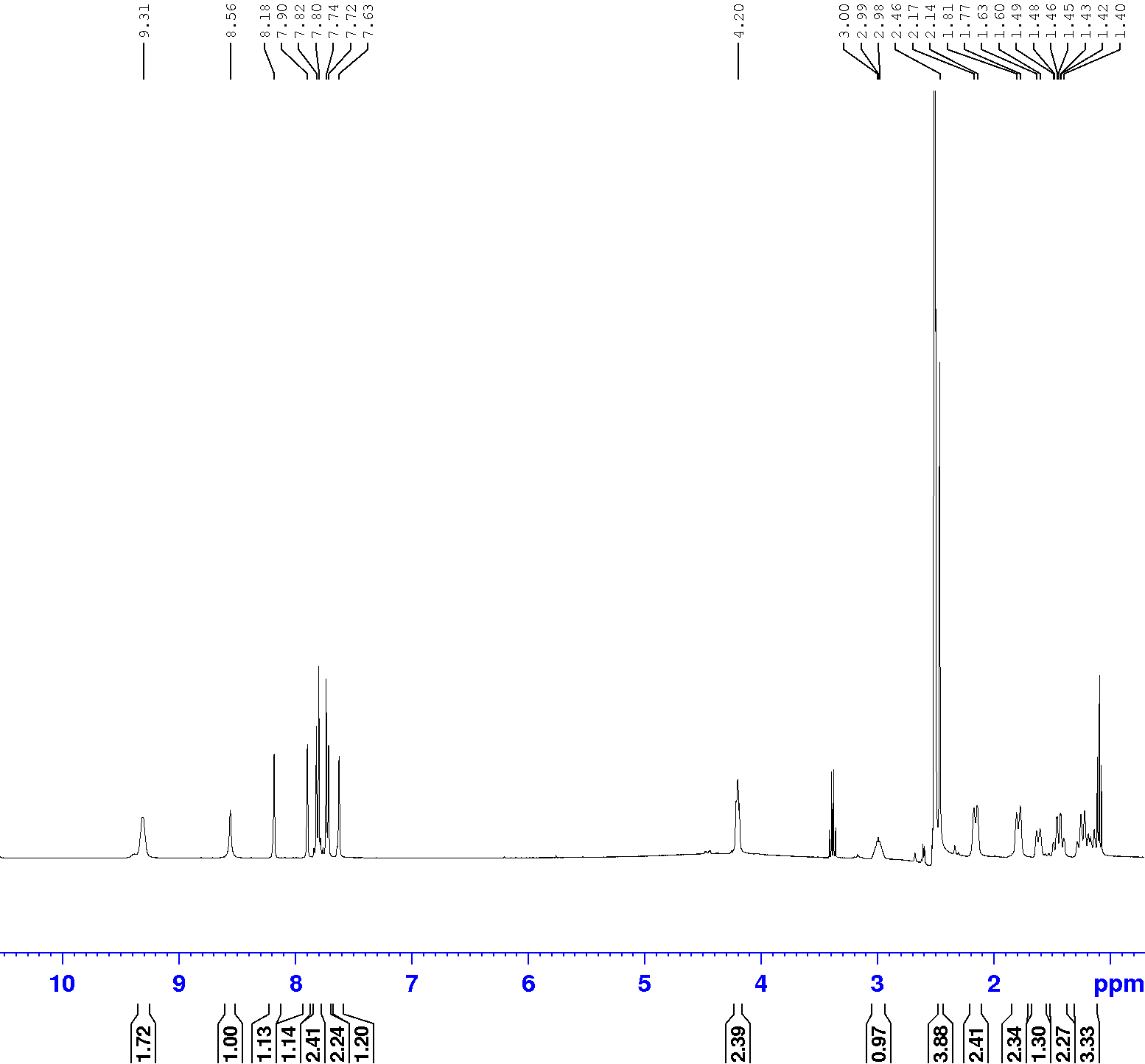

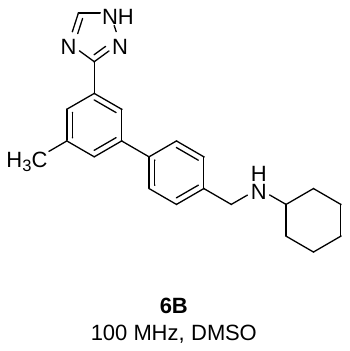

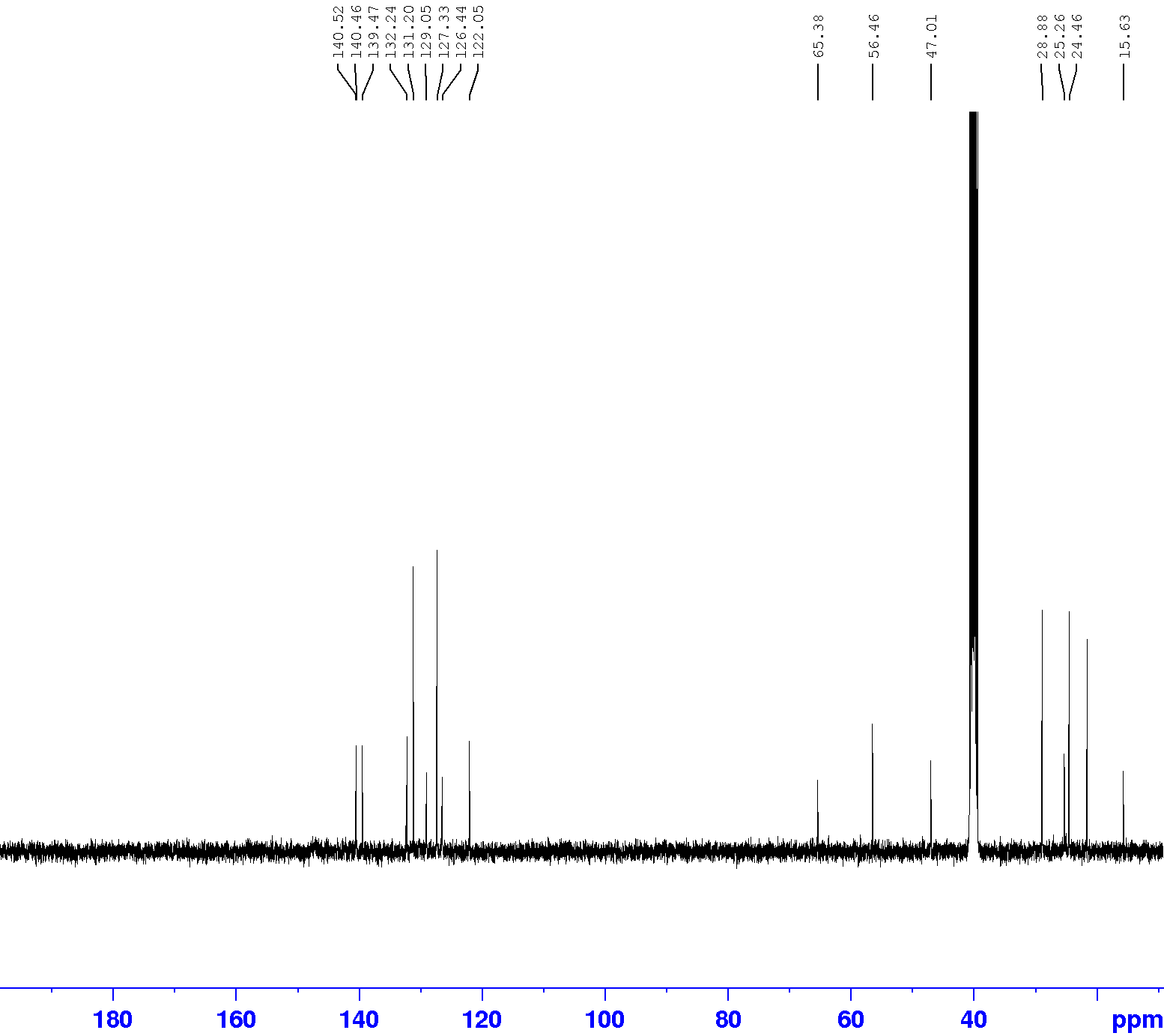

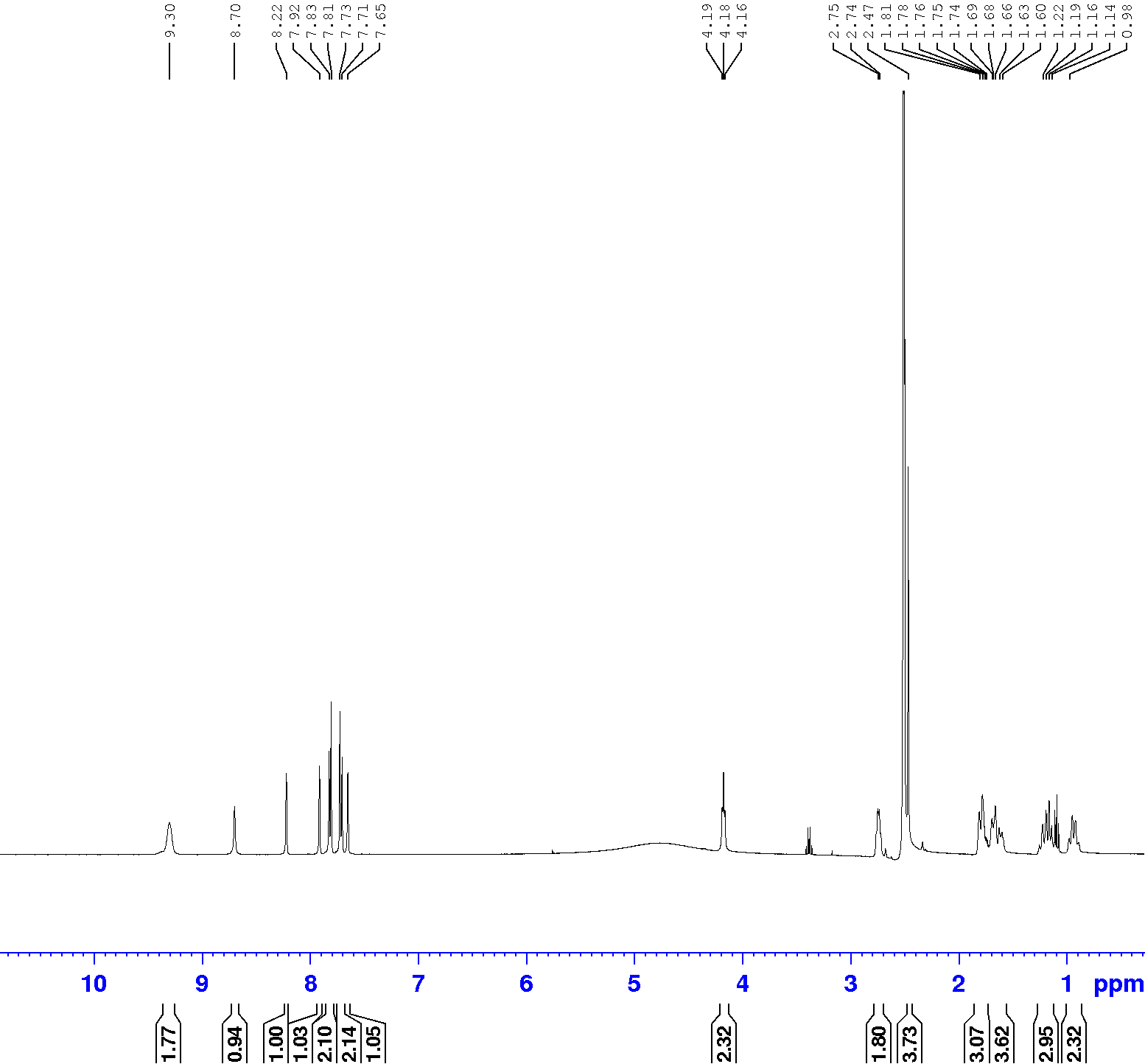

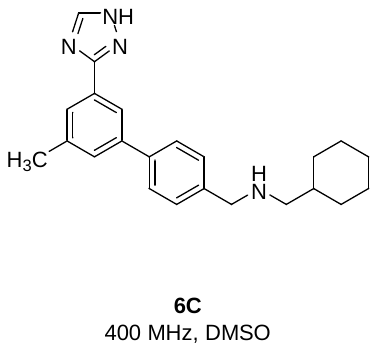

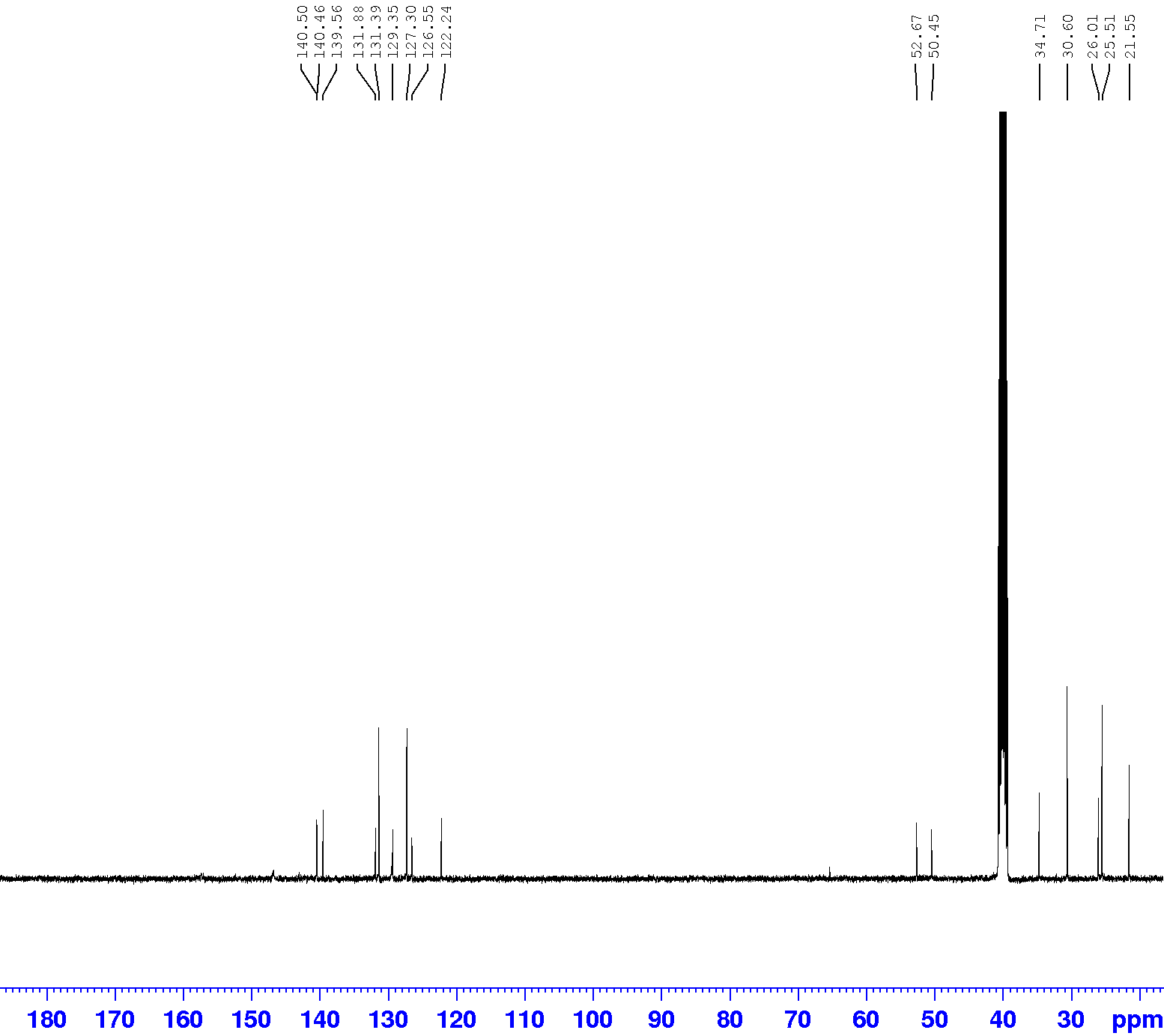

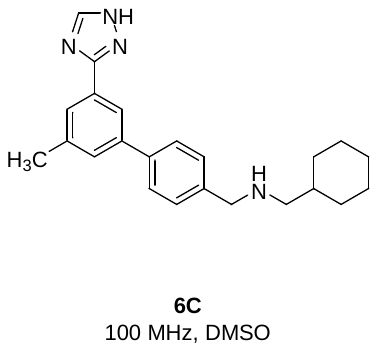

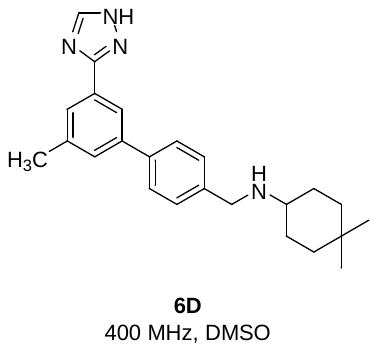

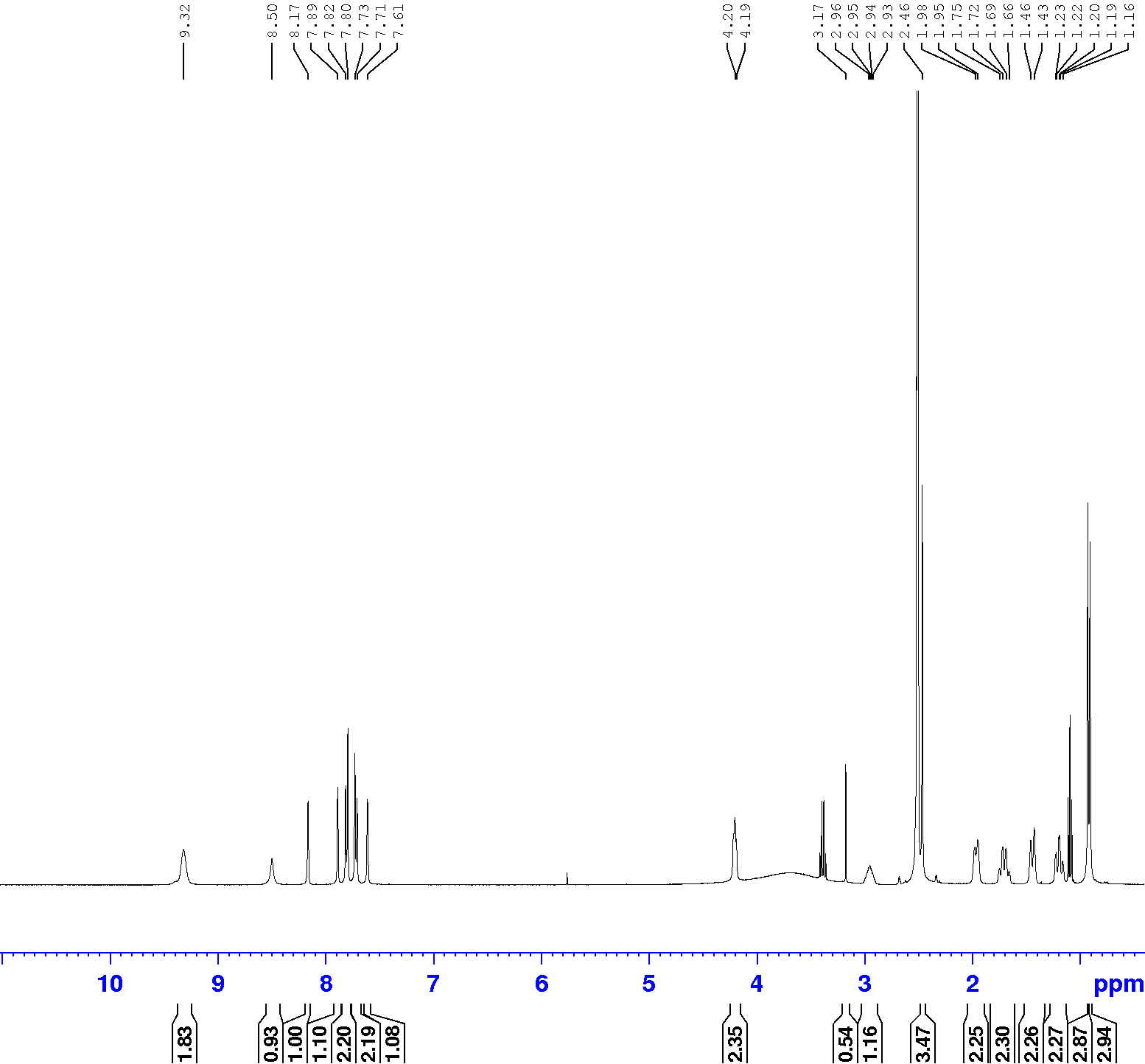

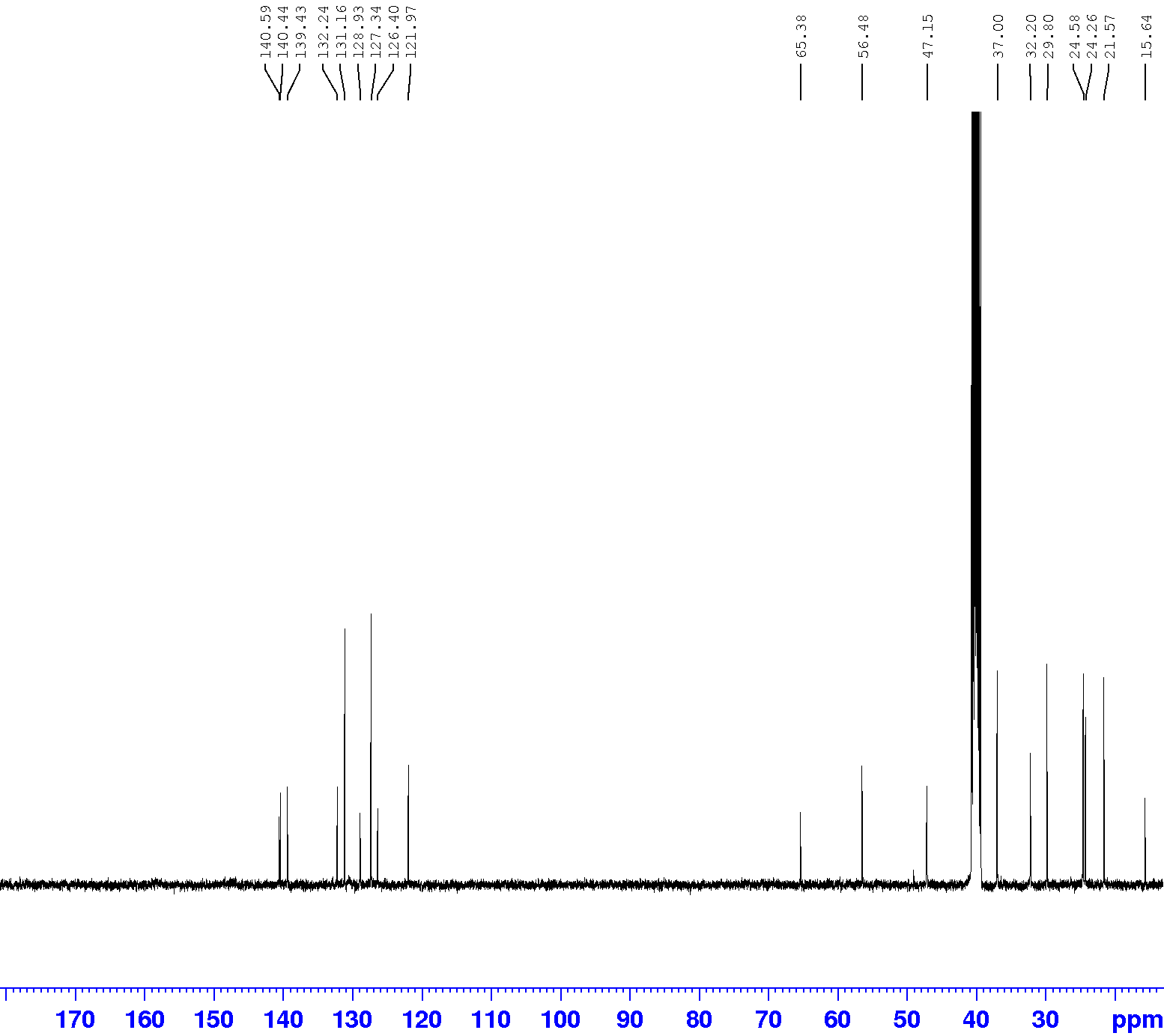

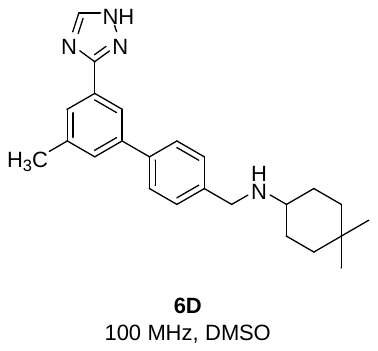
